# *Plasmodium* DDI1 is a potential therapeutic target and important chromatin-associated protein

**DOI:** 10.1101/2021.10.29.466443

**Authors:** Nandita Tanneru, M Angel Nivya, Navin Adhikari, Kanika Saxena, Zeba Rizvi, Renu Sudhakar, Amit Kumar Nagwani, Atul, Faisal Mohammed Abdul Al-Nihmi, Arun Kumar Kota, Puran Singh Sijwali

## Abstract

DDI1 proteins are involved in a variety of cellular processes, including proteasomal degradation of specific proteins. All DDI1 proteins contain a ubiquitin-like (UBL) domain and a retroviral aspartyl protease (RVP) domain. Some DDI1 proteins also contain a ubiquitin-associated (UBA) domain. The three domains confer distinct activities to DDI1 proteins. The presence of RVP domain makes DDI1 a potential target of HIV protease inhibitors, which also block the development of malaria parasites. Hence, we investigated the DDI1 of malaria parasites to identify its roles during parasite development and potential as a therapeutic target. DDI1 proteins of *Plasmodium* and other Apicomplexan parasites share the UBL-RVP domain architecture, and some also contain the UBA domain. *Plasmodium* DDI1 is expressed across all the major life cycle stages and is important for parasite survival, as conditional depletion of DDI1 protein in the mouse malaria parasite *Plasmodium berghei* and the human malaria parasite *Plasmodium falciparum* compromised parasite development. Infection of mice with DDI1 knock-down *P. berghei* was self-limiting and protected the recovered mice from subsequent infection with homologous as well as heterologous parasites, indicating potential of DDI1 knock-down parasites as a whole organism vaccine. *P. falciparum* DDI1 (PfDDI1) is associated with chromatin and DNA-protein crosslinks. PfDDI1-depleted parasites accumulated DNA-protein crosslinks and showed enhanced susceptibility to DNA damaging chemicals, indicating a role of PfDDI1 in removal of DNA-protein crosslinks. Knock-down of PfDDI1 increased susceptibility to the retroviral protease inhibitor lopinavir and antimalarial artemisinin, which suggests that simultaneous inhibition of DDI1 could potentiate antimalarial activity of these drugs. As DDI1 knock-down parasites confer protective immunity and it could be a target of HIV protease inhibitors, *Plasmodium* DDI1 is a potential therapeutic target for malaria control.

## 1. Introduction

The ubiquitin proteasome system (UPS) is a major degradation machinery for clearance of unwanted and misfolded proteins in eukaryotes, which is vital for maintaining cellular homeostasis (Goldberg, 2005; Livneh et al., 2016). Proteasome inhibitors have been shown to block the development of malaria parasites at multiple stages, indicating a critical role of the UPS during parasite development (Czesny et al., 2009; Gantt et al., 1998; Lindenthal et al., 2005). Hence, the proteasome of protozoan parasites, including *Plasmodium*, has emerged as a potential drug target (Khare et al., 2016; Li et al., 2016; Ng et al., 2017; Wyllie et al., 2019).

The UPS includes ubiquitin, enzymes involved in tagging the substrate with ubiquitin and the 26S proteasome wherein ubiquitinated proteins are degraded (Ciechanover, 2009; Hershko and Ciechanover, 1982). Several non-proteasomal ubiquitin-binding proteins also contribute to the proteasome function. One such class of proteins are the shuttle proteins, which include Rad23, Dsk2 and DDI1 (Finley, 2009; Saeki, 2017). These proteins contain a ubiquitin-like (UBL) domain and a ubiquitin-associated (UBA) domain, which mediate their interactions with the proteasome and ubiquitin chains of the ubiquitinated protein, respectively. DDI1 also contains an aspartyl protease domain similar to the retroviral aspartic protease (RVP) of HIV protease (Sirkis et al., 2006).

DDI1 was first discovered as one of the over expressed proteins upon treatment of *Saccharomyces cerevisiae* with DNA damaging agents, hence, it was named as the DNA damage inducible 1 protein (DDI1) (Liu and Xiao, 1997). A later study named it v-SNARE master 1 (VSM1) based on its interaction with Snc2 v-SNARE proteins, and showed that knockout of VSM1 increased protein secretion (Lustgarten and Gerst, 1999). VSM1/DDI1 inhibits assembly of Sso t-SNAREs with Sec9 t-SNARE, which is required for exocytosis (Gabriely et al., 2008; Marash and Gerst, 2003). Deletion analysis demonstrated that both UBL and RVP domains of *S. cerevisiae* DDI1 (ScDDI1) are necessary for inhibition of protein secretion. However, the ScDDI1 C-terminal that contains the Sso t-SNARE-binding region and UBA domain was found to be dispensable for suppression of protein secretion (White et al., 2011a), which indicated that suppression of protein secretion by VSM1/DDI1 is independent of its interaction with Sso t- SNARE. Over expression of ScDDI1 rescued the S-phase checkpoint defect of a temperature sensitive Pds1 mutant, which required the UBA but not UBL domain (Clarke et al., 2001). Another study showed that UBL, catalytically competent RVP and UBA domains of ScDDI1 are required for the rescue of S-phase checkpoint defect (Gabriely et al., 2008). ScDDI1 has been shown to shuttle ubiquitinated Ho endonuclease to the proteasome for degradation, which requires interaction of UBA and UBL domains with ubiquitinated Ho endonuclease and the Rpn1 subunit of 26S proteasome, respectively (Kaplun et al., 2005; Voloshin et al.). Interaction of the ScDDI1 UBL domain with the ubiquitin interaction motif (UIM) of the F-box protein Ufo1 has been shown to be critical for proteasomal degradation of Ufo1 (Ivantsiv et al., 2006). The UBA domain of ScDDI1 homolog in Schizosaccharomyces *pombe*, Mud1, has also been shown to bind Lys48-linked polyubiquitin chains (Trempe et al., 2005). Interestingly, DDI1 itself is a substrate of E3 ligase UBE3A in *Drosophila melanogaster*, which does not target ubiquitinated DDI1 for degradation, and the biological significance of ubiquitination of DDI1 is not known yet (Ramirez et al., 2018). Recent studies have demonstrated important roles of ScDDI1 in removing the replication termination factor RTF2 from stalled replication forks and DNA-topoisomerase complexes (Serbyn et al., 2020; Svoboda et al., 2019), which require catalytically competent ScDDI1, indicating that ScDDI1 functions as a protease to remove DNA- protein crosslinks.

The DDI1 homolog of *Drosophila melanogaster*, Rngo, is essential for oocyte formation (Morawe et al.). As has been shown for ScDDI1, Rngo also dimerizes through the RVP domain, binds ubiquitin via the UBA domain and interacts with the Rpn10 subunit of the proteasome via the UBL domain. The *Homo sapiens* DDI2 (HsDDI2) and *Caenorhabditis elegans* DDI1 (CeDDI1) have been shown to cleave the ER-associated Nrf1/Skn-1A, which results in the release and translocation of the cleaved form to the nucleus wherein it upregulates the proteasome genes (Koizumi et al., 2016; Lehrbach and Ruvkun, 2016). Despite multiple studies indicating requirement of catalytically competent DDI1, a direct demonstration of the protease activity of DDI1 proteins has been elusive until recently. Two recent studies directly demonstrated that HsDDI2 and ScDDI1 cleave ubiquitinated Nrf1 and a peptide substrate corresponding to the Nrf1 cleavage site, respectively (Dirac-Svejstrup et al., 2020; Yip et al., 2020). These two studies also indicated that ScDDI1 and HsDDI2 do not cleave ubiquitin chains, hence, unlikely to function as deubiquitinases.

As described above, multiple studies highlight contributions of UBL, RVP and UBA domains in functions of DDI1 proteins. How these domains crosstalk with each other in the full-length DDI1 is not known due to the unavailability of a full-length DDI1 structure. Structural studies of the RVP domains of ScDDI1, LmDDI1 and human DDI2 revealed that RVP domain exists as a dimer in which aspartyl protease motifs from both the monomers contribute to the formation of active site (Kumar and Suguna, 2018; Sirkis et al., 2006; Sivá et al., 2016; Trempe et al., 2016). The active site of ScDDI1 has identical geometry with that of HIV protease, but has wider substrate binding groove than HIV protease, suggesting that ScDDI1 can accommodate bulkier substrates (Sirkis et al., 2006). Interestingly, in addition to the UBA domain, the UBL domains of ScDDI1 and HsDDI2 also contain a ubiquitin-interaction motif (UIM) that mediates interaction with ubiquitin (Nowicka et al., 2015; Sivá et al., 2016; Trempe et al., 2016).

As a therapeutic target, not much is explored on the role of DDI1 in many economically important protozoan parasites, despite a significant adverse impact of these on human and domestic animal health. The DDI1 proteins of *Leishmania major* and *Toxoplasma gondii* are the only reported representatives. The *Leishmania major* DDI1 (LmDDI1) has been proposed to be the major target of the anti-leishmanial effect of HIV protease inhibitors (White et al., 2011b). Recombinant LmDDI1 has also been shown to degrade peptides and BSA at pH 4-5, which appears to be contrary to the presence of DDI1 proteins in cytosolic pH environment (Perteguer et al., 2013). Knock-out of *Toxoplasma gondii* DDI1 (TgDDI1) has been shown to cause accumulation of ubiquitinated proteins and loss of virulence (Zhang et al., 2020). HIV protease inhibitors have been shown to block the development of malaria parasites at multiple stages (Hobbs et al., 2013; Hobbs et al., 2009; Nsanzabana and Rosenthal, 2011b; Parikh et al., 2005; Skinner-Adams et al., 2004). Furthermore, a study in humans showed that treatment with HIV protease inhibitor-based antiretroviral therapy reduced incidence of malaria by 41% compared to the group treated with non-protease inhibitor-based anti-retroviral therapy, and the lower incidence was attributable to decreased recurrence of malaria (Achan et al., 2012). As DDI1 is important for several cellular processes and the presence of RVP domain makes it a potential target of HIV protease inhibitors, we undertook cellular, genetic and biochemical approaches to investigate the *Plasmodium* DDI1. Our study revealed that *Plasmodium* DDI1 is critical for parasite development, associated with chromatin and contributes to DPC repair, and it could be a target of HIV protease inhibitors.

## 2. Materials and methods

All the biochemicals were from Merck, Sigma-Aldrich or SERVA unless otherwise mentioned. Cell culture reagents were from Gibco and Lonza. DNA modifying enzymes were from New England Biolabs, Thermo Fisher Scientific or TaKaRa. DNA and RNA isolation kits were from Qiagen or MACHEREY-NAGEL. Ni-NTA agarose was from Qiagen or Thermo Fisher Scientific. SuperSignal West Chemiluminescent Substrates were from Pierce. Secondary antibodies were from Pierce or Sigma-Aldrich. ProLong^TM^ gold anti-fade reagent was from Thermo Fischer Scientific. *P. falciparum* and *P. berghei* ANKA strains were obtained from the Malaria Research and Reagent Reference Resource centre (MR4). Basic Parasite Nucleofector^TM^ Kit 2 was from Lonza. Inhibitors and drugs (etoposide, trimethoprim, hydroxyurea, methyl methanesulfonate MMS, HIV protease inhibitors, E64, chloroquine and artemisinin) were from Sigma-Aldrich, Santa Cruz Biotechnology, Biorbyt, Selleckchem or Tocris Bioscience. Human blood was collected from healthy volunteers after taking written consent under medical supervision at the medical dispensary of the institute according to the protocols approved by Institutional Ethics Committee (IEC) of Centre from Cellular and Molecular Biology, Hyderabad, India. All animal experiments were carried out according to the protocols approved by the Institutional Animal Ethics Committees (IAEC) of Centre from Cellular and Molecular Biology, Hyderabad, India. Animals used in this study were maintained at the animal house of Centre from Cellular and Molecular Biology, Hyderabad under standard environmental conditions (22–25°C, 40–70% humidity, and 12:12 hour dark/light photoperiod), and euthanized using CO_2_ at the end of experiment. The studies in this work abide by the Declaration of Helsinki principles.

### 2.1. Sequence analysis of DDI1 proteins

The PfDDI1 amino acid sequence (PF14_0090) was obtained from the *Plasmodium* genome database PlasmoDB by BLAST search using ScDDI1 amino acid sequence (P40087) as a query. The sequences of DDI1 homologs were obtained from the UniProtKB and EuPathDB databases using PfDDI1 and ScDDI1 amino acid sequences as queries. Sequence alignment of DDI1 proteins was performed using the Clustal Omega (https://www.ebi.ac.uk/Tools/msa/clustalo/). Sequences of DDI1 proteins were analyzed for the presence of conserved domains and catalytic site using the Conserved Domain Database (https://www.ncbi.nlm.nih.gov/Structure/cdd/wrpsb.cgi) and Pfam (http://pfam.xfam.org/) softwares. For phylogenetic analysis, sequences were aligned using MUSCLE, alignment was curated with Gblocks, phylogenetic tree was constructed using PhyML and edited with TreeDyn (http://www.phylogeny.fr/simple_phylogeny.cgi). A structure of full-length DDI1 is not available, yet. Hence, we superimposed the AlphaFold structure of PfDDI1 on the reported structures of ScDDI1 UBL (PDB IDs: 2N7E) and RVP (PDB ID: 2I1A) domains using the PyMOL Molecular Graphics System. The DDI1 proteins of representative Apicomplexan parasites were also analyzed for the presence of UBL, RVP and UBA domains by the Swiss Model online software. Homology models of *Cytauxzoon felis* UBA (Cf-UBA), *Theilaria annulata* UBA (Ta-UBA) and *Gregarina niphandrodes* UBL (Gn-UBL) were generated using the closest related templates structures (PDB: 1Z96 for *S. pombe* UBA (39.47% sequence identity with Cf-UBA and 26.32% Ta-UBA), 2N7E for ScDDI1 UBL (15.94% sequence identity with Gn-UBL)) by Swiss Model online software. The modelled structures were superimposed on the template structures and RMSD values were calculated using the PyMOL Molecular Graphics System, version 1.3, Schrodinger, LLC.

### 2.2. Parasite culture

*P. falciparum* 3D7 and D10 strains were cultured in RPMI-1640 medium (supplemented with 2.0 g/l glucose, 0.5% AlbuMAX II, 41.1 mg/l hypoxanthine, 300 mg/l glutamine, 25 mg/l gentamicin and RBCs at 2% hematocrit) at 37°C under a gas mixture (5% CO_2_ + 5% O_2_ + 90% N_2_). Giemsa stained smears of the culture were regularly prepared and observed under the 100× magnification of a light microscope to monitor parasite development. Synchrony was maintained by treating the cultures at ring stage with 5% sorbitol (Lambros and Vanderberg, 1979). The cultures were harvested at 10-15% parasitemia whenever required and parasites were purified from infected-RBCs by saponin (0.05% in PBS) lysis method. For *P. berghei* ANKA, a frozen stock was injected intra-peritoneally into a 4-6 weeks old naïve female BALB/c mice. The infection and was monitored by observing Giemsa stained smears of the tail-snip blood on every other day of post-infection. Mice were sacrificed at 6-10% parasitemia, the blood was collected by cardiac puncture in Alsever’s solution (2.05 % Glucose, 0.8% sodium citrate, 0.055% citric acid, and 0.42% sodium chloride) and the cell pellet was treated with 0.1% saponin to isolate parasites. The pellets of purified parasites were used immediately or stored at -80°C till further use. *P. falciparum* gametocytes were obtained by inducing gametocytogenesis with spent medium as has been described previously (Fivelman et al., 2007; Singhal et al.). Gametocyte development was monitored by observing Giemsa stained smears of the culture on daily basis and samples at different gametocyte stages were collected to determine localization of PfDDI1. Genomic DNA was isolated from trophozoite/schizont stage parasites using the Puregene Blood Core Kit B as instructed by the manufacturer.

### 2.3. Production of recombinant proteins and generation of anti-PfDDI antiserum

The complete PfDDI1 coding region was expressed in *E. coli* as an N-terminal His-tagged protein _His_PfDDI1, and purified by Ni-NTA affinity chromatography. A codon optimized synthetic version of PfDDI1 gene with C-terminal Myc-tag (synPfDDI1_Myc_) was expressed in *E. coli* as a C-terminal His-tag (PfDDI1_Myc/His_), and purified by Ni-NTA chromatography. Wild type *P. berghei* DDI1 with C-terminal Myc-tag was expressed in *E. coli* as an N-terminal His-tag (wPbDDI_Myc_), and purified by Ni-NTA affinity chromatography. A catalytic mutant (D268A) of PbDDI1 with C-terminal Myc-tag was generated by overlap PCR, expressed as an expressed in *E. coli* as an N-terminal His-tag (mPbDDI_Myc_), and purified by Ni-NTA affinity chromatography. See supplementary material for details.

Three months old female Wistar rats were immunized intraperitoneally with emulsion of recombinant _His_PfDDI1 (200 µg/immunization) in Freund’s complete (day 0) or incomplete adjuvant (day 21, 42, 84 and 105). Blood was collected (day -1, 14, 35, 70, 91 and 125). The day-125 antiserum was adsorbed with *E. coli* proteins to eliminate the antibodies cross-reactive with bacterial proteins as has been described earlier (Navale et al.). The adsorbed PfDDI1antiserum was supplemented with 0.01% sodium azide and glycerol (50% final) for storage at -80°C, and used to determine expression and localization of PfDDI1 or PbDDI1 in various parasite stages.

### 2.4. Western blotting and immunofluorescence assay

A synchronized culture of *P. falciparum* 3D7 (∼10% parasitemia) was harvested at ring, early trophozoite, mid-trophozoite and late trophozoite/schizont stages. Parasites were purified by saponin lysis, resuspended in 4x pellet volume of SDS-PAGE sample buffer, equal amounts of parasite lysates (corresponding to approximately 1×10^8^ parasites/lane) and uninfected RBC lysate were resolved on 12% SDS-PAGE and the proteins were transferred onto the PVDF membrane. The membrane was sequentially incubated at room temperature with blocking buffer (5% non-fat milk in TBST) for 1 hour, rat PfDDI1 antiserum (at 1/4000 dilution in blocking buffer) for 1 hour, blocking buffer for washing, and HRP-conjugated goat anti-rat IgG (at 1/10,000 dilution in blocking buffer). The membrane was washed with blocking buffer followed by with TBST, incubated with the Supersignal^TM^ west chemiluminescent substrate, and the signal was captured using a ChemiDoc Imaging System. PbDDI1 expression in *P. berghei* erythrocytic stage parasites was checked by western blotting using rat PfDDI1 antiserum as has been described for *P. falciparum*. The membranes were stripped and reprobed with mouse anti-β actin- HRP conjugated antibodies for detection of β actin as a loading control.

For localization of PfDDI1, different developmental stages of *P. falciparum* erythrocytic and gametocyte cultures were collected, the cells were washed with PBS, immobilized on poly-L coated slides, fixed (4% paraformaldehyde/0.002% glutaraldehyde in PBS) for 45 minutes, and permeabilized (0.01% Triton-X 100). The slides were sequentially incubated at room temperature in blocking buffer (3% BSA in PBS), rat PfDDI1 antiserum (at 1/500 dilution in blocking buffer) or mouse anti-Myc antibodies (1/200 dilution in blocking buffer), and appropriate secondary antibodies (Alexa fluor-488 conjugated donkey anti-rat IgG, Alexa fluor- 594 conjugated donkey anti-rat IgG, Alexa fluor-488 conjugated donkey anti-mouse IgG (all at 1/2000 dilution in blocking buffer, with DAPI at 10 µg/ml)). The slides were air dried, mounted with ProLong Gold antifade, and images were captured using Zeiss AxioImager Z with Apotome or Zeiss Axioimager 2 microscope under the 100× objective. Images were processed using Zeiss Axiovision and Adobe Photoshop softwares. For Z-section images, wild type *P. falciparum* trophozoite stage parasites were fixed and probed using rat PfDDI1 antiserum followed by Alexa fluor-488 conjugated donkey anti-rat IgG as described for IFA, images were captured using the Leica TCS SP8 confocal laser scanning microscope, and edited using the Leica Application Suite software. For PbDDI1 localization in *P. berghei* erythrocytic trophozoites, 10 μl blood was collected from the tail-snip of a *P. berghei* ANKA-infected mouse and the cells were processed for IFA using rat PfDDI1 antiserum as has been described for PfDDI1 localization in *P. falciparum*. For PbDDI localization in sporozoite and liver stages, female Anopheles mosquitoes were fed on *P. berghei* ANKA-infected BALB/c mice and dissected on day 18 for isolation of salivary gland sporozoites. The sporozoites were spotted on a glass slide and air dried. 1×10^4^ sporozoites were added on the HepG2 monolayer cultured in DMEM with 10% FBS in labtek chamber slides and grown for 36 hours. The sporozoite and liver stage samples were processed for IFA using rat PfDDI1 antiserum for PbDDI1, mouse anti-circumsporozoite protein (CSP) antibodies for the sporozoites, and rabbit anti-upregulated in infective sporozoites-4 (UIS-4) antibodies for the liver stage as has been described earlier (Al-Nihmi et al., 2017). The slides were incubated with appropriate secondary antibodies (Alexa Fluor-594 conjugated donkey anti-rat IgG, Alexa flour-488 conjugated rabbit anti-mouse IgG for CSP and Alexa flour-488 conjugated mouse anti-rabbit IgG for UIS-8), observed under the 40× objective, images were captured and processed as has been described for PfDDI1 localization in *P. falciparum*.

### 2.5. Generation of PbKD parasites

Transfection plasmids to knock-out (HB-pbDDIKO) and knock-down (HB-pbDDIKD) the PbDDI1 gene were constructed as described in the supplementary material. Transfection of *P. berghei* with HB-pbDDIKO and HB-pbDDIKD plasmids would replace the PbDDI1 coding sequence with GFP and PfDDI_Myc_/cDD_HA_ coding regions, respectively. A frozen stock of *P. berghei* ANKA was injected intraperitoneally into a naïve BALB/c mouse. Infection was monitored and blood was collected as has been described in the parasite culture section. The cells were washed with FBS-RPMI medium (RPMI-1640, 20% FBS, 2 g/l Glucose, 2 g/l, Sodium bicarbonate, 41.1 mg/ml Hypoxanthine, pH-7.5), resuspended in FBS-RPMI medium at 2% hematocrit, gassed (5% CO_2_, 5% O_2_, 90% N_2_), and incubated at 37°C for 12-14 hours with shaking of 55 rpm. Parasites were purified on a 65% Nycodenz gradient cushion when the majority of the parasites reached schizont stage. The cells were washed with FBS-RPMI medium and aliquoted, each with 15-20 µl of packed cell volume. An aliquot was mixed with 100 µl of Amaxa T cell nucleofactor and 2.5-5 µg of pbDDIKO or pbDDIKD transfection constructs, pulsed using the Amaxa Nucleofector (U303 program), and injected intravenously into a naive mouse. One day after the transfection, the infected mouse was given drinking water with 70 µg/ml pyrimethamine (for knock-out) or 70 µg/ml pyrimethamine + 40 µg/ml trimethoprim (for knock-down) for 7-10 days to selected resistant parasites. Blood Smears were checked for the presence of parasites on day 7 post-transfection and then every other day. The genomic DNA and lysates of resistant parasites were checked for integration and expression of desired protein by PCR and western blotting, respectively. Cloned lines of *P. berghei* knock-down parasites (PbKD) were obtained by infecting 15 naïve BALB/c mice intravenously with 200 µl of diluted parasite suspension (0.5 parasite/mouse). Mice were given pyrimethamine + trimethoprim in drinking water and infection was monitored by making blood smears. Blood was collected from infected mice as has been described in the parasite culture section. Part of the blood was used for making frozen stocks for later use, and the remaining blood was processed for isolation of genomic DNA and preparation of parasite lysate to check for desired integration and expression of the desired protein by PCR and western blotting, respectively.

The genomic DNAs of wild type parasites and two PbKD clonal lines were assessed for the presence of recombinant locus by PCR with primers specific for 5’ specific integration (CON5’-F/Pvac-R), 3’ integration (Hrp2-F/CON3’-R), knocked-in gene (Pfspec-F/DDimyc Rep- R), wild type locus (Pbspec-F/PbDdiexpRm) and a control target (α6FL2–F/α6FL2-R). The PCR products were separated by agarose gel electrophoresis. The erythrocytic stage parasite lysates of wild type *P. berghei* ANKA and two PbKD clones were processed for western blotting using appropriate antibodies (mouse anti-Myc, mouse anti-HA and mouse anti-β actin for loading control), followed by HRP-conjugated goat anti-mouse IgG as described in the western blotting section.

### 2.6. Evaluation of PbKD parasites

The PbKD parasites were assessed for the effect of knock-down on PfDDI_Myc_/cDD_HA_ protein level, erythrocytic growth and virulence. For the effect on PfDDI_Myc_/cDD_HA_ protein level, a 2-3 months old naïve BALB/c mouse was infected intraperitoneally with a fresh stock of PbKD parasites, maintained under 70 µg/ml pyrimethamine + 40 µg/ml trimethoprim in drinking water, and blood was collected at 8-10% parasitemia. The blood was washed with FBS-RPMI medium, resuspended in FBS-RPMI medium at 2% haematocrit without or with 50 µM trimethoprim, gassed, and incubated for 12 hours at 37°C. Parasites were purified by saponin lysis and processed for western blotting using mouse anti-HA and mouse anti-β actin (as a loading control), followed by HRP-conjugated goat anti-mouse IgG as described in the western blotting section.

For the effect of growth and virulence, 2-3 months old female BALB/c mice were infected intraperitoneally (2x10^5^ parasites/mouse) with wild type *P. berghei* ANKA (5 mice) or PbKD parasites (10 mice). The PbKD-infected mice were divided into 2 groups of 5 mice each; one group was given 70 µg/ml pyrimethamine in drinking water (-TMP) and another group was given 70 µg/ml pyrimethamine + 40 µg/ml trimethoprim in drinking water (+TMP). Blood was collected from each mouse one day before the infection (pre-immune sera) and during later time points (days 14 and 28 of post-infection). Each mouse was monitored for infection by making Giemsa stained blood smears, parasitemia was determined by counting a minimum of 1000 cells per smear, and the data was plotted against days post-infection using the GraphPad Prism software. To determine the effect of delayed knock-down, nine female BALB/c mice were infected intraperitoneally (2x10^5^ parasites/mouse) with PbKD parasites and given 70 µg/ml pyrimethamine + 40 µg/ml trimethoprim in drinking water. Trimethoprim was withdrawn from the drinking water of 6 mice at 5% parasitemia (-TMP), while the remaining 3 mice were given 70 µg/ml pyrimethamine + 40 µg/ml trimethoprim in drinking water (+TMP). Infection was monitored regularly by making Giemsa stained blood smears, parasitemia was determined by counting a minimum of 1000 cells per smear, and the data was plotted against days post-infection using the GraphPad Prism software.

### 2.7. Challenge of recovered mice

The PbKD parasite-infected BALB/c that were never given trimethoprim (-TMP) or trimethoprim was withdrawn from the drinking water at 5% parasitemia (delayed –TMP) cleared infection. These were called recovered mice, which were monitored for almost 3 months and then challenged with wild type *P. berghei* ANKA (2x10^5^ parasites/mouse, intraperitoneally). A group of age-matched naïve mice was also challenged similarly. Infection was monitored regularly by observing Giemsa stained blood smears, the parasitemia was determined by counting at least 1000 cells, and plotted against days post-infection using the GraphPad Prism software. 25-50 μl blood was collected from each mouse one day before the challenge (pre-challenge sera) and assessed for reactivity with parasite proteins. A group of recovered mice (4 mice) and age-matched naïve mice (5 mice) were infected with *P. yoelii* 17XNL strain (2 x 10^5^ parasites/mouse, intra-peritoneally). Infection was monitored and data was analyzed as described above for challenge with *P. berghei* ANKA.

### 2.8. Immunization of mice with recombinant PbDDI1

2-3 months old 5 female BALB/c mice were immunized intraperitoneally with emulsion of recombinant mPbDDI_Myc_ (25 µg/immunization) in Freund’s complete (day 0) or incomplete (days 15, 45, 75 and 105) adjuvant. A group of 5 mice was kept as a naive control and another group of 5 mice was immunized with adjuvant only. Blood was collected from each mouse for obtaining preimmune sera (day -1) and immune sera (day 30, 60, 90 and 120). The sera were assessed for reactivity against recombinant mPbDDI_Myc_ by enzyme-linked immunosorbent assay (ELISA). The day 120 immune sera were pooled for each group, and assessed for the titer of anti-mPbDDI_Myc_ antibodies by ELISA. Three months after the final serum collection, the immunized, naïve control and adjuvant control mice were infected intraperitoneally with wild type *P. berghei* ANKA (2 x 10^5^ parasites/mouse), infection was monitored and data was analyzed as described above for recovered mice.

### 2.9. Generation of DDI1 knock-down P. falciparum parasites

Transfection plasmid (HF-pfDDIKD) to replace wild type PfDDI1 coding sequence with PfDDI1_Myc_/cDD_HA_ coding region was constructed as described in the supplementary material. Ring stage *P. falciparum* D10 parasites were transfected with HF-pfDDIKD plasmid by electroporation as has been described earlier (Fidock and Wellems, 1997; Govindarajalu et al., 2019). Briefly, a frozen stock of *P. falciparum* D10 was cultured to obtain ∼10% early ring stage parasites. The culture was harvested, ∼100 μl packed cell volume (PCV) was washed with cytomix (120 mM KCl, 0.15 mM CaCl_2_, 2 mM EGTA, 5 mM MgCl_2_, 10 mM K_2_HPO_4_/KH_2_PO_4_, 25 mM HEPES, pH 7.6) and mixed with 320 μl of cytomix containing 100 μg HF-pfDDIKD plasmid DNA. The suspension was transferred to a chilled 0.2 cm electroporation cuvette and pulsed (950 μF, 310 mV, and infinite resistance) using the Bio-Rad gene pulser Xcell™. The transfected parasites were maintained and selected with 0.5 nM WR99210 till the emergence of resistant parasites. Trimethoprim was added to the culture at 10 μM when WR99210 resistant parasites reached to 5% parasitemia. The culture was harvested for isolation of genomic DNA at 3-4 weeks interval, which was used to check for plasmid integration into the target site by PCR. Primer sets specific for 5’-integration (DDi-con5U/cDD-R), 3’-integration (HRP2seq-F/DDi3’int- R), wild type locus (DDi-con5U/ DDi3’int-R) and a positive control target (PfVMP1- Fep/PfVMP1-R) were used to determine integration at the target site. Once plasmid integration at the target site was observed, parasites were cloned by dilution cloning (Sijwali et al., 2006), and the clonal lines were again evaluated for plasmid DNA integration by PCR and expression of PfDDI_Myc_/cDD_HA_ protein by western blotting. For western blotting, erythrocytic stage lysates of wild type and PfDDI1 knock-down (PfKD) parasites were processed using rabbit anti-HA (at 1/1000 dilution in blocking buffer) and anti-β actin antibodies as a loading control, followed by appropriate secondary antibodies (HRP-conjugated goat anti-rabbit IgG and HRP-conjugated goat anti-mouse IgG, at 1/1000 dilution in blocking buffer) as has been described in the western blotting section.

### 2.10. The effect of PfDDI1 knock-down on parasite growth

The PfKD parasites were cultured in the presence of 10 μM trimethoprim and synchronized as has been described in the cell culture section. A 50 ml synchronized culture with ∼10% rings was washed with RPMI1640-albumax medium, resuspended in 50 ml RPMI 1640-albumax medium, and divided into two 25 ml portions. One portion was supplemented with trimethoprim and the other portion was supplemented with DMSO (0.02%). Both the portions were gassed and incubated at 37°C. After 30 hours of incubation, 20 ml of the culture was harvested from each portion (30 hour time point) and the remaining culture was expanded to 20 ml with additional medium, fresh RBCs and trimethoprim or DMSO. The cultures were gassed, incubated for 30 hours and harvested (60 hour time point). The harvested cultures were processed for isolation of parasites, which were assessed for the level of PfDDI_Myc_/cDD_HA_ protein by western blotting using mouse anti-Myc antibodies or rat PfDDI1 antiserum, followed by appropriate HRP-conjugated secondary antibodies as described in the western blotting section. The membranes were stripped and reprobed with mouse anti-β actin-HRP conjugated antibodies to detect β actin as a loading control as described in the western blotting section. For quantitation of PfDDI_Myc_/cDD_HA_ protein level in western blots, the signal intensity of full-length DDI_Myc_/cDD_HA_ and β-actin bands were measured using the ImageJ software. The signal intensity of full-length DDI_Myc_/cDD_HA_ was normalised with that of the corresponding β-actin band, and plotted using the GraphPad Prism software.

For assessment of PfDDI knock-down on erythrocytic growth, synchronized PfKD parasites were cultured for four consecutive cycles (with or without 10 μM trimethoprim), beginning with 3% parasitemia. The cultures were diluted at the end of each growth cycle and fresh medium and RBCs maintain 3-5% parasitemia. At the end of each growth cycle, the cells were fixed with 2% formaldehyde (in PBS), stained with YOYO-1 (PBS with 0.1% Triton X-100 and 10 nM YOYO-1), and the number of infected cells was determined by counting 10,000 cells using the BD Fortessa FACS Analyzer (λex 491 nm and λem 509 nm). The parasitemia was normalized to initial parasitemia, and plotted as fold multiplication over growth cycles using the GraphPad Prism software.

To assess the effect of DDI1 knock-down on ubiquitinated proteome of PfKD parasites, a synchronized 80 ml ring stage PfKD culture (∼15% parasitemia) was divided into two halves. One half was grown with trimethoprim and the other without trimethoprim. 30 ml of each culture was harvested at 30 hour stage (30 hour time point) and parasites were purified. Each of the remaining 10 ml cultures was expanded to 30 ml with fresh medium and RBCs, grown under the same trimethroprim condition for another 48 hours, and parasites were purified (78 hour time point). The parasite pellets were resuspended in 10× pellet volume of the lysis buffer (25 mM Tris-Cl, 10 mM KCl, 0.1 mM EDTA, 0.1 mM EGTA, 1 mM DTT, pH 7.4, 0.25% NP-40 and protease inhibitor cocktail) and incubated on ice for 30 minutes. The lysate was passed twice through a 25G needle and centrifuged at 2300g for 20 minutes at 4°C. The supernatants were processed for western blotting using mouse anti-ubiquitin (P4D1) antibodies, followed by HRP-conjugated goat anti-mouse IgG antibodies as has been described in the western blotting section. The membrane was stripped and reprobed with mouse anti-β actin-HRP conjugated antibodies for detection of β actin as a loading control as has been described in the western blotting section. For quantitation of ubiquitinated protein levels in western blots, the signal intensity of ubiquitinated protein bands were measured using the ImageJ software, and plotted using the GraphPad Prism software.

### 2.11. Effect of inhibitors on PfKD parasites

The wild type *P. falciparum* D10 and PfKD parasites were compared for susceptibility to HIV protease inhibitors (lopinavir, nelfinavir and saquinavir), artemisinin, E64 and epoxomicin. For PfKD parasites, trimethoprim was maintained at 10 μM during the assay cycle (+T) or excluded during the assay cycle (-T1) or excluded in the previous cycle and during the assay cycle (-T2). For each compound, the stock was serially diluted 2-fold in 50 μl of RPMI1640-albumax medium across the wells of a 96-well tissue-culture plate. Control wells contained DMSO (0.5%) or chloroquine (500 nM). 50 μl of parasite suspension (1% ring-infected erythrocytes at 4% haematocrit) was added to each well, the plate was incubated in a modular incubator chamber (Billups-Rothenberg, Inc.) filled with the gas mixture at 37°C for 48-50 hours. At the end of incubation, the cells were fixed with 2% formaldehyde (in PBS), stained with YOYO-1 (PBS with 0.1% Triton X-100 and 10 nM YOYO-1), and the number of infected cells was determined by counting 10,000 cells using the BD Fortessa FACS Analyzer (λex 491 nm and λem 509 nm). The parasitemia of chloroquine control was subtracted from those of DMSO control and test samples to adjust for background. The adjusted parasitemias of test samples were normalized as % of the DMSO control and plotted against the concentrations of compounds to determine IC_50_ concentrations using the GraphPad Prism software.

### 2.12. Production of recombinant P. falciparum pre-mRNA processing factor 19 and interaction with recombinant PfDDI1

The complete coding region of *P. falciparum* pre-mRNA processing factor 19 (PfPRP19) was amplified from *P. falciparum* genomic DNA using primers PfPPF19-Fexp/PfPPF19-Rexp and cloned into the pGEX-6P-1 plasmid at BamHI-XhoI to obtain pGEX-PfPRP19, which was sequenced to confirm the coding region and transformed into BL21(DE3) *E. coli* cells. pGEX- PfPRP19 would express PfPRP19 as an N-terminal GST-tagged protein, facilitating purification of the recombinant _GST_PfPRP19 protein using the glutathione agarose resin. A pGEX-PfPRP19 expression clone was induced with IPTG (1.0 mM) at OD_600_ of 0.6 for 12 hours at 25°C. The induced cell pellet of was resuspended in lysis buffer (PBS with 1 mg/ml lysozyme, at 5 ml/g weight of the pellet), incubated for 30 min at 4°C, sonicated using the SONICS Vibra-Cell ultrasonic processor (5 secs pulses at 20% amplitude for 5-30 min), and centrifuged 39191g for 1 hr at 4°C. The supernatant was applied on a 1.0 ml GSTrap 4B column (GE healthcare), the column was washed with PBS, and bound proteins were eluted (50 mM Tris-Cl, 20 mM GSH, pH 8). The elutes were run on 12% SDS-PAGE, the fractions enriched with _GST_PfPRP19 were concentrated (Amicon Ultra centricon: 50 kDa cut off), quantitated using BCA and used for overlay assay. Different amounts of recombinant PfDDI_Myc/His_ were immobilized on nitrocellulose membrane, the membrane was blocked (3% BSA in PBST), overlaid with recombinant GST or _GST_PfPRP19 (2.5 µg/ml in blocking buffer) for overnight at 4°C, and washed with blocking buffer. The membrane was incubated with mouse anti-GST antibodies, followed by HRP-conjugated goat anti-Mouse IgG as described in the western blotting section.

### 2.13. Effect of DNA damaging chemicals on PfDDI1 knock-down parasites and chromatin association of PfDDI

Synchronized PfKD parasites was grown with or without trimethoprim till mid trophozoite stage, supplemented with DMSO (0.005%) or DNA damaging chemicals (0.1 mM hydroxyurea and 0.0005% MMS) and grown for 6 more hours. The parasites were washed with cell culture medium, grown in fresh culture medium under the initial trimethoprim conditions for 12 hours, and total parasitemia was determined by observing Giemsa smears. The parasitemias of hydroxyurea- and MMS-treated cultures were normalized as percent of DMSO-treated culture. The significance of difference (p-value) between two groups was determined using the t-test.

We carried out chromatin enrichment for proteomics (ChEP) to investigate association of PfDDI1 with chromatin as has been reported earlier (Batugedara et al., 2020; Kustatscher et al., 2014). A 50 ml synchronous culture of PfKD parasites (∼10% parasitemia) was grown in the presence of 10 μM trimethoprim till 30 hour stage. The culture was divided into two equal parts, one part was grown with MMS (0.005% volume/volume) and the other part was grown with DMSO (0.05% volume/volume) for 6 hours. The cultures were harvested and parasites were purified as mentioned in the parasite culture section. The parasite pellets were resuspended in 1% formaldehyde (in PBS), incubated for 10 minutes with shaking at 55 rpm and 37°C, and centrifuged 2300g. The parasite pellet was resuspended in 0.125 M glycine and incubated for 5 minutes at room temperature. The suspension was centrifuged at 2300g for 5 minutes at room temperature, the cells were washed with PBS, the parasite pellet was resuspended in 500 µl of nuclear extraction buffer (25 mM Tris-Cl, 10 mM KCl, 0.1 mM EDTA, 0.1 mM EGTA, 1 mM DTT, pH 7.4; supplemented with protease inhibitor cocktail and 0.25% NP-40), and incubated for 30 minutes in ice. The lysate was passed twice through a 25G needle, centrifuged at 2300g for 20 minutes at 4°C, and the supernatant (cytoplasm) was transferred to a microcentrifuge tube. The pellet, which contains nuclei, was resuspended in 500 µl of nuclear extraction buffer containing RNAse A (200 µg/ml) and incubated for 15 minutes at 37°C. The suspension was centrifuged at 2300g for 10 minutes at 4°C, the supernatant was discarded, and the pellet was washed twice with PBS to remove non cross-linked proteins. The pellet was resuspended in 500 µl of Tris-SDS buffer (50 mM Tris-Cl, 10 mM EDTA, pH 7.4, 4% SDS, protease inhibitor cocktail) using a hydrophobic pipette tip, and incubated for 10 minutes at room temperature. 1.5 ml of urea buffer (10 mM Tris-Cl, 1 mM EDTA, 8 M urea, pH 7.4) was added to the suspension and the sample was centrifuged at 15000g for 30 minutes at room temperature. The supernatant was discarded and the pellet was resuspended in 500 µl of Tris-SDS buffer, mixed with 1.5 ml urea buffer, and centrifuged at 15000g for 25 minutes at room temperature. The supernatant was discarded, the pellet was resuspended in 2 ml of Tris-SDS buffer, centrifuged at 15000g for 25 minutes at room temperature. The supernatant was discarded, the pellet was resuspended in 500 µl of storage buffer (10 mM Tris-Cl, 1 mM EDTA, 25 mM NaCl, 10% glycerol, pH 7.4, and protease inhibitor cocktail), and the suspension was sonicated (Diagenode Bioruptor UCD-200) at 4°C for 5 minutes to solubilize chromatin completely. The sonicated sample was centrifuged at 15000g for 30 minutes at 4°C, the supernatant containing the solubilized chromatin was transferred to a microcentrifuge tube, mixed with equal volume of SDS-PAGE sample buffer, and incubated at 98°C for 30 minutes to reverse the cross-links. The chromatin and the cytoplasmic samples were assessed for purity by western blotting using rabbit anti-Histone 2B (1/1000 diluted in blocking buffer) and rabbit anti-α2 subunit of 20S proteasome (at 1/1000 dilution in blocking buffer) antibodies, followed by appropriate secondary antibodies. Both chromatin and cytoplasmic samples were assessed for the presence of DDI1_Myc_/CDD_HA_ protein by western blot using mouse anti-Myc antibodies, followed by HRP-conjugated goat anti-mouse IgG as described in the western blotting section. The ImageJ software was used to quantitate full- length DDI1_Myc_/CDD_HA_ protein levels in western blots, and plotted using the GraphPad Prism software.

### 2.14. Isolation and assays with DNA-protein crosslinks

DPCs were generated and isolated from human embryonic kidney cells (HEK293T), wild type *P. falciparum* 3D7 and PfKD parasites as has been previously reported (Hu et al., 2020). HEK293T cells were cultured in DMEM (supplemented with 10% FBS, penicillin and streptomycin) at 37°C in a CO_2_ incubator. The cells were treated at 70-80% confluency with 50 µM etoposide or DMSO (0.001%) and grown further for 12 hours at 37LJC in a CO_2_ incubator. The culture medium was aspirated, 3 ml of the lysis buffer (5.6 M guanidine thiocyanate, 10 mM Tris-Cl, 20 mM EDTA, 4% Triton X-100, 1% sarkosyl, 1% DTT, pH 6.5) was added over the HEK293T cells (∼8×10^6^), and the cells were scraped off the flask. The cell suspension was mixed by pipetting using a 1 ml pipette tip and passed 6-8 times through a 24G needle. 3 ml of the chilled ethanol (-20°C) was added to the lysate and centrifuged at 21,000g for 20 minutes at 4°C. The pellet was washed twice with cold wash buffer (20 mM Tris-Cl, 150 mM NaCl, pH 6.5, 50% ethanol), solubilized in 1 ml of pre-warmed solution 1 (1% SDS, 20 mM Tris-Cl, pH 7.5) at 42°C for 6 minutes, sheared by passing 5-6 times through a 26G needle, and mixed with 1 ml of the precipitation buffer (200 mM KCl, 20 mM Tris-Cl, pH 7.5). The sample was incubated on ice for 6 min and centrifuged at 21,000g for 5 min at 4°C. The DPC pellet was washed twice with the low-salt buffer (100 mM KCl, 20 mM Tris-Cl, pH 7.5), each time by incubating at 55°C for 10 min, followed by on ice for 6 min, and centrifuged at 21,000g for 5 min at 4°C. The DPC pellet was resuspended in PBS and incubated at 37°C to solubilize. The amount of DPC- associated DNA was estimated at OD_260._ For isolation of DPCs from the parasite, a synchronized 100 ml ring stage culture of *P. falciparum* 3D7 (∼10% parasitemia) was divided into two equal parts and grown till 24 hour stage. One part was supplemented with 50 μM etoposide and another part with DMSO (0.001%). Both the cultures were grown for another 10 hours, parasites were purified and processed for isolation of DPCs as has been described for HEK293T cells.

We assessed PfKD parasites for the presence of DPCs and the association of PfDDI1 with DPCs. A synchronized 140 ml ring stage culture of PfKD parasites (∼15% parasitemia) was divided into two equal parts. One part was grown with trimethoprim (10 μM) and another without trimethoprim. 50 ml of each culture was harvested at 30 hour stage (30 hour time point) and parasites were purified. The remaining 20 ml culture was expanded to 50 ml with fresh medium and RBCs, grown under the same trimethoprim condition for another 48 hours, and the parasites were purified (78 hour time point). The parasite pellets were processed for isolation of DPCs and DPC-associated DNA. DPCs were isolated as has been described for HEK293T cells. For DPC-associated DNA, parasites were processed upto the precipitation step (precipitation with 200 mM KCl, 20 mM Tris-Cl, pH 7.5) as has been described for HEK293T cells. After precipitation, the supernatant was processed for isolation of free DNA and the pellet was processed for isolation of DPC-associated DNA. The DPC pellet was washed twice, each time with 1.5 ml of the low-salt buffer (100 mM KCl, 20 mM Tris-Cl, pH 7.5) by incubating at 55°C for 10 min, followed by on ice for 6 min, and centrifuged at 21,000g at 4°C for 5 min. The washes were pooled with the supernatant from the precipitation step. The DPC pellet was resuspended in 1 ml of proteinase K buffer (100 mM KCl, 20 mM Tris-Cl, pH 7.5 and 10 mM EDTA, proteinase K at 0.2 mg/ml), incubated at 55°C for 45 min, followed by on ice for 6 min. The sample was centrifuged at 21000g for 10 min at 4°C, the pellet (debris and proteins released from the digestion of DPCs) was discarded and the supernatant that contains the DPC-associated DNA was retained. The amount of DNA in DPC-associated DNA sample (supernatant of the proteinase K digested sample) and DPC-free DNA sample (pooled supernatant and washes) was quantitated using the SYBR green dye as per the manufacturer’s instructions. The amount of DPC-associated DNA was expressed as the percentage of total DNA (DNA present in the DPC sample and DPC-free DNA sample). For detection of DPC-associated PfDDI1, the 78 hour time point DPC preparations were spotted on a nitrocellulose membrane. The membrane was blocked (3% BSA in TBST), incubated with mouse anti-HA antibodies, followed by HRP-conjugated goat anti-mouse IgG antibodies, and signal was developed as has been described in the western blotting section.

We assessed DPCs for interaction with recombinant PfDDI1. DPC preparations from the DMSO-treated and etoposide-treated wild type *P. falciparum* 3D7 parasites and HEK293T cells were spotted on a nitrocellulose membrane, and the membrane was blocked (3% BSA in TBST). The membranes with HEK293T DPC spots were assessed for the presence of DPCs of topoisomerase II and histone 2B (as a chromatin control) using antibodies to rabbit human topoisomerase II and rabbit anti-H2B, followed by HRP-conjugated goat anti-rabbit IgG. For interaction of recombinant PfDDI1 with DPCs, pure parasite genomic DNA was also spotted as a control along with the wild type *P. falciparum* DPC preparations. These membranes were overlaid with purified recombinant PfDDI1_Myc/His_ (2.5 μg/ml in blocking buffer) for 12-14 hours at 4°C, washed with blocking buffer, incubated with rabbit anti-His antibodies, washed with blocking buffer and then with HRP-conjugated goat anti-rabbit IgG antibodies. The blots were developed as has been described in the western blotting section. The signal intensity DPC-associated native PfDDI1 or recombinant PfDDI1 was measured using the ImageJ software, and plotted using the GraphPad Prism software.

### 2.15. Statistical analysis

The data were analyzed to determine significance of difference (p-value) between two groups using the t-test or two-way ANOVA.

## 3. Results

### 3.1. DDI1 proteins of apicomplexan parasites exhibit diverse domain organizations

*S. cerevisiae* DD1 (ScDDI1) is the most studied among all DDI1 proteins, and we used it as a query sequence to search the *P. falciparum* genome database (PlasmoDB). A protein encoded by the PF3D7_1409300 gene and annotated as “DNA damage-inducible protein 1, putative” came out as the closest homolog. This protein was termed as the *P. falciparum* DNA damage-inducible protein 1 (PfDDI1), as it shares 33% identity with ScDDI1, contains putative UBL and RVP domains, and has a putative aspartyl protease motif (FVDSGA) in the RVP domain (Fig. S1). Canonical aspartyl proteases like pepsin contain two copies of the catalytic motif “VFDTGS” (Xaa-Xaa-Asp-Xbb-Gly-Xcc, where Xaa is a hydrophobic residue, Xbb is Thr or Ser, and Xcc is Ser), which together form the active site. The HIV aspartyl protease or retroviral aspartyl proteases possess a single catalytic motif (LLDTGA in HIV protease), hence, exists as a dimer in which the catalytic motif of both the monomers form the active site (Wlodawer and Erickson, 1993). DDI1 proteins, including PfDDI1, contain a single catalytic motif (Fig. S1). The X-ray crystal structures of RVP domains of ScDDI1, HsDDI2 and LmDDI1 indicate that they exist as dimers in which catalytic motifs of both the monomers form the active site (Kumar and Suguna, 2018; Sirkis et al., 2006; Trempe et al., 2016). Since a full-length DDI1 structure is not available yet, we compared the AlphaFold structure of full-length PfDDI1 (AF-Q8IM03-F1) with ScDDI1 UBL and RVP domains (PDB IDs: 2N7E for ScDDI1 UBL, 2I1A for ScDDI1 RVP), which were found to be comparable (RMSD: 1.165 for UBL and 0.544 for RVP domains; Fig. S2) and support multi-domain architecture of PfDDI1.

*Plasmodium* species encode for a single DDI1 ortholog, which have the same domain architecture and are 69-94% identical at amino acid level. The apicomplexan parasites sequenced to date also encode a single DDI1 ortholog, which vary in size, sequence and domain architecture. The catalytic motif in the majority of apicomplexan DDI1 proteins is “FVDSGA”, whereas it is “LVDTGA” in *Theileria*, *Cytauxzoon* and the majority of *Babesia* species. Interestingly, *Babesia microti* contains “FVDTGA”, which is a chimera of “FVDSGA” and “LVDTGA”. Analysis of the Apicomplexan DDI1 protein sequences for conserved domains and structurally similar domains revealed diversity in their domain architectures (Fig. 1A). All Apicomplexan DDI1 proteins contain the UBL and RVP domains. DDI1 proteins from the parasites of Sarcocystidae (*Besnoitia, Cystoisospora, Hammondia, Neospora, Sarcocystis* and *Toxoplasma*), Cryptosporidiidae (*Cryptosporidium*) and Theileriidae (*Theileria* and *Cytauxzoon*) families also contain the UBA domain. The DDI1 proteins of Babesiidae parasites contain all three domains (*Babesia microti*), UBL and RVP domains (*Babesia bovis*) or DUF676, UBL and RVP domains (*Babesia divergens* and *Babesia bigemina*). The sizes of Apicomplexan DDI1 proteins vary from 275 amino acid residues (*Gregarina niphandrodes*) to 1191 amino acid residues (*Babesia bigemina*). We also performed homology modelling of the representative DDI1 proteins listed in Fig. 1 (data not shown), which was consistent with the conserved domain analysis-based prediction. Homology modelling predicted UBL domain in *Gregarina niphandrodes* DDI1, and UBA domain in *Theileria annulata* and *Cytauxzoon felis* DDI1 proteins (Fig. S2), which could not be predicted using the conserved domain analysis. The diversity in domain architecture may impart species-specific functions or regulation of DDI1 proteins in Apicomplexan parasites. The Apicomplexan DDI1 proteins clustered on different branches in a phylogenetic tree (Fig. 1B), indicating sequence diversity across the Apicomplexa phylum. Nonetheless, the phylogram reflected evolutionary relatedness between different parasites, as DDI1 proteins of the same family cluster on the same branch and those of the same order share the root. For example, parasites of the families Babesiidae and Theileriidae in the order Piroplasmida share a common root.

**Fig. 1.**
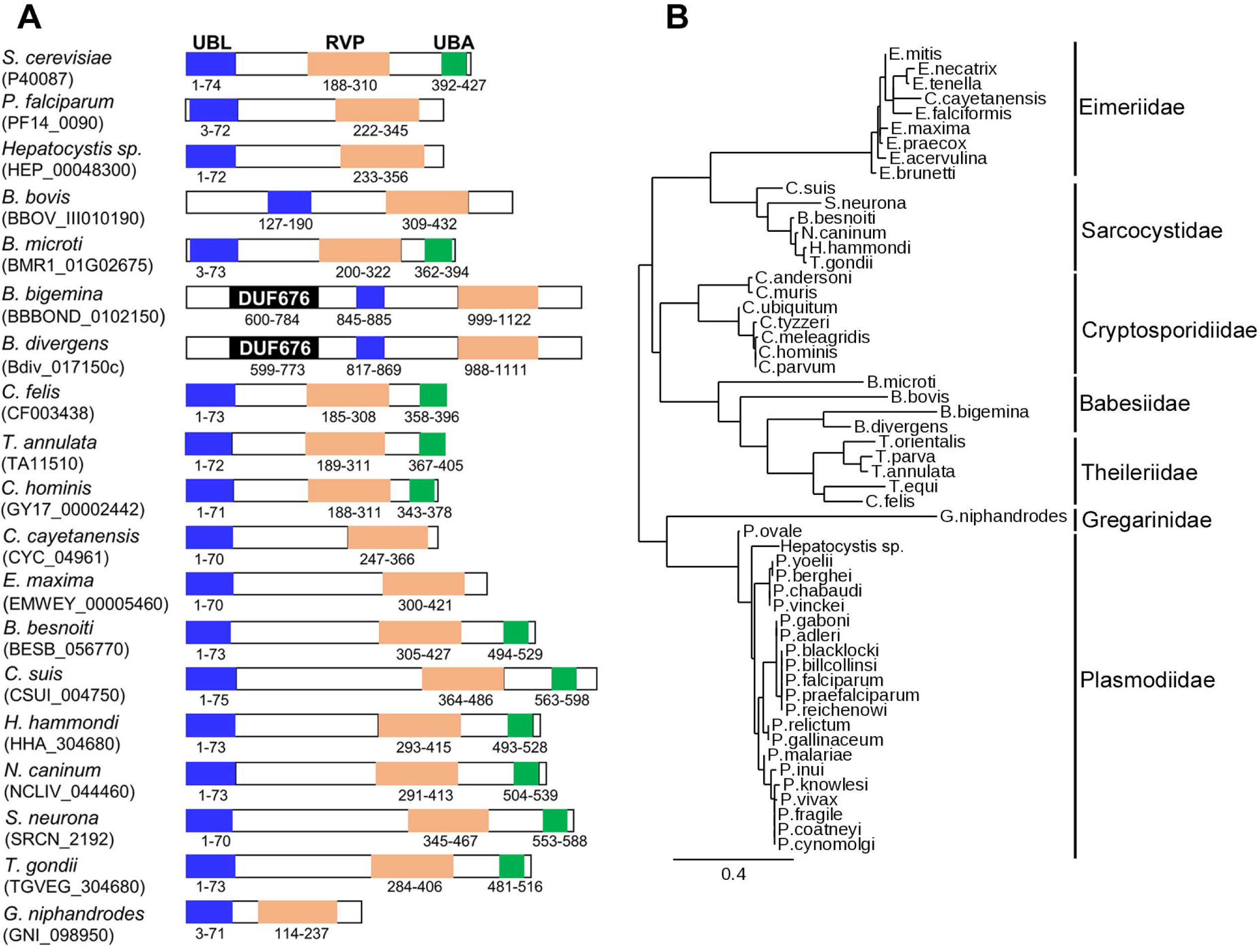
Domain organization and phylogenetic relatedness of DDI1 proteins. **A.** Sequences of the DDI1 proteins of indicated Apicomplexan parasites were analyzed for the presence of conserved domains and compared with the *S. cerevisiae* DDI1 domain architecture. The location and position of boundary amino acids of UBL, RVP and UBA domains are shown. The length of each schematic is sized according to the length of respective DDI1 protein except for *B. bigemina* and *B. divergens*. **B.** The sequences of 54 DDI1 proteins of the indicated parasite species of different families were used for generation of the phylogram.

### 3.2. Plasmodium DDI1 is expressed in all major parasite stages

The full-length PfDDI1 coding region was expressed as an N-terminal His-tagged protein in M15(pREP4) cells using the pQE30 vector, purified under denaturing conditions, refolded and used for antibody generation in rats (Fig. S3). The antiserum was adsorbed on M15(pREP4) cells to remove cross-reactive antibodies to *E. coli* proteins, and the adsorbed PfDDI1 antiserum was used to determine the expression and localization of PfDDI1 in different parasite stages. The PfDDI1 antiserum did not react with uninfected RBC lysate, but detected a prominent band of the size of full-length PfDDI1 (∼43.8 kDa) in the western blot of *P. falciparum* lysates (Fig. 2A), confirming expression in these stages, which was more predominant in ring and late trophozoite/schizont stages than early and mid trophozoite stages. For cellular localization, we performed immunofluorescence assay (IFA) of *P. falciparum* asexual and sexual erythrocytic stages with PfDDI1 antiserum, which indicated expression of PfDDI1 in both asexual and sexual stages (Fig. 2B and 2C). To check if DDI1 is expressed in insect and liver stages, we turned to *P. berghei*, which offers studies in insect and liver stages with ease compared to *P. falciparum*. As PfDDI1 and *P. berghei* DDI1 (PbDDI1) share 70.5% identity, PfDDI1 antiserum detected PbDDI1 in the western blot of trophozoites and showed prominent signal in the IFA of trophozoite, sporozoite and liver stage schizont of *P. berghei* (Fig. 3), indicating expression of PbDDI1 in these stages. The localization of both PfDDI1 and PbDDI1 appears to be throughout the parasite except the food vacuole in all stages. In addition to the full-length PfDDI1, we also observed smaller size bands in the western blots, which could be the processed form of full-length protein. The western blot and IFA data of *P. falciparum* and *P. berghei* DDI1 proteins together indicate that *Plasmodium* DDI1 is expressed in all major developmental stages of malaria parasites.

**Fig. 2.**
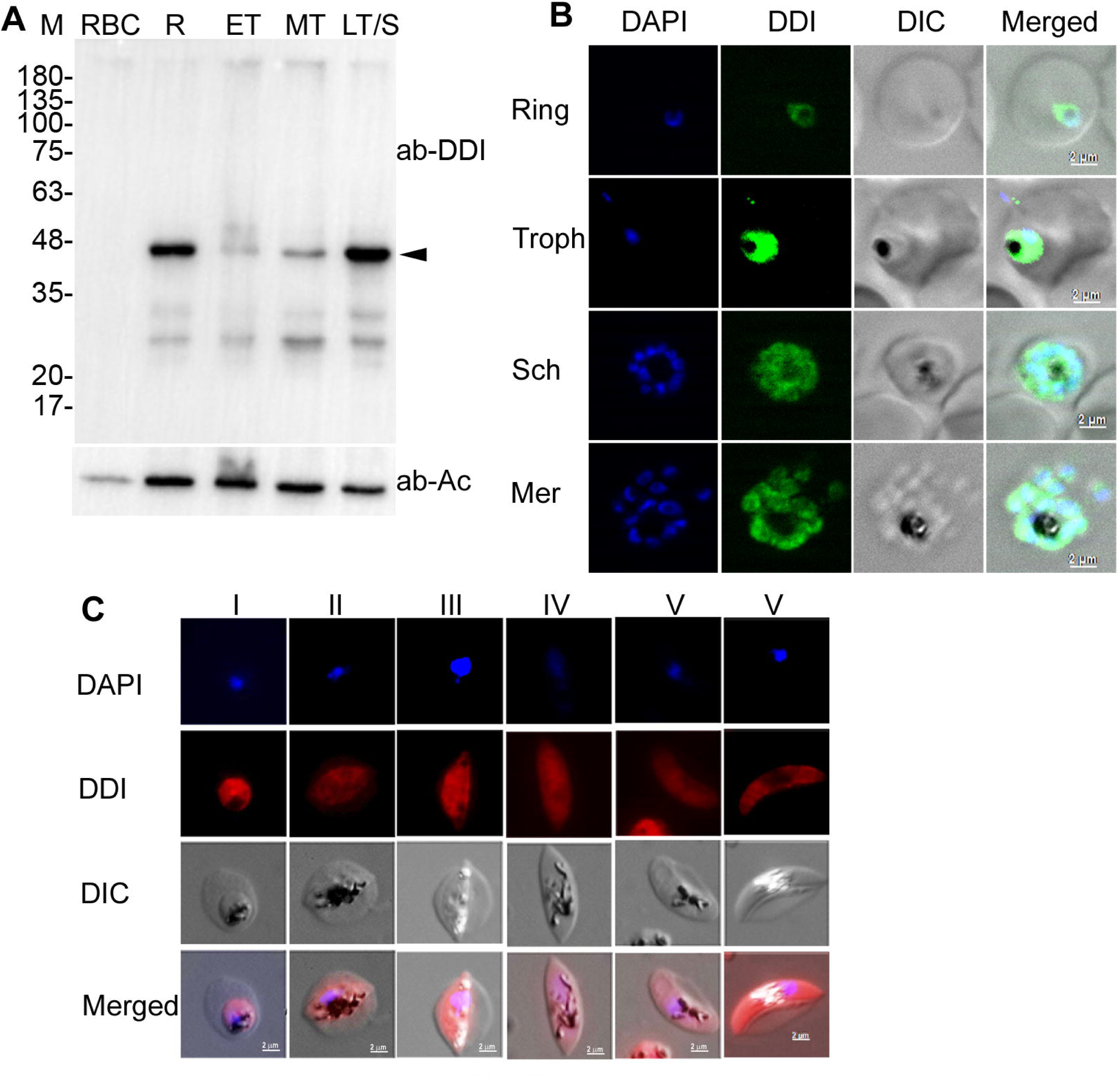
Expression and localization of PfDDI1 in asexual and sexual erythrocytic stages. A synchronized culture of *P. falciparum* was harvested at different stages and evaluated for expression and localization of PfDDI1. **A.** The western blot of lysates of ring (R), early trophozoite (ET), mid trophozoite (MT) and late trophozoite/schiziont (LT/S) parasite stages and uninfected red blood cells (RBC) was probed using PfDDI1 antiserum (ab-DDI). β-actin (ab-Ac) was used as a loading control. The position of full-length PfDDI1 is indicated by the arrowhead and protein size markers are in kDa (M). **B**. The ring, trophozoite (Troph), schizont (Sch) and free merozoite (Mer) stages were assessed for localization of PfDDI1 by IFA using PfDDI1 antiserum. The panels show nucleic acid staining (DAPI), PfDDI1 signal (DDI), bright field with RBC and parasite boundaries (DIC) and the overlap of all three images (Merged). The black substance is haemozoin, a food vacuole-resident pigment. **C.** The indicated gametocyte stages of *P. falciparum* were evaluated for expression and localization of PfDDI1 by IFA using PfDDI1 antiserum. The panels are as in B. The size scale bars are shown in the merged image.

**Fig. 3.**
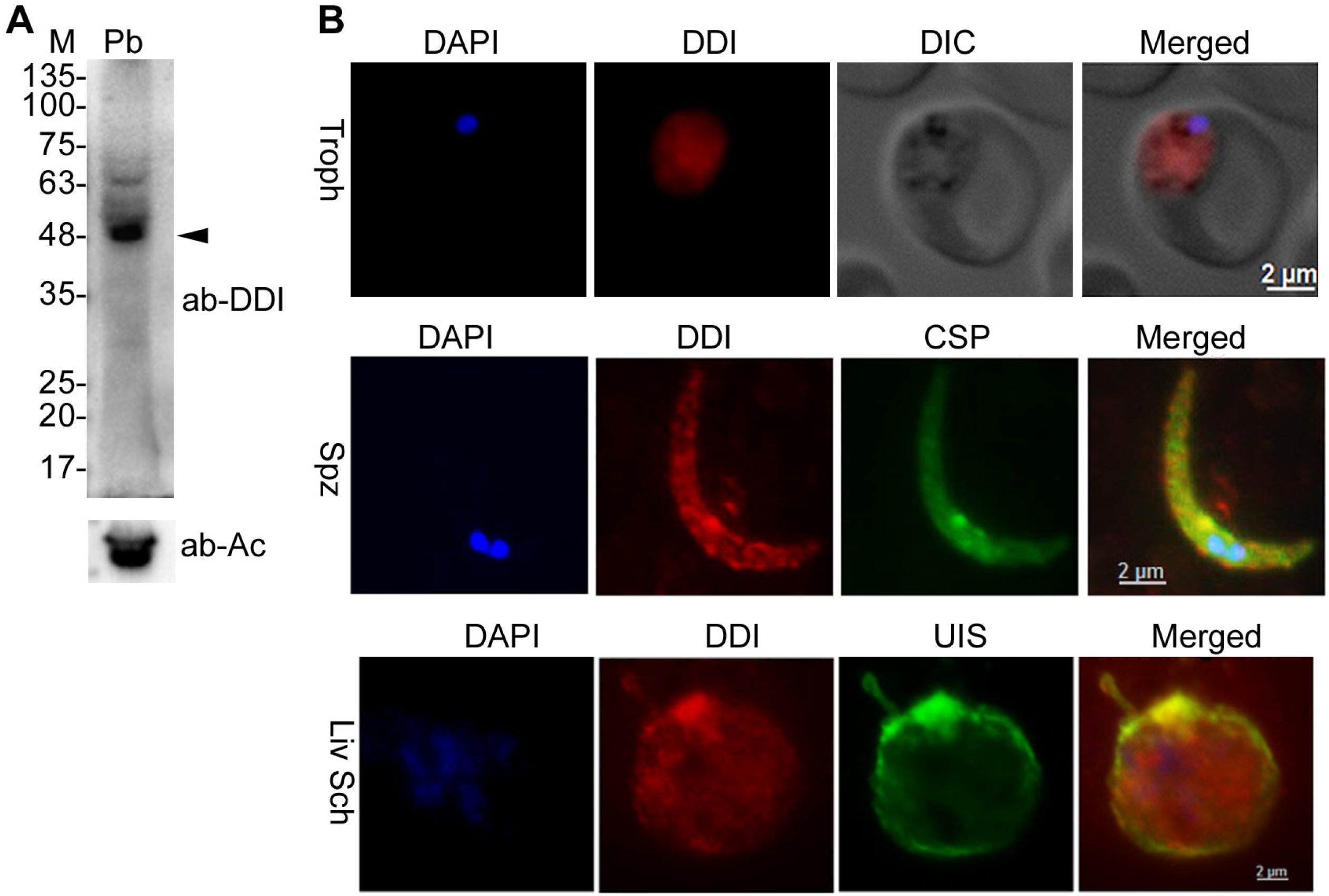
Expression and localization of PbDDI1. **A.** The western blot of *P. berghei* asexual erythrocytic stage parasites was probed for PbDDI1 expression using PfDDI1 antiserum (ab- DDI), and β-actin (ab-Ac) was used as a loading control. The arrowhead indicates the predicted full-length PbDDI1 (∼44.3 kDa). Protein size markers are in kDa (M). **B.** *P. berghei* erythrocytic trophozoite (Troph), salivary gland sporozoite (Spz) and liver schizont (Liv Sch) stages were evaluated for localization of PbDDI1 by IFA using PfDDI1 antiserum along with antibodies to CSP (a sporozoite membrane marker) and UIS (a liver stage parasitophorous vacuole marker). The panels are for nucleic acid staining (DAPI), PbDDI1 signal (DDI), signal for CSP and UIS, bright field with RBC and parasite boundaries (DIC), and the overlap of three images (Merged). The size scale bars are shown in the merged image.

### 3.3. DDI1 is critical for parasite development

Expression of DDI1 in all the major parasite stages suggests its importance for parasite development. Hence, we attempted to knock-out the PbDDI1 gene for investigation of its functions during parasite development. However, multiple knock-out attempts were unsuccessful. We next employed a conditional knock-down approach in *P. berghei* by replacing the wild type PbDDI1 coding region with PfDDI_Myc_/cDD_HA_ coding sequence, which would express fusion of PfDDI_Myc_ with HA-tagged mutant *E. coli* DHFR (cDD_HA_). cDD binds trimethoprim and is stable, but undergoes proteasomal degradation in the absence of trimethoprim, thereby, causing a knock-down effect. The replacement of wild type PbDDI1 with PfDDI_Myc_/cDD_HA_ and expression of DDI_Myc_/cDD_HA_ fusion protein were successful (Fig. S4), which confirmed that PbDDI1 and PfDDI1 are functionally conserved. The knock-down parasites (PbKD) showed reduction in DDI_Myc_/cDD_HA_ protein level in the absence of trimethoprim compared to that in the presence of trimethoprim, indicating the knock-down effect (Fig. 4A). PbKD parasites grew similar to wild type parasites in the presence of trimethoprim, but showed drastically reduced growth in the absence of trimethoprim and eventually disappeared (Fig. 4B). In the presence of trimethoprim, PbKD-infected mice had to be euthanized, whereas in the absence of trimethoprim, the parasitemia barely reached to 0.5% and the infection was self-limiting. Withdrawal of trimethoprim from PbKD-infected mice at about 5% parasitemia also resulted in complete clearance of parasites (Fig. 4C).

**Fig. 4.**
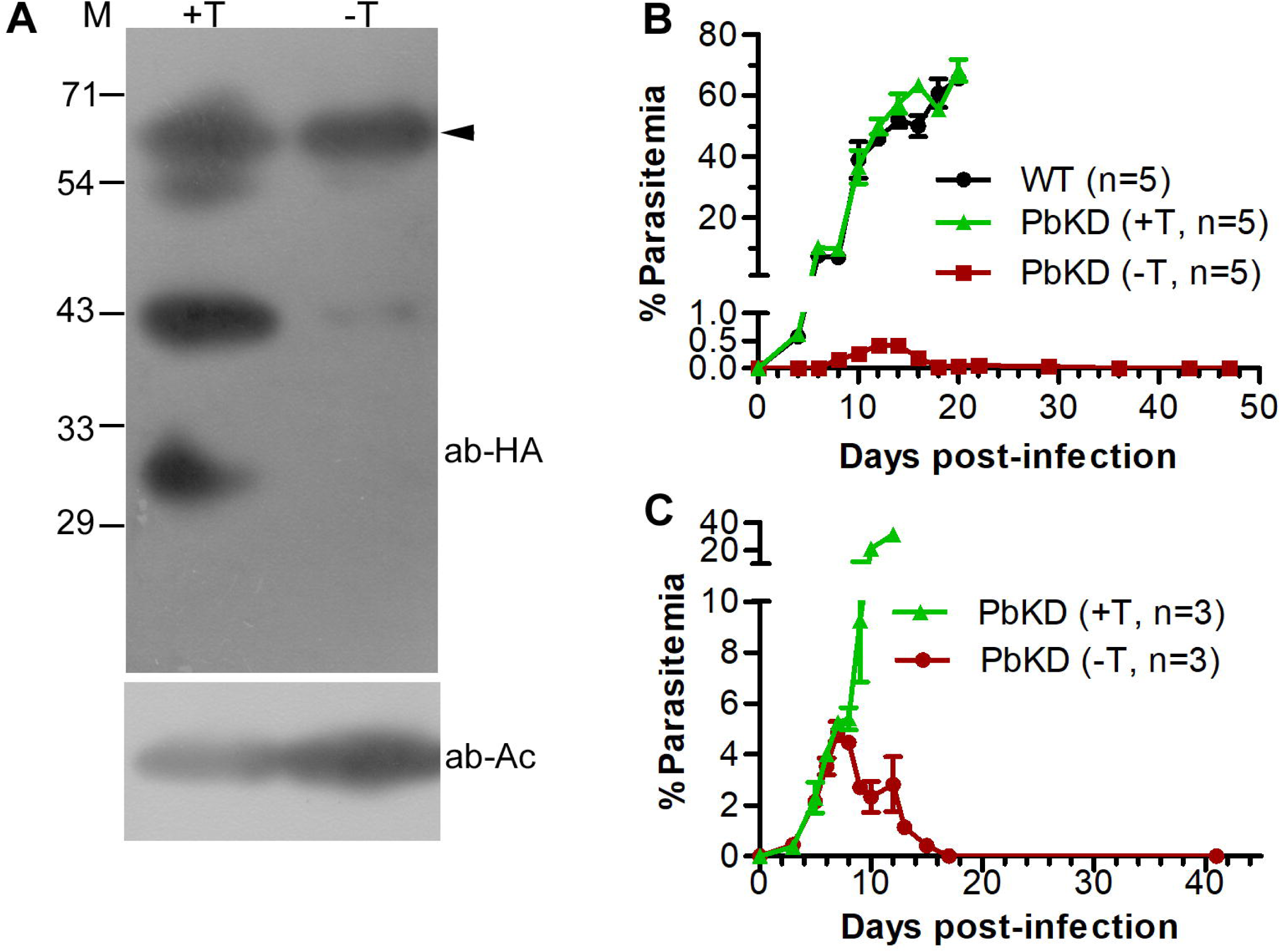
DDI1 is critical for development of *P. berghei*. **A.** DDI1 knock-down *P. berghei* parasites (PbKD) were cultured in the presence (+T) or absence (-T) of trimethoprim, and parasite lysates were assessed for DDI_Myc_/cDD_HA_ protein level by western blotting using antibodies to HA (ab-HA) and β-actin (ab-Ac) as a loading control. The top band indicated by the arrowhead corresponds to the predicted size of full-length DDI_Myc_/cDD_HA_ (∼64.4 kDa) and the sizes of protein markers are in kDa (M). **B.** Mice were infected with equal number of wild type *P. berghei* ANKA (WT) or PbKD parasites and infection was monitored. The mice infected with PbKD parasites were kept under (+T) or without (-T) trimethoprim. The number of mice in each group is shown with “n”, and parasite growth is shown as % parasitemia over days post- infection. **C.** Mice were infected with PbKD parasites and kept under trimethoprim until ∼5% parasitemia. Trimethoprim was withdrawn from one group of mice (-T), whereas the other group was maintained under trimethoprim (+T). Infection was monitored, “n” is the number of mice in each group, and parasite growth is shown as % parasitemia over days post-infection.

We similarly generated a *P. falciparum* line expressing PfDDI_Myc_/cDD_HA_ in place of the wild type PfDDI1, and assessed the knock-down parasites (PfKD) for various parameters, including the parasite development. PfKD parasites showed successful gene replacement and expression of DDI_Myc_/cDD_HA_ fusion protein (Fig. S5). The PfKD parasites grown without trimethoprim showed significantly decreased DDI_Myc_/cDD_HA_ protein level as compared to those grown in the presence of trimethoprim, indicating the knock-down effect (Fig. 5A, B, C). Consistent with PbKD parasites, the growth of PfKD parasites was significantly reduced in the absence of trimethoprim as compared to that in the presence of trimethoprim (Fig. 5D), indicating that DDI1 is important for asexual erythrocytic stage development of *P. falciparum* as well. The knock-down effect was more prominent both as reduction in protein level and growth with increased period of growth without trimethoprim. Failure to knock-out the PbDD1 gene, significantly decreased growth of parasites upon knock-down of DDI1 protein, and loss of virulence of PbKD parasites indicate a critical role of DDI1 during asexual stage parasite development.

**Fig. 5.**
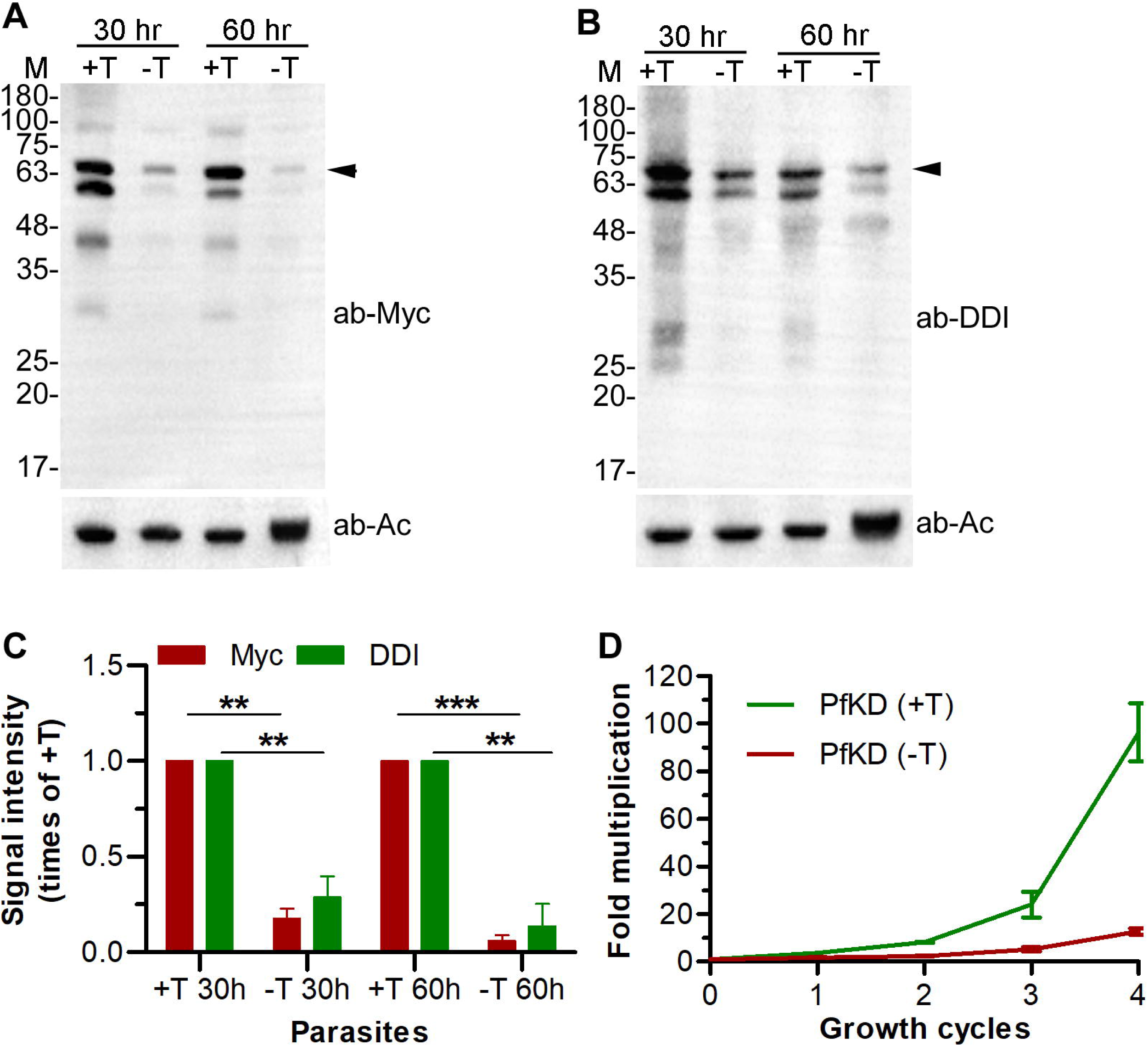
DDI1 is critical for *P. falciparum* development. Synchronized ring stage PfKD parasites were cultured in the presence (+T) or absence (-T) of trimethoprim, harvested at 30 hour and 60 hour stages of development, and processed for western blotting using anti-Myc antibodies (ab-Myc; **A**) or PfDDI1 antiserum (ab-DDI; **B**). β-actin (ab-Ac) was used as a loading control. The arrowhead indicates the predicted full-length DDI_Myc_/cDD_HA_ protein (∼64.4 kDa) and the sizes of protein markers are in kDa (M). Note that the blots A and B have nearly identical pattern of the full-length DDI_Myc_/cDD_HA_ protein and smaller size species, indicating that both the antibodies are specific for DDI_Myc_/cDD_HA_ and decreased level of DDI_Myc_/cDD_HA_ protein in -T parasite lysates is due to knock-down of the protein. **C.** The signal intensity of full-length DDI_Myc_/cDD_HA_ and β-actin bands in A and B were measured, and the signal intensity of full- length DDI_Myc_/cDD_HA_ was normalised with that of the corresponding β-actin band. The plot shows signal intensity of full-length DDI_Myc_/cDD_HA_ band in -T parasite lysates as times of the corresponding +T parasite lysates on y-axis for different parasite lysates (parasite) on x-axis. The data is mean of three independent experiments with SD error bar. **D.** PfKD parasites were cultured with (+T) or without (-T) trimethoprim for four consecutive cycles. Parasitemia was determined at the beginning and end of each growth cycle, normalized to the initial parasitemia, and shown as fold multiplication over growth cycles. The data is mean of two independent experiments with SD error bar, each done in triplicates.

### 3.4. Prior infection of mice with PbKD parasites protects from subsequent infection

Since mice infected with PbKD parasites resolved infection in the absence of trimethoprim, we assessed if the recovered mice developed immunity. Groups of naïve and recovered mice were challenged with a lethal inoculum of wild type *P. berghei* ANKA and monitored for infection. The naïve mice developed high parasitemia and succumbed to infection, whereas the recovered mice either did not develop infection or cleared infection after developing a low level of parasitemia (Fig. 6A). This indicated that prior exposure of mice to PbKD parasites conferred protective immunity. The recovered mice also showed protection against a lethal *P. yoelii* 17XNL challenge (Fig. 6B), which indicated that prior exposure of mice with PbKD parasites protects from a heterologous challenge as well. The pooled serum, which was collected from the recovered mice one day before the challenge, showed reactivity with soluble parasite extract in ELISA, with a titer as high as 1/128000, suggesting that anti-parasite antibodies could have contributed to protection from the subsequent parasite challenge.

**Fig. 6.**
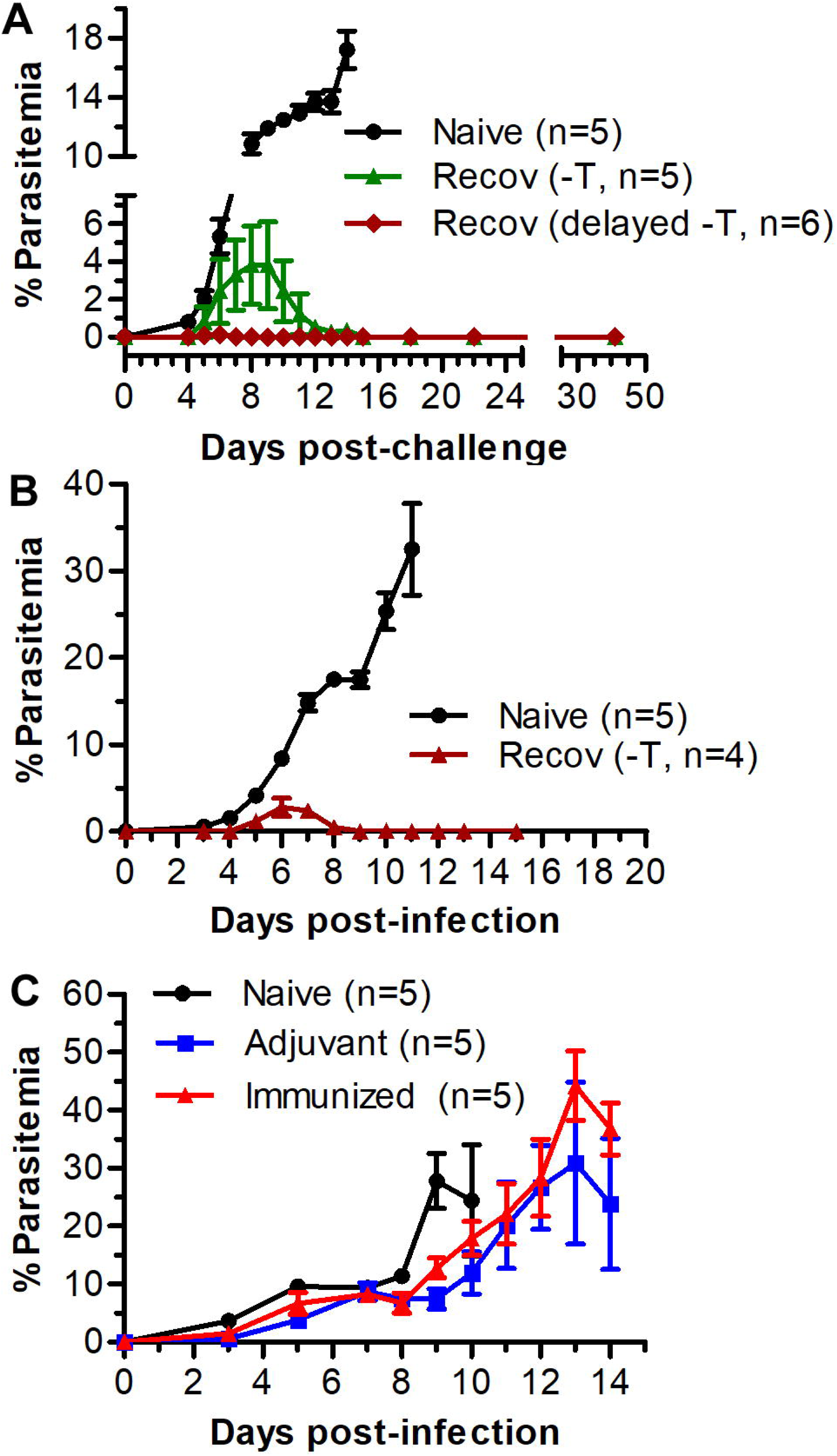
Prior infection of mice with PbKD parasites elicits protective immunity. **A.** Naïve and the recovered mice were challenged with equal number of wild type *P. berghei* ANKA, infection was monitored and shown as % parasitemia over days post-infection. The recovered group that was never given trimethoprim during PbKD parasite infection is indicated with “Recov (-T)” and those mice that recovered from PbKD infection after late withdrawal of trimethoprim are indicated with “Recov (delayed, –T)”. **B.** Naïve and recovered (Recov) mice were infected with equal number of *P. yoelii* 17XNL parasites, infection was monitored and shown as % parasitemia over days post-infection. **C.** Mice were immunized with recombinant PbDDI1+adjuvant (immunized), adjuvant only (adjuvant) or left without immunization (naïve). All the three groups of mice were challenged with equal number of *P. berghei* ANKA parasites, infection was monitored and shown as % parasitemia over days post-infection. For all three plots, “n” represents the number of mice and each data point represents the mean of % parasitemias of the respective group.

Owing to the critical role and prominent expression of PbDDI1 in blood stages, we produced recombinant PbDDI1 (Fig. S6), immunized mice with it and challenged the mice with wild type *P. berghei* ANKA. The mice immunized with recombinant PbDDI1 developed high antibody titer against recombinant PbDDI1 (ELISA antibody titer: 1/600000), but showed infection just like naïve and adjuvant control mice (Fig. 6C). This ruled out PbDDI1 as a natural target of immune response, and indicated that prior infection of mice with PbKD parasites elicits a protective immune response, most likely against the whole parasite.

### 3.5. DDI1 knock-down enhanced drug sensitivity of parasites

The inhibition of LmDDI1 by HIV protease inhibitors and conservation of the HIV protease-like retroviral aspartic protease fold in DDI1 proteins suggest that DDI1 could be a target of HIV protease inhibitors, which have been shown to block the development of all major stages of malaria parasites (Hobbs et al., 2013; Hobbs et al., 2009; Nsanzabana and Rosenthal, 2011a; Parikh et al., 2005; Perteguer et al., 2013; White et al., 2011b). Hence, we assessed PfKD parasites for susceptibility to various inhibitors and drugs, including the HIV protease inhibitors. The IC_50_ concentrations of the drugs/inhibitors tested were similar for wild type and PfKD parasites (grown in the presence of trimethoprim) (Table 1), indicating that mere genetic manipulation did not affect the parasite. On the other hand, PfKD parasites grown in the absence of trimethoprim showed 1.4-1.9 fold decrease in IC_50_ concentrations for artemisinin and HIV protease inhibitor lopinavir that was more prominent in the 2^nd^ cycle (Table 1), which is consistent with increased knock-down effect in the 2^nd^ cycle. E64 was similarly effective on all the parasites regardless of the presence or absence of trimethoprim, indicating that increased susceptibility to artemisinin and HIV protease inhibitor is specific to the knock-down of PfDDI1.

**Table 1.** Susceptibility of PfKD parasites to inhibitors and drugs. Wild type (WT) and PfKD parasites were assessed for susceptibility to inhibitors (cysteine protease inhibitor E64, proteasome inhibitor epoxomicin, antimalarial artemisinin and HIV protease inhibitors lopinavir, nelfinavir and saquinavir). For PfKD parasites, trimethoprim was maintained during the assay (+T), excluded during the assay (-T1) or excluded in the previous growth cycle and during the assay (-T2). The IC_50_ concentrations of the indicated compounds are mean with SD of at least three independent experiments, each done in duplicates. The p-value for significance of difference between the IC_50_ concentrations for WT and PfKD (-T2) parasites is indicated.

| Inhibitor | WT | PfKD (+T) | PfKD (-T1) | PfKD (-T2) | p-value for WT vs PfKD (-T2) |
| --- | --- | --- | --- | --- | --- |
| Art (nM) | 20.73 ± 0.17 | 21.90 ± 0.03 | 15.14 ± 0.10 | 13.66 ± 0.68 | 0.0006 |
| E64 (μM) | 2.25 ± 0.32 | 2.30 ± 0.10 | 1.85 ± 0.38 | 2.09 ± 0.27 | 0.5003 |
| Lopinavir (μM) | 2.31 ± 0.10 | 2.00 ± 0.21 | 1.23 ± 0.38 | 1.22 ± 0.06 | 0.0001 |
| Nelfinavir (μM) | 7.84 ± 0.99 | 7.97 ± 0.18 | 6.71 ± 0.44 | 6.49 ± 0.56 | 0.0314 |
| Saquinavir (μM) | 13.05 ± 0.33 | 13.42 ± 0.39 | 11.90 ± 0.79 | 11.80 ± 1.19 | 0.1418 |
| Epoxomicin (nM) | 9.32 ± 1.09 | 10.06 ± 1.06 | 8.86 ± 0.56 | 7.24 ± 0.69 | 0.0259 |

### 3.6. DDI1 associates with chromatin and DNA-protein crosslinks

ScDDI1 was first identified as one of the upregulated proteins in response to DNA damaging chemicals, and two recent studies showed roles of ScDDI1 in removing the replication termination factor RTF2 from stalled replication forks and DNA-topoisomerase complexes (Serbyn et al., 2020; Svoboda et al., 2019). Hence, we assessed the effect of DNA damaging chemicals like hydroxyurea and methyl methanesulfonate (MMS) on the development of PfKD parasites. The PfKD parasites showed increased susceptibility to both hydroxyurea and MMS under knock-down conditions as compared to the parasites grown under normal conditions (Fig. 7A). How would *Plasmodium* DDI1 protect from DNA damage? We hypothesized that PfDDI1 is likely present in the nucleus and associates with chromatin. To address the hypothesis, we prepared chromatin and cytoplasmic fractions of parasites grown with or without MMS by the chromatin enrichment for proteomics (ChEP) method, and assessed for the presence of PfDDI1. The ChEP procedure involves crosslinking of chromatin-associated proteins in intact cells, followed by treatment of the nuclear fraction with buffers containing SDS and urea, which would remove all the proteins not associated with chromatin and merely present in the nucleus. Western blot of the chromatin fraction showed the presence of H2B, a nuclear protein, and the absence of α2 proteasome subunit, a cytoplasmic protein (Fig. 7B). Similarly, western blot of the cytoplasmic fraction showed the presence of α2 proteasome subunit and absence of H2B (Fig. 7B), indicating purity of the respective fractions. PfDDI1 was present both in chromatin and cytoplasmic fractions, and had higher level in chromatin fraction of MMS-treated parasites than that of DMSO-treated control parasites (Fig. 7D), indicating that PfDDI1 is also present in the nucleus and associates with chromatin. The widefield IFA images showed PfDDI1 localization throughout the parasite except the food vacuole (Fig. 2B), and the confocal microscopy image showed colocalization of PfDDI1 with DAPI (Fig. 7C), indicating that PfDDI1 is present both in cytoplasm and nucleus, which is consistent with its presence in cytoplasmic and chromatin fractions. We next checked the effect of PfDDI knock-down on DNA-protein crosslinks (DPCs). We first optimized the procedure for preparation of etoposide-induced topoisomerase II-DPCs for HEK293T cells, as has been reported earlier (Fig. S7) (Hu et al., 2020). The same procedure was used to isolate DPCs from wild type *P. falciparum* 3D7 and PfKD parasites. The knock- down of PfDDI1 increased DPCs, as indicated by increased amount of DPC-associated DNA (Fig. 8A). DDI_Myc_/cDD_HA_ was detected in the DPC preparation from PfKD parasites, and had lesser level in DPC preparation from the PfKD parasites grown in the absence of trimethoprim than those grown in the presence of trimethoprim (Fig. 8B and Fig. S8A), which is consistent with the knock-down level of DDI_Myc_/cDD_HA_. Consistently, recombinant PfDDI1_Myc/His_ interacted with DPCs in a dose dependent manner, but not with purified parasite genomic DNA (Fig. 8C and 8D, Fig. S8B). Hence, association of PfDDI1 with chromatin and DPCs, and accumulation of DPCs upon knock-down of PfDDI1 indicate a role for it in DNA damage response and repair of DNA-protein crosslinks.

**Fig. 7.**
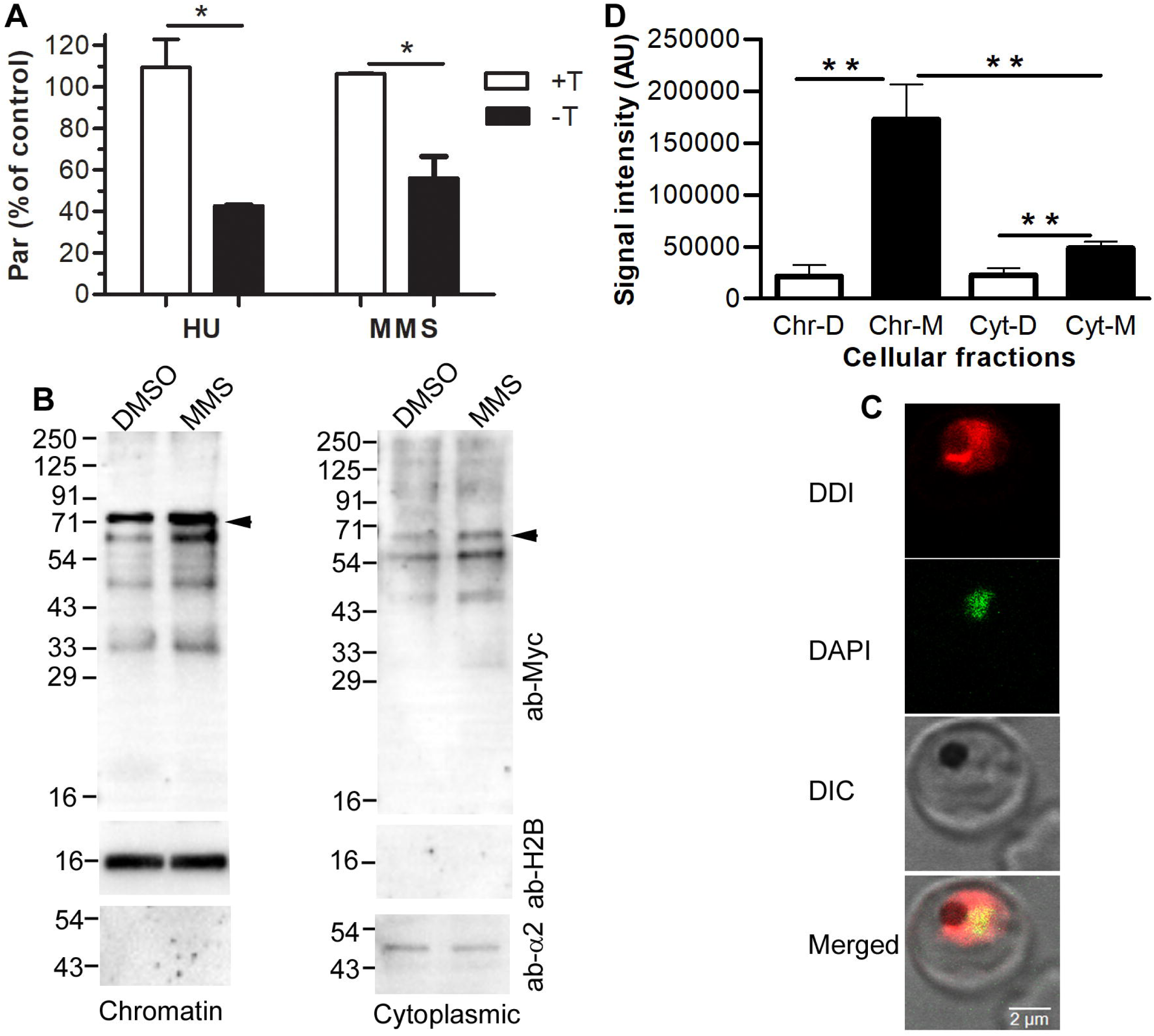
PfKD parasites show increased susceptibility to DNA damaging chemicals and DDI1 associates with chromatin. PfKD parasites were grown with (+T) or without (-T) trimethoprim and supplemented with hydroxyurea (HU), MMS or DMSO at mid trophozoite stage for 6 hours. The parasites were washed and grown in fresh medium (+T or –T) for 12 hours and total parasitemia was determined. **A.** The plot represents parasitemia as percent of DMSO-treated group (par (% of control)) for the respective group, and the data is average of two independent experiments with SD error bar. **B.** PfKD parasites were maintained in trimethoprim and treated with DMSO or MMS (0.005% v/v) at mid trophozoite stage for 6 hours. The parasites were purified and processed for separation of cytosolic and chromatin fractions. The chromatin and cytoplasmic fractions were evaluated for the presence of PfDDI1 by western blotting using anti-Myc antibodies (ab-Myc). Anti-histone 2B (ab-H2B) and anti-α2 proteasome subunit (ab-α2) antibodies were used to assess the purity of chromatin and cytoplasmic fractions, respectively. The arrowhead indicates the predicted full-length DDI_Myc_/cDD_HA_ protein (∼64.4 kDa) and the sizes of protein markers are in kDa (M). **C.** Wild type *P. falciparum* trophozoites were evaluated for nuclear localization PfDDI1 using PfDDI1 antiserum by confocal microscopy. The images show PfDDI1 signal (DDI), false color for nucleic acid staining (DAPI), bright field with RBC and parasite boundaries (DIC) and the overlap of all three images (Merged). The yellow signal in the merged panel indicates colocalization of PFDDI1 and DAPI, and the black substance is haemozoin, a food vacuole-resident pigment. **D.** The signal intensities of full-length DDI_Myc_/cDD_HA_ bands in “B” were measured, and plotted as arbitrary units (AU) on y-axis for the indicated cellular fractions on x-axis (chromatin (Chr-D) and cytoplasmic (Cyt-D) fractions of DMSO-treated parasites; chromatin (Chr-M) and cytoplasmic (Cyt-M) fractions of MMS-treated parasites). The data is mean of three independent experiments with SD error bar.

**Fig. 8.**
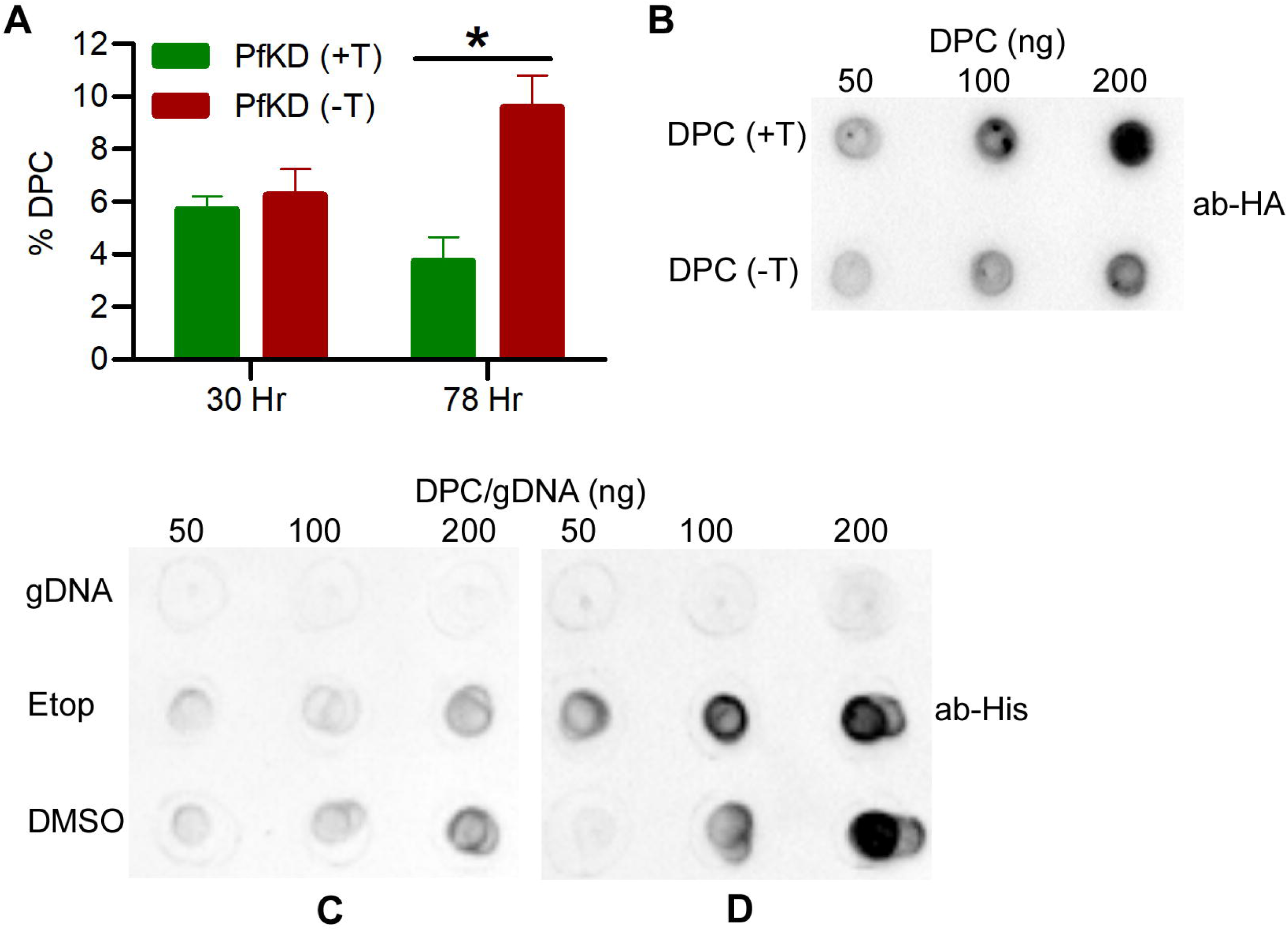
Accumulation and association of DPCs with PfDDI1. DPCs were prepared from PfKD and wild type *P. falciparum* 3D7 parasites, and assessed for the associated DNA, presence of PfDDI1_Myc_-cDD_HA_ and interaction with recombinant PfDDI1_Myc/His_. **A.** PfKD parasites were grown in the presence (+T) or absence (-T) of trimethoprim, harvested at 30 hr and 78 hr time points, and processed for isolation of DPC-associated DNA and free DNA. The graph shows DPC-associated DNA (% DPC) as percent of the total DNA. “*” represents p value (0.0321). **B.** The indicated amounts of 78 hr time point DPC samples from PfKD parasites (grown in the presence (+T) or absence (-T) of trimethoprim) were spotted on a nitrocellulose membrane, and evaluated for the presence of PfDDI_Myc_/cDD_HA_ using anti-HA antibodies (ab-HA). **C** and **D.** Wild type *P. falciparum* 3D7 parasites were treated with DMSO or etoposide (Etop) and DPCs were isolated. The indicated amounts of DPCs and purified parasite gDNA (DPC/gDNA) were spotted on a nitrocellulose membrane, the membrane was overlaid with BSA (C) or recombinant PfDDI1_Myc/His_ (D), and the membrane was probed using anti-His antibodies (ab-His).

### 3.7. DDI1 knock-down caused accumulation of ubiquitinated proteins

LmDDI1 has been shown to hydrolyze a HIV protease peptide substrate and BSA under acidic conditions (Perteguer et al., 2013), the HsDDI2 has been shown to cleave Nrf1 (Koizumi et al., 2016), and ScDDI1 has been shown to have ubiquitin-dependent protease activity (Yip et al., 2020). To investigate biochemical properties of PfDDI1, we expressed a synthetic and codon optimized PfDDI1 coding sequence as a C-terminal Myc/His-tag (PfDDI_Myc/His_) in Rosetta-gami 2(DE3)pLysS cells using the pET-26b(+) plasmid. Recombinant PfDDI_Myc/His_ was purified from the soluble fraction and used for various assays. The purified protein sample contained full- length PfDDI_Myc/His_ as the prominent species in addition to multiple smaller bands (Fig. S9A). The full-length protein species was separated from the smaller size species by gel filtration chromatography with a relative size of 158 kDa (Fig. S9B and S9C). The predicted size of PfDDI_Myc/His_ is 46.1 kDa, suggesting that recombinant PfDDI_Myc/His_ exists as an oligomer. However, we did not observe degradation of BSA by recombinant PfDDI_Myc/His_ in our experimental settings (Fig. S10A). Nonetheless, depletion of PfDDI1 caused accumulation of ubiquitinated proteins (Fig. S10B and S10C), suggesting a role in degradation of ubiquitinated proteins, which may be via itself and/or through the proteasome.

### 3.8. DDI1 interacts with pre-mRNA-processing factor 19

ScDDI1 has been shown to interact with t-SNARE and v-SNARE (Lustgarten and Gerst, 1999; Marash and Gerst, 2003), ubiquitin (Bertolaet et al., 2001a, b; Nowicka et al., 2015; Trempe et al., 2016), Ho endonuclease (Kaplun et al., 2005), and proteasome subunits. CeDDI1/Rngo has been shown to bind ubiquitin and the 26S proteasome (Morawe et al.). SpDDI1/Mud1 binds K48-linked polyubiquitin (Trempe et al., 2005). Hence, we immunoprecipitated PfDDI_Myc_/cDD_HA_ and subjected it to mass spectrometry to identify the interacting proteins. 62 proteins were reproducibly present in the immunoprecipitate from three biological repeats (Table S2), including the *P. falciparum* pre-mRNA-processing factor 19 (PfPRP19), proteins predicted to be associated with the ubiquitin proteasome system and protein translation. Since the human PSO4/PRP19 is an E3 ubiquitin ligase and has been shown to play critical roles in multiple DNA repair pathways and mRNA splicing (Mahajan, 2016; Yin et al., 2012), we produced recombinant PfPRP19 (Fig. S11), and assessed it for interaction with recombinant PfDDI1_Myc/His_ by protein-protein overlay assay. Both the recombinant proteins showed dose-dependent interaction (Fig. 9), indicating that these two proteins interact directly. We speculate that PfPRP19 may mediate chromatin association of PfDDI1, as PSO4/PRP19, but not PfDDI1, contain a nuclear localization and DNA-binding domains.

**Fig. 9.**
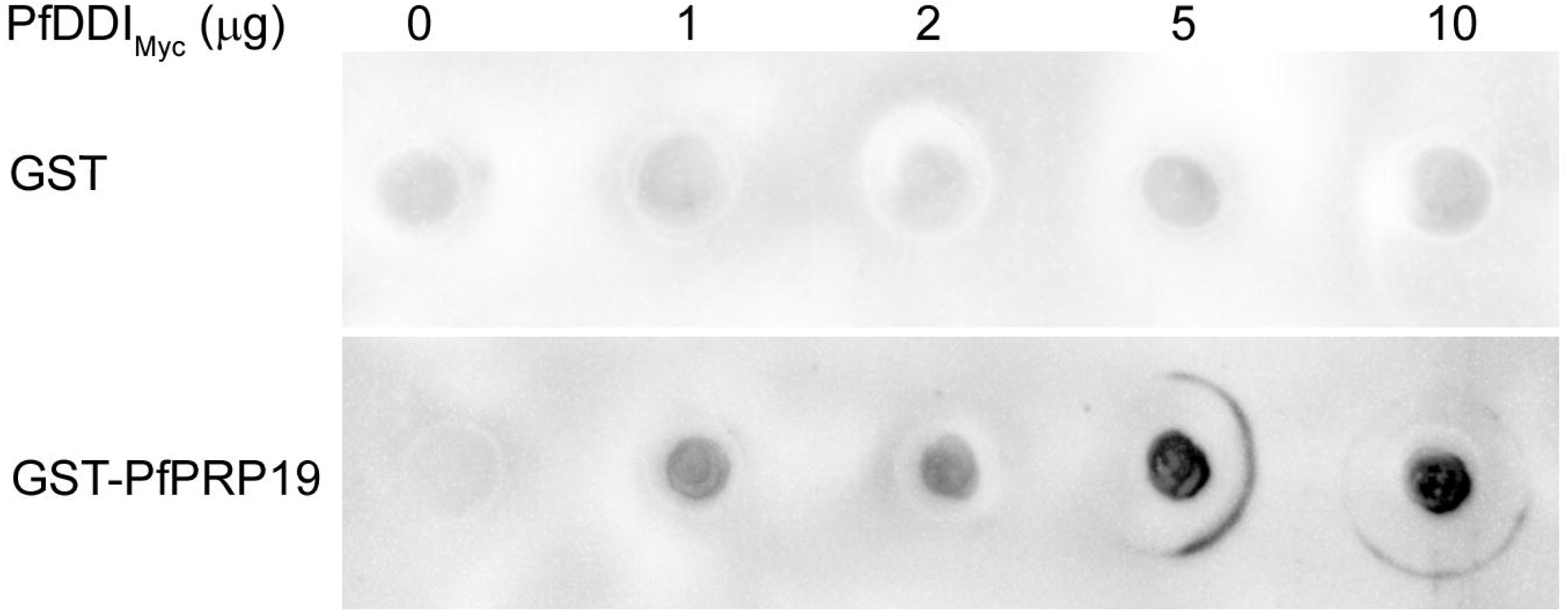
PfDDI1-PfPRP19 overlay assay. The blot containing spots of the indicated amounts of recombinant PfDDI_Myc/His_ (PfDDI_Myc_) was overlaid with recombinant GST or GST/PfPRP19 and probed with anti-GST antibodies.

### 3.9. PfDDI1 reversed the hypersecretion phenotype of ScDDI1 knock-out strain

We generated a *S. cerevisiae* strain lacking the DDI1 gene (ScDDIko). The ScDDIko strain was complemented by expressing wild type PfDDI1 (Sc-wPfDDIki) or catalytic mutant of PfDDI1 (Sc-mPfDDIki) under the native ScDDI1 promoter (Fig. S12A-D). These strains were compared with the wild type strain for growth under various conditions. We did not observe any growth defect in ScDDIko, Sc-wPfDDIki and Sc-mPfDDIki strains as compared to the wild type strain (Fig. S12E). The ScDDIko strain showed hypersecretion phenotype as compared to the wild type, which was reversed to wild type level in Sc-wPfDDIki strain (Fig. S12F), indicating that PfDDI1 is functionally similar to ScDDI1 as a negative regulator of protein secretion. ScDDIko, Sc-wPfDDIki and Sc-mPfDDIki strains were as sensitive as the wild type strain to agents causing DNA damage (hydroxyurea), DNA-protein crosslinks (etoposide), antimalarials (artemisinin and chloroquine) and HIV protease inhibitors (lopinavir and nelfinavir) (data not shown).

## 4. Discussion

DDI1 proteins have been shown to have important roles in several cellular processes, including negative regulation of protein secretion, repair of DPCs and ubiquitin-dependent proteolysis. Several of these functions are associated with the presence of UBL, RVP and UBA domains in DDI1 proteins, including interaction with ubiquitin chains via the UBA and UBL domains, interaction with the proteasome via the UBL domain and cleavage of selected substrates via the RVP domain. LmDDI1 has also been proposed to be the major target of HIV protease inhibitors, which have also been shown to block the development of malaria parasites at multiple stages. However, the *Plasmodium* DDI1 has not been studied yet. We report functional characterization of *Plasmodium* DDI1 using the rodent malaria parasite *P. berghei* and the human malaria parasite *P. falciparum*, which revealed that it is important for parasite development, is a potential target of HIV protease inhibitors and contributes to DPC repair.

PfDDI1 contains the UBL and RVP domains, but lacks the UBA domain. DDI1 homologs are present in all apicomplexa parasites, which vary in size and domain organization. Of the 54 Apicomplexan DDI1 proteins available in the EuPathDB, all contain UBL and RVP domains, and 10 DDI1 proteins also contain the UBA domain. Interestingly, the DDI1 proteins of *B. divergens* and *B. bigemina* contain a putative DUF676 domain in addition to the UBL and RVP domains. Except the Babesiidae parasites, domain organization is conserved among parasites of the same family. Although Apicomplexan DDI1 proteins vary in size from 275 amino acid residues (*G. niphandrodes*) to 1191 amino acid residues (*B. bigemina*), the differences in domain architecture is unlikely due to the differences in protein size. For example, *C. hominis* DDI1 is 384 amino acids long and contains all three domains, whereas *B. bigemina* DDI1 is 1191 amino acids long but contains UBL and RVP domains only. The DUF676 domain is predicted to be a serine esterase and is unique to *B. divergens* and *B. bigemina*, as none of the DDI1 proteins available in the UniProtKB database contains the DUF676 domain. It is a characteristic of the alpha/beta hydrolase superfamily, which contains diverse enzymes (esterases, lipases, proteases, epoxide hydrolases, peroxidases and dehalogenases). The closest related proteins to DUF676 domain of *Babesia* DDI1 are FAM135A proteins. However, there is no report of the characterization of any FAM135A protein. Interestingly, a search of the Apicomplexa genomes using the sequence of DUF676-containing region of *B. bigemina* DDI1 identified DUF676-containing proteins in all parasites except *Plasmodium* and *Piliocolobus*. This suggests that DDI1 and the DUF676-containing protein function together in some pathway. As DUF676 domain is yet to be characterized, it is difficult to predict a role for it or any additional property it might impart to the DDI1 proteins of *B. divergens* and *B. bigemina*. Nonetheless, this domain may confer ubiquitin esterase activity to DDI1 for cleavage of non-lysine ubiquitin chains. Ubiquitin esterase activity is important to cleave the ubiquitin linked to Ser, Cys or Thr residue of the substrate (De Cesare et al., 2021; McClellan et al., 2019). A comparative analysis of the domain organization of DDI1 proteins indicates conservation of UBL and RVP domains in Apicomplexan DDI1 proteins. The diversity in domain architecture might impart species-specific functions to DDI1 proteins, which could enable these parasites to develop in diverse host environments.

Expression of *Plasmodium* DDI1 in asexual erythrocytic, gametocyte, sporozoite and liver stages is in agreement with stage-wise transcription and proteomics data for *Plasmodium* DDI1 proteins, and suggest a major role for it throughout the parasite development. Failure to knock-out the DDI1 gene and drastic growth defects in DDI1 knock-down parasites support a critical role of *Plasmodium* DDI1 during asexual erythrocytic stage development, which is likely to be critical for non-erythrocytic stage development also. The knock-down effect of DDI1 was stronger in *P. berghei* than that in *P. falciparum*, which could be due to the host immune response in the first case. Intrinsic stability of PfDDI1 and degradation efficiency of *Plasmodium* UPS may contribute to the partial knock-down effect in *P. falciparum*. *T. gondii* and *L. major* DDI1 proteins have also been shown to be critical for parasite development (White et al., 2011b; Zhang et al., 2020), which is consistent with our data and support a key role for DDI1 in apicomplexan parasites.

DDI1 knock-down in *P. berghei* not only decreased virulence with self-resolving infection, it also elicited protective immunity to subsequent lethal infections of wild type *P. berghei* ANKA and *P. yoelii* 17XNL. On the other hand, immunization of mice with recombinant PbDDI1 protein did not induce any protection, suggesting that protective immunity could be due to the presence of anti-parasite antibodies and cellular immune responses against the whole parasite. As in our study, prior infection of mice with knock-out parasites (purine nucleoside phosphorylase, plasmepsin 4, plasmepsin-4 + merozoite surface protein-7, nucleoside transporter 1) has also been shown to cause self-limiting infection with development of protective immunity (Aly et al., 2010; Spaccapelo et al., 2011; Spaccapelo et al., 2010; Ting et al., 2008). As has been proposed for knock-out parasites, infection with DDI1 knock-down parasites in the absence of trimethoprim could be a strategy to illicit protective immune response. However, to substantiate the vaccine potential of DDI1 knock-down human malaria parasites, it needs to be investigated whether PfDDDI1 knock-down parasites or knock-down of DDI1 in other *Plasmodium* species would also elicit protective immunity in suitable animal models. As partial knock-down poses a risk of infection, more efficient knock-down strategies should be evaluated to ensure complete knock-down of DDI1 in DDI1-based whole organism vaccines. Furthermore, to achieve stronger attenuation of the whole organism vaccine, DDI1 knock-down can be combined with knock-out of some of the genes that produce attenuated parasites (Aly et al., 2010; Spaccapelo et al., 2011; Spaccapelo et al., 2010; Ting et al., 2008). Attenuated parasites are candidates in efforts to develop a whole organism malaria vaccine and PfDDI1 knock-down parasites could add to the arsenal of whole organism blood stage malaria vaccine candidates.

Knock-down of PfDDI1 significantly increased parasite sensitivity to artemisinin and HIV protease inhibitor lopinavir, which suggested PfDDI1 as a target or role for it in the associated pathways. Artemisinin has been reported to compromise the proteasome function (Bridgford et al., 2018), suggesting that increased susceptibility of PfDDI1 knock-down parasites to artemisinin could be due to the compromised proteasome system. Increased susceptibility to lopinavir upon knock-down of PfDDI1 suggests that it is a target of the HIV protease inhibitor. Of note, the PfKD parasites were not as susceptible to the HIV protease inhibitor nelfinavir and saquinavir as lopinavir. This could be due to differences in inhibitor specificity for PfDDI1, as has been shown for human and *Leishmania* DDI1 proteins previously (White et al., 2011b). Our result is consistent with the previous studies indicating that LmDDI1 is the major target of the anti-leishmanial activity of HIV protease inhibitors (Perteguer et al., 2013; White et al., 2011b). However, it is possible that HIV protease inhibitors also inhibit other targets in malaria parasites, as has been reported for Plasmepsin-V in malaria parasites and multiple pathways in mammalian cells, including inhibition of the chymotryptic activity of proteasome, induction of endoplasmic reticulum stress and cell cycle arrest (Boddey et al., 2010; Brüning et al., 2009; Jiang et al., 2007; Kraus et al., 2008; Russo et al., 2010; Schmidtke et al., 1999).

PfDDI1 knock-down parasites showed increased susceptibility to DNA damaging chemicals and accumulation of DPCs. This is also corroborated by the association of PfDDI1 with chromatin and DPCs, which indicates a role for PfDDI1 in DNA damage response and DPC repair. A role for ScDDI1 in DPC repair has been recently proposed (Serbyn et al., 2020), which is in agreement with our data supporting the involvement of PfDDI1 in DPC repair. Although the mechanism by which DDI1 contributes to the repair of DPCs is not known, DDI1 could help in degradation of DPCs directly owing to its ubiquitin-dependent protease activity and/or by recruiting the repair pathway proteins (Yip et al., 2020).

In contrast to the previously reported protease activity of LmDDI1 (Perteguer et al., 2013), we did not observe degradation of BSA by recombinant PfDDI1 under our experimental settings. Nonetheless, depletion of PfDDI1 caused accumulation of ubiquitinated proteins, which may be a result of the reduced PfDDI1 activity or compromised proteasome function. Ubiquitin-dependent protease activity has been demonstrated for some DDI1 proteins (Dirac-Svejstrup et al., 2020; Yip et al., 2020), and PfDDI1 may be a ubiquitin-dependent protease. PfDDI1 interacted with putative PfPRP19, a homolog of human and *S. cerevisiae* PSO4/PRP19, which is an E3 ubiquitin ligase and has been found to have critical roles in mRNA splicing and multiple DNA repair pathways (Mahajan, 2016; Yin et al., 2012). PfDDI1-PfPRP19 interaction in our study is in line with the involvement of the homologs of these two proteins in DNA damage repair. We speculate that chromatin association of PfDDI1 may be mediated by PfPRP19, as PfPRP19 and other PSO4/PRP19 proteins, but not PfDDI1, contain a DNA-binding domain and a nuclear localization signal.

Expression of PfDDI1 in ScDDI1 knock-out strain reversed the hypersecretion phenotype that has been reported previously (Lustgarten and Gerst, 1999), suggesting that PfDDI1 has a role in regulation of protein secretion. Notably, humans treated with HIV protease inhibitor-based antiretroviral therapy showed reduced incidence of malaria compared to the group treated with non-protease inhibitor-based anti-retroviral therapy (Achan et al., 2012). It is possible that HIV protease inhibitor-treated parasites and DDI1 knock-down *P. berghei* parasites secrete more proteins than wild type parasites, which would act as antigens and elicit stronger immune response than the wild type parasites. ScDDIko, Sc-wPfDDIki and Sc-mPfDDIki strains were similarly sensitive to agents causing DNA damage and DPC, suggesting that deletion of ScDDI1 alone does not alter sensitivity to these agents. Similar effect of these chemicals on ScDDIko, Sc-wPfDDIki and Sc-mPfDDIki strains might be due to dispensability of DDI1 for *S. cerevisiae* growth. Previous studies related to the role of ScDDI1 in DNA damage and DPC repair pathways have been done in combination with additional genes, and knock-out of ScDDI1 alone did not affect sensitivity to some of these chemicals (Serbyn et al., 2020; Svoboda et al., 2019). On the other hand, depletion of PfDDI1 alone increased parasite susceptibility to DNA damaging chemicals with accumulation of DPCs, which indicate a more important role of DDI1 in malaria parasites than that in *S. cerevisiae*.

During the preparation of this manuscript, we also noted two independent studies on *Plasmodium* DDI1. One study is in bioRxiv and another study has been published (Ridewood et al., 2021; Onchieku et al., 2021). The study in bioRxiv, in agreement with our results, demonstrates essentiality of PfDDI1 for asexual stage parasite development and accumulation of ubiquitinated proteins in parasites under conditional PfDDI1 knock-out state. The another study reported that artemisinin inhibits PfDDI1 protease activity, and causes DNA damage and accumulation of PfDDI1 in the nucleus of artemisinin-treated parasites (Onchieku et al., 2021). Consistent with this study as well, our study shows increased chromatin association of PfDDI1 upon DNA damage.

We demonstrate that DDI1 is critical for the malaria parasite development, DDI1 knock- down parasites confer protective immunity and it could be a target of HIV protease inhibitors, which make it a potential candidate for developing a whole organism vaccine and antimalarial drugs. Accumulation of DPCs upon DDI1 knock-down and association of PfDDI1 with chromatin and DPCs support a role for it in DNA damage response and removal of DPC.

## Acknowledgements

NT, MAN, NA and PSS conceived the project, designed experiments, interpreted the results and wrote the manuscript. NT, MAN and NA carried out the majority of experiments. KS, ZR, MAN and RS carried out experiments related to parasite growth, DPC and complementation in yeast. Atul carried out experiments related to generation of PfDDI1 antiserum and localization in *P. falciparum*. AKN carried out mouse experiments. FMAA and AKK carried out experiments related to *P. berghei* sporozoite and liver stages. AKK contributed to manuscript writing. This study and the salaries of MAN, NA and AKN were supported by funding from the Department of Biotechnology, India (BT/COE/34/SP15138/2015). NT and KS are recipients of Research Fellowship from CSIR, India. RS and ZR were supported by fellowships from DBT. The authors would like to acknowledge CCMB central microscopy, Animal and proteomics facilities for their technical support.

## References

1. Achan, J., Kakuru, A., Ikilezi, G., Ruel, T., Clark, T.D., Nsanzabana, C., Charlebois, E., Aweeka, F., Dorsey, G., Rosenthal, P.J., et al. (2012). Antiretroviral agents and prevention of malaria in HIV-infected Ugandan children. N Engl J Med 367, 2110–2118.

2. Al-Nihmi, F.M., Kolli, S.K., Reddy, S.R., Mastan, B.S., Togiri, J., Maruthi, M., Gupta, R., Sijwali, P.S., Mishra, S., and Kumar, K.A. (2017). A Novel and Conserved Plasmodium Sporozoite Membrane Protein SPELD is Required for Maturation of Exo-erythrocytic Forms. Scientific reports 7, 40407.

3. Aly, A.S., Downie, M.J., Mamoun, C.B., and Kappe, S.H. (2010). Subpatent infection with nucleoside transporter 1-deficient Plasmodium blood stage parasites confers sterile protection against lethal malaria in mice. Cell Microbiol 12, 930–938.

4. Batugedara, G., Lu, X.M., Saraf, A., Sardiu, M.E., Cort, A., Abel, S., Prudhomme, J., Washburn, M.P., Florens, L., Bunnik, E.M., et al. (2020). The chromatin bound proteome of the human malaria parasite. Microbial genomics 6.

5. Bertolaet, B.L., Clarke, D.J., Wolff, M., Watson, M.H., Henze, M., Divita, G., and Reed, S.I. (2001a). UBA domains mediate protein-protein interactions between two DNA damage-inducible proteins. Journal of molecular biology 313, 955–963.

6. Bertolaet, B.L., Clarke, D.J., Wolff, M., Watson, M.H., Henze, M., Divita, G., and Reed, S.I. (2001b). UBA domains of DNA damage-inducible proteins interact with ubiquitin. Nature structural biology 8, 417–422.

7. Bhattacharjee, M., Adhikari, N., Sudhakar, R., Rizvi, Z., Das, D., Palanimurugan, R., and Sijwali, P.S. (2020). Characterization of Plasmodium falciparum NEDD8 and identification of cullins as its substrates. Scientific reports 10, 20220.

8. Boddey, J.A., Hodder, A.N., Günther, S., Gilson, P.R., Patsiouras, H., Kapp, E.A., Pearce, J.A., de Koning-Ward, T.F., Simpson, R.J., Crabb, B.S., et al. (2010). An aspartyl protease directs malaria effector proteins to the host cell. Nature 463, 627–631.

9. Bridgford, J.L., Xie, S.C., Cobbold, S.A., Pasaje, C.F.A., Herrmann, S., Yang, T., Gillett, D.L., Dick, L.R., Ralph, S.A., Dogovski, C., et al. (2018). Artemisinin kills malaria parasites by damaging proteins and inhibiting the proteasome. Nature Communications 9, 3801.

10. Brüning, A., Burger, P., Vogel, M., Rahmeh, M., Gingelmaiers, A., Friese, K., Lenhard, M., and Burges, A. (2009). Nelfinavir induces the unfolded protein response in ovarian cancer cells, resulting in ER vacuolization, cell cycle retardation and apoptosis. Cancer Biol Ther 8, 226–232.

11. Ciechanover, A. (2009). Tracing the history of the ubiquitin proteolytic system: the pioneering article. Biochem Biophys Res Commun 387, 1–10.

12. Clarke, D.J., Mondesert, G., Segal, M., Bertolaet, B.L., Jensen, S., Wolff, M., Henze, M., and Reed, S.I. (2001). Dosage suppressors of pds1 implicate ubiquitin-associated domains in checkpoint control. Mol Cell Biol 21, 1997–2007.

13. Czesny, B., Goshu, S., Cook, J.L., and Williamson, K.C. (2009). The Proteasome Inhibitor Epoxomicin Has Potent Plasmodium falciparum Gametocytocidal Activity. Antimicrobial Agents and Chemotherapy 53, 4080–4085.

14. De Cesare, V., Carbajo Lopez, D., Mabbitt, P.D., Fletcher, A.J., Soetens, M., Antico, O., Wood, N.T., and Virdee, S. (2021). Deubiquitinating enzyme amino acid profiling reveals a class of ubiquitin esterases. Proceedings of the National Academy of Sciences 118, e2006947118.

15. Dirac-Svejstrup, A.B., Walker, J., Faull, P., Encheva, V., Akimov, V., Puglia, M., Perkins, D., Kümper, S., Hunjan, S.S., Blagoev, B., et al. (2020). DDI2 Is a Ubiquitin-Directed Endoprotease Responsible for Cleavage of Transcription Factor NRF1. Molecular Cell.

16. Fidock, D.A., and Wellems, T.E. (1997). Transformation with human dihydrofolate reductase renders malaria parasites insensitive to WR99210 but does not affect the intrinsic activity of proguanil. Proc Natl Acad Sci U S A 94, 10931–10936.

17. Finley, D. (2009). Recognition and processing of ubiquitin-protein conjugates by the proteasome. Annu Rev Biochem 78, 477–513.

18. Fivelman, Q.L., McRobert, L., Sharp, S., Taylor, C.J., Saeed, M., Swales, C.A., Sutherland, C.J., and Baker, D.A. (2007). Improved synchronous production of Plasmodium falciparum gametocytes in vitro. Mol Biochem Parasitol 154, 119–123.

19. Gabriely, G., Kama, R., Gelin-Licht, R., and Gerst, J.E. (2008). Different domains of the UBL-UBA ubiquitin receptor, Ddi1/Vsm1, are involved in its multiple cellular roles. Mol Biol Cell 19, 3625–3637.

20. Gantt, S.M., Myung, J.M., Briones, M.R., Li, W.D., Corey, E.J., Omura, S., Nussenzweig, V., and Sinnis, P. (1998). Proteasome inhibitors block development of Plasmodium spp. Antimicrob Agents Chemother 42, 2731–2738.

21. Goldberg, A.L. (2005). Nobel Committee Tags Ubiquitin for Distinction. Neuron 45, 339–344.

22. Govindarajalu, G., Rizvi, Z., Kumar, D., and Sijwali, P.S. (2019). Lyse-Reseal Erythrocytes for Transfection of Plasmodium falciparum. Scientific reports 9, 19952.

23. Hershko, A., and Ciechanover, A. (1982). Mechanisms of intracellular protein breakdown. Annu Rev Biochem 51, 335–364.

24. Hobbs, C.V., Tanaka, T.Q., Muratova, O., Van Vliet, J., Borkowsky, W., Williamson, K.C., and Duffy, P.E. (2013). HIV treatments have malaria gametocyte killing and transmission blocking activity. The Journal of infectious diseases 208, 139–148.

25. Hobbs, C.V., Voza, T., Coppi, A., Kirmse, B., Marsh, K., Borkowsky, W., and Sinnis, P. (2009). HIV protease inhibitors inhibit the development of preerythrocytic-stage plasmodium parasites. The Journal of infectious diseases 199, 134–141.

26. Hu, Q., Klages-Mundt, N., Wang, R., Lynn, E., Kuma Saha, L., Zhang, H., Srivastava, M., Shen, X., Tian, Y., Kim, H., et al. (2020). The ARK Assay Is a Sensitive and Versatile Method for the Global Detection of DNA-Protein Crosslinks. Cell Reports 30, 1235–1245.e1234.

27. Ivantsiv, Y., Kaplun, L., Tzirkin-Goldin, R., Shabek, N., and Raveh, D. (2006). Unique Role for the UbL-UbA Protein Ddi1 in Turnover of SCF^Ufo1^ Complexes. Molecular and Cellular Biology 26, 1579–1588.

28. Janke, C., Magiera, M.M., Rathfelder, N., Taxis, C., Reber, S., Maekawa, H., Moreno-Borchart, A., Doenges, G., Schwob, E., Schiebel, E., et al. (2004). A versatile toolbox for PCR-based tagging of yeast genes: new fluorescent proteins, more markers and promoter substitution cassettes. Yeast (Chichester, England) 21, 947–962.

29. Jiang, W., Mikochik, P.J., Ra, J.H., Lei, H., Flaherty, K.T., Winkler, J.D., and Spitz, F.R. (2007). HIV protease inhibitor nelfinavir inhibits growth of human melanoma cells by induction of cell cycle arrest. Cancer Res 67, 1221–1227.

30. Kaplun, L., Tzirkin, R., Bakhrat, A., Shabek, N., Ivantsiv, Y., and Raveh, D. (2005). The DNA damage- inducible UbL-UbA protein Ddi1 participates in Mec1-mediated degradation of Ho endonuclease. Mol Cell Biol 25, 5355–5362.

31. Khare, S., Nagle, A.S., Biggart, A., Lai, Y.H., Liang, F., Davis, L.C., Barnes, S.W., Mathison, C.J.N., Myburgh, E., Gao, M.-Y., et al. (2016). Proteasome inhibition for treatment of leishmaniasis, Chagas disease and sleeping sickness. Nature 537, 229–233.

32. Koizumi, S., Irie, T., Hirayama, S., Sakurai, Y., Yashiroda, H., Naguro, I., Ichijo, H., Hamazaki, J., and Murata, S. (2016). The aspartyl protease DDI2 activates Nrf1 to compensate for proteasome dysfunction. eLife 5.

33. Kraus, M., Malenke, E., Gogel, J., Müller, H., Rückrich, T., Overkleeft, H., Ovaa, H., Koscielniak, E., Hartmann, J.T., and Driessen, C. (2008). Ritonavir induces endoplasmic reticulum stress and sensitizes sarcoma cells toward bortezomib-induced apoptosis. Mol Cancer Ther 7, 1940–1948.

34. Kumar, S., and Suguna, K. (2018). Crystal structure of the retroviral protease-like domain of a protozoal DNA damage-inducible 1 protein. FEBS open bio 8, 1379–1394.

35. Kustatscher, G., Hégarat, N., Wills, K.L.H., Furlan, C., Bukowski-Wills, J.-C., Hochegger, H., and Rappsilber, J. (2014). Proteomics of a fuzzy organelle: interphase chromatin. The EMBO Journal 33, 648–664.

36. Lambros, C., and Vanderberg, J.P. (1979). Synchronization of Plasmodium falciparum erythrocytic stages in culture. The Journal of parasitology 65, 418–420.

37. Lehrbach, N.J., and Ruvkun, G. (2016). Proteasome dysfunction triggers activation of SKN-1A/Nrf1 by the aspartic protease DDI-1. eLife 5, e17721.

38. Li, H., O’Donoghue, A.J., van der Linden, W.A., Xie, S.C., Yoo, E., Foe, I.T., Tilley, L., Craik, C.S., da Fonseca, P.C.A., and Bogyo, M. (2016). Structure- and function-based design of Plasmodium-selective proteasome inhibitors. Nature 530, 233–236.

39. Lindenthal, C., Weich, N., Chia, Y.S., Heussler, V., and Klinkert, M.Q. (2005). The proteasome inhibitor MLN-273 blocks exoerythrocytic and erythrocytic development of Plasmodium parasites. Parasitology 131, 37–44.

40. Liu, Y., and Xiao, W. (1997). Bidirectional regulation of two DNA-damage-inducible genes, MAG1 and DDI1, from Saccharomyces cerevisiae. Mol Microbiol 23, 777–789.

41. Livneh, I., Cohen-Kaplan, V., Cohen-Rosenzweig, C., Avni, N., and Ciechanover, A. (2016). The life cycle of the 26S proteasome: from birth, through regulation and function, and onto its death. Cell research 26, 869–885.

42. Lustgarten, V., and Gerst, J.E. (1999). Yeast VSM1 encodes a v-SNARE binding protein that may act as a negative regulator of constitutive exocytosis. Mol Cell Biol 19, 4480–4494.

43. Mahajan, K. (2016). hPso4/hPrp19: a critical component of DNA repair and DNA damage checkpoint complexes. Oncogene 35, 2279–2286.

44. Marash, M., and Gerst, J.E. (2003). Phosphorylation of the autoinhibitory domain of the Sso t-SNAREs promotes binding of the Vsm1 SNARE regulator in yeast. Mol Biol Cell 14, 3114–3125.

45. McClellan, A.J., Laugesen, S.H., and Ellgaard, L. (2019). Cellular functions and molecular mechanisms of non-lysine ubiquitination. Open Biol 9, 190147.

46. Morawe, T., Honemann-Capito, M., von Stein, W., and Wodarz, A. Loss of the extraproteasomal ubiquitin receptor Rings lost impairs ring canal growth in Drosophila oogenesis. J Cell Biol 193, 71–80.

47. Navale, R., Atul, Allanki, A.D., and Sijwali, P.S. Characterization of the autophagy marker protein Atg8 reveals atypical features of autophagy in Plasmodium falciparum. PLoS One 9, e113220.

48. Ng, C.L., Fidock, D.A., and Bogyo, M. (2017). Protein Degradation Systems as Antimalarial Therapeutic Targets. Trends Parasitol 33, 731–743.

49. Nowicka, U., Zhang, D., Walker, O., Krutauz, D., Castañeda, C.A., Chaturvedi, A., Chen, T.Y., Reis, N., Glickman, M.H., and Fushman, D. (2015). DNA-damage-inducible 1 protein (Ddi1) contains an uncharacteristic ubiquitin-like domain that binds ubiquitin. Structure (London, England : 1993) 23, 542–557.

50. Nsanzabana, C., and Rosenthal, P.J. (2011a). In vitro activity of antiretroviral drugs against Plasmodium falciparum. Antimicrob Agents Chemother 55, 5073–5077.

51. Nsanzabana, C., and Rosenthal, P.J. (2011b). In vitro activity of antiretroviral drugs against Plasmodium falciparum. Antimicrobial agents and chemotherapy 55, 5073–5077.

52. Onchieku, N.M., Kumari, S., Pandey, R., Sharma, V., Kumar, M., Deshmukh, A., Kaur, I., Mohmmed, A., Gupta, D., Kiboi, D., et al. (2021). Artemisinin Binds and Inhibits the Activity of Plasmodium falciparum Ddi1, a Retroviral Aspartyl Protease. Pathogens 10, 1465.

53. Parikh, S., Gut, J., Istvan, E., Goldberg, D.E., Havlir, D.V., and Rosenthal, P.J. (2005). Antimalarial activity of human immunodeficiency virus type 1 protease inhibitors. Antimicrob Agents Chemother 49, 2983–2985.

54. Perez-Riverol, Y., Csordas, A., Bai, J., Bernal-Llinares, M., Hewapathirana, S., Kundu, D.J., Inuganti, A., Griss, J., Mayer, G., Eisenacher, M., et al. (2019). The PRIDE database and related tools and resources in 2019: improving support for quantification data. Nucleic Acids Res 47, D442–d450.

55. Perteguer, M.J., Gómez-Puertas, P., Cañavate, C., Dagger, F., Gárate, T., and Valdivieso, E. (2013). Ddi1- like protein from Leishmania major is an active aspartyl proteinase. Cell stress & chaperones 18, 171–181.

56. Ramirez, J., Lectez, B., Osinalde, N., Sivá, M., Elu, N., Aloria, K., Procházková, M., Perez, C., Martínez- Hernández, J., Barrio, R., et al. (2018). Quantitative proteomics reveals neuronal ubiquitination of Rngo/Ddi1 and several proteasomal subunits by Ube3a, accounting for the complexity of Angelman syndrome. Human Molecular Genetics 27, 1955–1971.

57. Ridewood, S., Dirac-Svejstrup, A.B., Howell, S., Weston, A., Lehmann, C., Patel, A.P., Collinson, L., Bingham, R., Powell, D., Snijder, A., et al. (2021). The Aspartyl Protease Ddi1 Is Essential for Erythrocyte Invasion by the Malaria Parasite. bioRxiv, 2021.2005.2011.443575.

58. Russo, I., Babbitt, S., Muralidharan, V., Butler, T., Oksman, A., and Goldberg, D.E. (2010). Plasmepsin V licenses Plasmodium proteins for export into the host erythrocyte. Nature 463, 632–636.

59. Saeki, Y. (2017). Ubiquitin recognition by the proteasome. J Biochem 161, 113–124.

60. Schmidtke, G., Holzhütter, H.G., Bogyo, M., Kairies, N., Groll, M., de Giuli, R., Emch, S., and Groettrup, M. (1999). How an inhibitor of the HIV-I protease modulates proteasome activity. J Biol Chem 274, 35734–35740.

61. Serbyn, N., Noireterre, A., Bagdiul, I., Plank, M., Michel, A.H., Loewith, R., Kornmann, B., and Stutz, F. (2020). The Aspartic Protease Ddi1 Contributes to DNA-Protein Crosslink Repair in Yeast. Mol Cell 77, 1066–1079.e1069.

62. Sijwali, P.S., Koo, J., Singh, N., and Rosenthal, P.J. (2006). Gene disruptions demonstrate independent roles for the four falcipain cysteine proteases of Plasmodium falciparum. Mol Biochem Parasitol 150, 96–106.

63. Singhal, N., Atul, Mastan, B.S., Kumar, K.A., and Sijwali, P.S. Genetic ablation of plasmoDJ1, a multi- activity enzyme, attenuates parasite virulence and reduces oocyst production. Biochem J 461, 189–203.

64. Sirkis, R., Gerst, J.E., and Fass, D. (2006). Ddi1, a eukaryotic protein with the retroviral protease fold. Journal of molecular biology 364, 376–387.

65. Sivá, M., Svoboda, M., Veverka, V., Trempe, J.F., Hofmann, K., Kožíšek, M., Hexnerová, R., Sedlák, F., Belza, J., Brynda, J., et al. (2016). Human DNA-Damage-Inducible 2 Protein Is Structurally and Functionally Distinct from Its Yeast Ortholog. Scientific reports 6, 30443.

66. Skinner-Adams, T.S., McCarthy, J.S., Gardiner, D.L., Hilton, P.M., and Andrews, K.T. (2004). Antiretrovirals as antimalarial agents. The Journal of infectious diseases 190, 1998–2000.

67. Spaccapelo, R., Aime, E., Caterbi, S., Arcidiacono, P., Capuccini, B., Di Cristina, M., Dottorini, T., Rende, M., Bistoni, F., and Crisanti, A. (2011). Disruption of plasmepsin-4 and merozoites surface protein-7 genes in Plasmodium berghei induces combined virulence-attenuated phenotype. Scientific reports 1, 39.

68. Spaccapelo, R., Janse, C.J., Caterbi, S., Franke-Fayard, B., Bonilla, J.A., Syphard, L.M., Di Cristina, M., Dottorini, T., Savarino, A., Cassone, A., et al. (2010). Plasmepsin 4-deficient Plasmodium berghei are virulence attenuated and induce protective immunity against experimental malaria. Am J Pathol 176, 205–217.

69. Sudhakar, R., Das, D., Thanumalayan, S., Gorde, S., and Sijwali, P.S. (2021). Plasmodium falciparum Atg18 localizes to the food vacuole via interaction with the multi-drug resistance protein 1 and phosphatidylinositol 3-phosphate. Biochem J 478, 1705–1732.

70. Svoboda, M., Konvalinka, J., Trempe, J.-F., and Saskova, K.G. (2019). The yeast proteases Ddi1 and Wss1 are both involved in the DNA replication stress response (bioRxiv).

71. Ting, L.M., Gissot, M., Coppi, A., Sinnis, P., and Kim, K. (2008). Attenuated Plasmodium yoelii lacking purine nucleoside phosphorylase confer protective immunity. Nat Med 14, 954–958.

72. Trempe, J.F., Brown, N.R., Lowe, E.D., Gordon, C., Campbell, I.D., Noble, M.E., and Endicott, J.A. (2005). Mechanism of Lys48-linked polyubiquitin chain recognition by the Mud1 UBA domain. Embo j 24, 3178–3189.

73. Trempe, J.F., Šašková, K.G., Sivá, M., Ratcliffe, C.D., Veverka, V., Hoegl, A., Ménade, M., Feng, X., Shenker, S., Svoboda, M., et al. (2016). Structural studies of the yeast DNA damage-inducible protein Ddi1 reveal domain architecture of this eukaryotic protein family. Scientific reports 6, 33671.

74. Voloshin, O., Bakhrat, A., Herrmann, S., and Raveh, D. Transfer of Ho endonuclease and Ufo1 to the proteasome by the UbL-UbA shuttle protein, Ddi1, analysed by complex formation in vitro. PLoS One 7, e39210.

75. White, R.E., Dickinson, J.R., Semple, C.A., Powell, D.J., and Berry, C. (2011a). The retroviral proteinase active site and the N-terminus of Ddi1 are required for repression of protein secretion. FEBS Lett 585, 139–142.

76. White, R.E., Powell, D.J., and Berry, C. (2011b). HIV proteinase inhibitors target the Ddi1-like protein of Leishmania parasites. Faseb j 25, 1729–1736.

77. Wlodawer, A., and Erickson, J.W. (1993). Structure-based inhibitors of HIV-1 protease. Annu Rev Biochem 62, 543–585.

78. Wyllie, S., Brand, S., Thomas, M., De Rycker, M., Chung, C.-w., Pena, I., Bingham, R.P., Bueren-Calabuig, J.A., Cantizani, J., Cebrian, D., et al. (2019). Preclinical candidate for the treatment of visceral leishmaniasis that acts through proteasome inhibition. Proceedings of the National Academy of Sciences 116, 9318–9323.

79. Yin, J., Zhu, J.M., and Shen, X.Z. (2012). New insights into pre-mRNA processing factor 19: A multi- faceted protein in humans. Biology of the cell 104, 695–705.

80. Yip, M.C.J., Bodnar, N.O., and Rapoport, T.A. (2020). Ddi1 is a ubiquitin-dependent protease. Proc Natl Acad Sci U S A 117, 7776–7781.

81. Zhang, H., Liu, J., Ying, Z., Li, S., Wu, Y., and Liu, Q. (2020). Toxoplasma gondii UBL-UBA shuttle proteins contribute to the degradation of ubiquitinylated proteins and are important for synchronous cell division and virulence. Faseb j 34, 13711–13725.

