## Supplementary Information for "*Plasmodium* DDI1 is a potential therapeutic target and important chromatin-associated protein"

**Note: Supplementary data associated with this article**

#### Materials and methods

##### *1. Production of recombinant proteins*

The complete PfDDI1 coding region was amplified from *P. falciparum* genomic DNA using primers PfDdi-expF/PfDdi-expR, cloned into the pQE-30 plasmid at BamHI-HindIII to obtain pQE30-PfDDI1, and transformed into M15(pREP4) *E. coli* cells. pQE30-PfDDI1 would express PfDDI1 as an N-terminal His-tagged protein, facilitating purification of the protein by Ni-NTA affinity chromatography. For purification of $_{\text{His}}$PfDDI1, the IPTG-induced cell pellet was resuspended in urea buffer (8 M Urea, 10 mM Tris, 250 mM NaCl, pH 8.0), incubated for 30 minutes at room temperature, and the lysate was sonicated for 5 minutes (9 seconds on-off cycle at 20% amplitude) using SONICS Vibra-Cell ultrasonic processor. The lysate was centrifuged at 18000g for 30 minutes at 4$^{o}$C, the supernatant was supplemented with imidazole (10 mM) and incubated with Ni-NTA resin (0.2 ml resin/5g of initial cell pellet) at room temperature for 30 minutes. The resin was transferred to a column and washed with urea buffer containing imidazole (20-50 mM). The bound protein was eluted with elution buffer (250 mM imidazole in urea buffer) and resolved in a 12% SDS-PAGE. Elution fractions enriched with His-PfDDI1 were pooled and dialyzed against the refolding buffer-1 (10 mM Tris, 1 mM GSH, 0.5 mM GSSG, 50 mM NaCl, 10% glycerol, pH 8.0) for 12 hours at 4$^{o}$C, followed by against the refolding buffer-2 (10 mM Tris, 50 mM NaCl, 0.5 mM DTT, pH-8.0) with 2 changes at an interval of 3 hours. The refolded protein was concentrated using a 3-kDa cut off Amicon Ultra-15, estimated by Bradford assay, and stored at -80°C till further use.

A codon optimized synthetic version of PfDDI1 gene with C-terminal Myc-tag (synPfDDI1$_{Myc}$) was purchased from Life Technologies (pMA-DDI), and used as a template for amplification of PfDDI1$_{Myc}$ coding region using PRPsyn-Fchis/PRPsyn-Rchis primers. The PCR fragment was cloned into pET26B at NdeI-XhoI site to obtain pET26B-PfDDI1$_{Myc}$ plasmid, and transformed into the Rosetta-gami 2(DE3) *E. coli* cells for expression. The recombinant protein would be expressed with a C-terminal His-tag (PfDDI1$_{Myc/His}$), enabling purification by Ni-NTA chromatography. A pET26B-PfDDI1$_{Myc}$ expression clone was grown in LB broth and induced with IPTG (0.5 mM final) at an $OD_{600}$ of 0.6 at 20ºC for 4 hours with shaking at 225 rpm. The induced cell pellet was resuspended in lysis buffer (20 mM Tris, 500 mM NaCl, 20 mM Imidazole, 2 mM β-mercaptoethanol, pH 8; lysozyme at 1 mg/ml; 5 ml buffer/g pellet), incubated for 30 min at 4°C and sonicated (30% amp with 5 sec on/off for 30 min) using SONICS Vibra-Cell ultrasonic processor. The lysate was centrifuged at 39191g for 1 hour at 4°C, the supernatant was applied on the HisTrap HP column (GE healthcare), washed with lysis buffer, and the bound proteins were eluted (20 mM Tris, 500 mM NaCl, 0-250 mM Imidazole, 2 mM β-mercaptoethanol, pH 8). The elution fractions were resolved in 12% SDS-PAGE and the fractions with high purity of the recombinant protein were concentrated using a 50 kDa cutoff Amicon Ultra concentrator. The concentrated protein was further purified by gel filtration chromatography (Hiload superde x 200 16/600, GE healthcare) with buffer (20 mM Tris, 150 mM NaCl, 2 mM β-mercaptoethanol, pH 7.5). The elution fractions were resolved on 12% SDS-PAGE to check for purity, fractions containing the pure protein were pooled, concentrated using a 10 kDa cut off Amicon Ultra-15, quantified using the BCA method (Thermo Fisher Scientific), and used immediately or stored at -80°C till further use. To determine the size of PfDDI1$_{Myc/His}$, reference protein size markers and gel filtration purified PfDDI$_{Myc/His}$ were loaded on the column (HiLoad Superdex 200 16/600, GE healthcare) with buffer (20 mM Tris, 150 mM NaCl, 2 mM β-mercaptoethanol, pH 7.5). The elution volumes of reference protein size markers were plotted against their sizes, and the plot was used to determine the size of $PfDDI_{Myc/His}$.

Wild type *P. berghei* DDI1 (PbDDI1) was amplified from *P. berghei* gDNA using PbDdiexpF/PbDdiexpRm primers (PbDdiexpRm encodes for an in-frame Myc-tag). The PCR fragment was cloned into pRSET-A at HindIII-XhoI site to obtain $pRSETA\text{-}wPbDDI1_{Myc}$ expression plasmid. A catalytic mutant of PbDDI1 (D268A) was generated by overlap PCR. Briefly, the 5'- and 3'-fragments of PbDDI1 coding region were amplified from the pRSETA-wtPbPRPmyc plasmid using PbDdiexpF/Pbddimut-R and PbDdimut-F/PbDdiexpRm primer sets, respectively. The two fragments overlapped around the mutation site, facilitating recombination by PCR using PbDdiexpF/PbDdiexpRm primers. The recombined fragment was cloned into pRSET-A at BamHI-HindIII sites to obtain $pRSETA\text{-}mPbDDI1_{Myc}$ expression plasmid. The expression plasmids were transformed into *BL21-CodonPlus* (*DE3*) *E. coli* cells, which would express recombinant proteins with N-terminal His-tag, thereby enabling purification by Ni-NTA affinity chromatography. The wild type ($wPbDDI_{Myc}$) and mutant ($mPbDDI_{Myc}$) recombinant proteins were purified from the IPTG-induced cells of respective expression clones under native conditions. Briefly, the expression clones were grown in LB broth and expression was induced with IPTG (1 mM final) at an $OD_{600}$ of 0.6 at 20ºC for 4 hours with shaking at 225 rpm. The cell pellet was resuspended in native buffer (50 mM $NaH_2PO_4$, 100 mM NaCl, pH 8.0; lysozyme at 1 mg/ml; 5 ml buffer/g pellet), incubated in ice for 30 min and sonicated for 4 min (9 second pulses at 20% amplitude) using SONICS Vibra-Cell ultrasonic processor. The lysate was centrifuged at 25,000g for 30 min, the supernatant was supplemented with imidazole (10 mM) and incubated with Ni-NTA agarose resin (0.4 ml slurry/g weight of the initial cell pellet) for 30-45 min at 4°C.

The resin suspension was transferred to a column, washed with wash buffer (native buffer with 20-50 mM imidazole) and the bound proteins were eluted with elution buffer (native buffer with 250 mM imidazole). The elution fractions were resolved on 12% SDS-PAGE. The fractions enriched with the recombinant protein were pooled and dialyzed against the storage buffer (20 mM Tris-Cl, 50 mM NaCl, pH 8.0) using a 10 kDa cut off dialysis tubing at 4°C. The dialyzed protein was concentrated using a 10 kDa cut off Amicon Ultra-15, quantified using the BCA method (Thermo Fisher Scientific), and stored at -80°C.

***2. Protease assay***

The assay samples containing 2 µg BSA and 1 µg recombinant $PfDDI_{Myc/His}$ in 30 µl buffer (20 mM Tris, 50 mM NaCl, 2 mM β-mercaptoethanol, pH 7.0 or 7.5) were incubated at 37°C. The reaction was stopped by adding 6 µl of 4 × SDS-PAGE sample buffer (1 × buffer contains 50 mM Tris-HCl, 20% glycerol, 2% SDS, 1% β-mercaptoethanol, 0.01% bromophenol blue, pH 6.8) to assay samples at different time points (0, 1, 2.5, 5 and 10 hours). Control assay samples contained the same amount of BSA or recombinant $PfDDI_{Myc/His}$, which were incubated at 37°C for 10 hours and stopped by adding 6 µl of 4 × SDS-PAGE sample buffer to the sample. The reaction and control samples were resolved in 12% SDS-PAGE and stained with coomassie brilliant blue.

***3. Construction of transfection plasmids***

The PbDDI1 gene was targeted for knock-out and knock-down using double cross-over homologous recombination approach. The 5'-UTR (flank 1) and 3’-UTR (flank 2) of PbDDI1 were amplified from *P. berghei* genomic DNA using PbDdi1-Fl1F/ PbDdi1-Fl1R and PbDdi1-Fl2F/PbDdi1-Fl2R primer sets, respectively. The flank 1 and flank 2 were cloned into the HB-DJ1KO plasmid at NotI-KpnI and AvrII-KasI sites, respectively, to obtain HB-PbDDI-(FL1+FL2) plasmid. The GFP coding sequence was excised from pGT-GFPbsc with KpnI-XhoI and subcloned into the similarly digested HB-PbDDI-(FL1+FL2) to obtain HB-pbDDIKO plasmid. For construction of knock-down plasmid, the $PfDDI_{Myc}$ coding sequence was amplified from the *P. falciparum* genomic DNA using DDiexp-F/DDimyc Rep-R primers, digested with KpnI-XhoI and subcloned into the similarly digested HB-PbDDIKO plasmid in place of GFP to obtain HB-PfDDIKI plasmid. The *E. coli* mutant DHFR coding sequence with HA-tag ($cDD_{HA}$) was amplified from the pPM2GDBvm plasmid (a kind gift from Dr. Praveen Balabaskaran Nina) using cDD-F/cDD-R primers, and cloned into the pGT-GFPbsc plasmid at KpnI/XhoI sites to obtain pGT-$cDD_{HA}$ plasmid. The $PfDDI_{Myc}$ coding sequence was amplified from HB-PfDDIKI plasmid using the PfDdi-reF/PfDdi-cDDR primers and cloned into the pGT-$cDD_{HA}$ plasmid at BglII-BamHI site to obtain the pGT-PfDDI-$cDD_{HA}$ plasmid. The pGT-$PfDDI_{Myc}$/$cDD_{HA}$ plasmid was digested with BglII-XhoI to release the $PfDDI_{Myc}$/$cDD_{HA}$ insert, which was cloned into the similarly digested HB-DDKI plasmid to obtain HB-pbDDIKD vector. All the flanks and coding regions were sequenced to ensure that they were free of undesired mutations, and the presence of different regions was confirmed by digestion with region-specific restriction enzymes. The HB-pbDDIKO and HB-pbDDIKD plasmids were purified using the NucleoBond® Xtra Midi plasmid DNA purification kit (MACHEREY-NAGEL), linearized with NotI-KasI, gel purified to obtain pbDDIKO and pbDDIKD transfection constructs, and used for transfection of *P. berghei.*

The flank 1 of HB-pbDDIKD was excised with NotI-BglII, the ends of plasmid backbone were filled and ligated. The flank 2 of this plasmid was excised with AvrII-KasI, and the plasmid backbone was ligated with the similarly digested PfDDI1-3'UTR, which was amplified from *P. falciparum* genomic DNA using PfDDi3'U-F/PfDDi3'U-R primers. The final plasmid was called HF-pfDDIKD, which contains $PfDDI1_{Myc}/cDD_{HA}$ coding region as flank1 and PfDDI1-3'UTR as flank2. HF-pfDDIKD was digested with region-specific restriction enzymes to ensure the presence of different regions, purified using the NucleoBond® Midi plasmid DNA purification kit (MACHEREY-NAGEL), and used for transfection of *P. falciparum*.

***4. Measurement of antibody titers***

The sera collected from recovered mice and immunized mice were tested for antibody titers against the soluble *P. berghei* ANKA parasite extract and recombinant $mPbDDI_{Myc}$ by ELISA, respectively. Recombinant $mPbDDI_{Myc}$ was coated on the wells of 96-well MicroWell™ MaxiSorp™ flat bottom plates (1 µg/well in 100 µl of 0.2 M bicarbonate buffer, pH 9.2) at 4°C for overnight. The plate was washed with PBS-T (PBS with 0.05% Tween 20) to remove the unbound protein, blocked (blocking solution: 2% BSA in PBS-T), and incubated with serial two-fold dilutions (in blocking solution) of sera (immune sera, adjuvant control sera and pre-immune sera) for 2 hours at room temperature. The plate was washed with PBS-T, incubated with HRP-conjugated horse anti-mouse IgG (at 1/2000 dilution in blocking solution) for 1 hour at room temperature, washed with PBS-T, and incubated with 100 µl of TMB ELISA substrate for 30 min. The reaction was stopped with 1N HCl and absorbance was measured at 450 nm using the BioTecPowerWave XS2 spectrophotometer. The absorbance values were adjusted for the background absorbance and absorbance values of the pre-immune sera. The data was plotted against the sera dilutions using the GraphPad Prism software. For antibody titers in the sera of recovered mice, a pellet of the wild type *P. berghei* ANKA erythrocytic parasites was resuspended in 5× pellet volume of the lysis buffer (PBS with 0.1% NP-40), subjected to 5 cycles of freeze-thaw, followed by 5 cycles of passage through a 27 G needle. The lysate was incubated in ice for 30 minutes and centrifuged at 25000g for 30 minutes at 4ºC. The supernatant that contains the soluble parasite extract was separated and the protein amount was estimated by BCA. The soluble parasite extract was coated on the wells of the 96-well MicroWell™ MaxiSorp™ flat bottom plates (2 µg protein/well in 0.2 M bicarbonate buffer, pH 9.2) and assessed for reactivity with the pre-challenge sera of recovered mice as described for recombinant $mPbDDI_{Myc}$.

##### *5. Immunoprecipitation of PfDDI1 and mass spectrometry*

Asynchronous cultures of wild type *P. falciparum* D10 and PfKD (with 10 µM trimethoprim) parasites were grown and parasites were isolated at 10-15% parasitemia as has been described in the parasite culture section. The parasite pellets were resuspended in 10× pellet volume of the lysis buffer (10 mM Tris-Cl, 10 mM HEPES, 150 mm NaCl, 0.5 mM EDTA, 0.5% TritonX-100, pH 7.5; MERCK protease inhibitor cocktail), and subjected to 2 cycles of freeze-thaw, followed by passing through the 26.5 G needle. The lysates were centrifuged at 20000g for 30 min at 4°C, and the supernatant was transferred into a fresh tube. The pellet was re-extracted with 3× pellet volume of the lysis buffer as described above, and the supernatant was combined with the first supernatant. The supernatant was incubated with Myc-Trap magnetic beads (ChromoTek, 5 µl slurry/mg protein) for 2 hours at 4°C with gentle mixing. The flow through was removed and the beads were washed (10 mM Tris-Cl, 10 mM HEPES, 150 mm NaCl, 0.5 mM EDTA, pH 7.5; protease inhibitor cocktail). The beads were boiled in 100 µl of 2× SDS-PAGE sample buffer for 15 min, and the eluate was processed for western blotting and mass spectrometry. 20 µl of the eluate along with appropriate controls (input, flow through and washes) was assessed for the presence of $PfDDI_{Myc}/cDD_{HA}$ by western blotting using mouse anti-Myc antibody, followed by appropriate secondary antibody as described in the western blotting section.

80 µl of the eluate was run on a 10% SDS-PAGE gel until the protein ladder completely entered into the resolving gel. The protein band was excised, cut into small pieces, treated with trypsin, peptides were extracted, vacuum dried, and resuspended in 11 µl of 2% formic acid as has been described in detail previously (Bhattacharjee et al., 2020; Sudhakar et al., 2021). 10 µl of the peptide sample was run on the Q-Exactive HF (Thermo Fischer Scientific) for HCD mode fragmentation and LC-MS/MS analysis. The raw data files were acquired on the proteome discoverer v2.2 (Thermo Fischer Scientific), analysed and searched against the Uniprot databases of *P. falciparum* 3D7 using the HTSequest algorithm. The analysis parameters included trypsin specificity, maximum two missed cleavages and some variable modifications (carbamidomethylation of cysteine, oxidation of methionine, deamidation of asparagine/glutamine). Other parameters included: precursor tolerance of 5 ppm, fragmentation tolerance of 0.05 Da and 1% peptide FDR threshold. The mass spectrometry proteomics data have been deposited to the ProteomeXchange Consortium via the PRIDE partner repository with the dataset identifier PXD030157 (Perez-Riverol et al., 2019). All other data is included in the manuscript and any related data will be made available upon request. The protein hits from the PfKD samples were compared with those from the wild type parasites. Proteins exclusively present in the PfKD sample with at least 5 times higher peptide spectrum matches (PSMs) than in the wild type sample were selected. The selected proteins were considered if present in at least 3 biological replicates with a minimum of 1 unique peptide.

***6. Complementation of S. cerevisiae DDI1***

The ScDDI1 coding region of *S. cerevisiae* BY4741strain was replaced with kanamycin cassette (for knock-out) or the GFP-PfDDI1$_{Myc}$ cassette containing the wild type or mutant PfDDI1 coding sequence (knock-in). The kanamycin cassette was amplified from pFA6a-kanMX6 plasmid using ScDDi-Fko/PfDDi-Rki primers. For wild type PfDDI transfection cassette, the pGEX-synPfDDI$_{Myc}$ plasmid was digested with XhoI, ends were filled with Klenow, and again digested with BamHI to excise the PfDDI1$_{Myc}$ region, which was cloned into the pFA6a-GFP plasmid at BamHI/SmaI site to obtain the pFA6-GFP/PfDDI$_{Myc}$ plasmid. The plasmid was used as a template for PCR amplification of the wild type PfDDI1 transfection cassette (GFP/wPfDDI$_{Myc}$) using PfDDi-Fki/PfDDi-Rki primers. The mutant PfDDI transfection cassette (GFP/wPfDDI$_{Myc}$) was generated by recombination PCR using the pFA6-GFP/PfDDI$_{Myc}$ plasmid as a template with PfDDi-Fki/PRPsynD-A-R and PRPsynD-A-F/PfDDi-Rki primer sets as described for recombinant mPbDDI1$_{Myc}$ in the production of recombinant proteins section. The kanamycin cassette, GFP/wPfDDI$_{Myc}$ and GFP/mPfDDI$_{Myc}$ PCR products were gel purified and transformed into the BY4741 cells by lithium acetate method (Janke et al., 2004). Briefly, the BY4741 strain was grown in complete YPD medium at 25ºC and 250 rpm till $OD_{600}$ of 1.0. The culture was centrifuged (3000g for 5 minutes at 25ºC), the cell pellet was washed with autoclaved Milli-Q water, followed by with lithium acetate/Tris EDTA (LTE) buffer (100 mM lithium acetate, 10 mM Tris-Cl, 4 mM EDTA, pH 7.5). The cell pellet was resuspended in 100 µl of transformation mix (LTE, 34.7% PEG, 10 µg of calf thymus DNA, 4 µg of desired PCR product), incubated at 30°C for one hour, followed by at 42ºC for 15 minutes, and then in ice for 10 minutes. The transformation reaction was centrifuged at 3300g for 2 minutes at room temperature, the cell pellet was resuspended in

YPD medium, and grown at 25°C with shaking at 250 rpm for overnight. The culture was centrifuged at 3000g for 5 minutes at room temperature, the cell pellet was washed with autoclaved Milli-Q water, resuspended in autoclaved Milli-Q water, spread on YPD agarose plates containing G418 (350 µg/ml), and the plates were incubated at 25ºC. The resistant colonies along with the BY4741strain as a control were grown overnight in 5 ml of YPD medium at 25ºC and 250 rpm. The cultures were harvested at 3000g for 5 minutes at 25ºC, and the cell pellets were used for western blot analysis and gDNA isolation as has been described earlier (Bhattacharjee et al., 2020). The integration of transfection cassette was checked by PCR using primers specific for the complete cassette (Sc/PfDDi-5con, Sc/PfDDi-3con), 5'-integration (Sc/PfDDi-5con and PfDDIUbl-R) and 3'-integration (RVP/UBA-F and Sc/pfDDi 3con) loci. The lysates of complemented (ΔScddi :: wPfDDI1$_{Myc}$ and ΔScDDI1 :: mPfDDI1$_{Myc}$) strains and BY4741 as a control were evaluated for expression of GFP/wPfDDI$_{Myc}$ or GFP/mPfDDI$_{Myc}$ proteins by western blotting using anti-GFP antibodies and anti-triose phosphate isomerase antibodies, followed by appropriate secondary antibodies as has been described in western blotting section.

The BY4741, ΔScDDI1 and complemented (ΔScddi :: wPfDDI1$_{Myc}$ and ΔScDDI1 :: mPfDDI1$_{Myc}$) strains were compared for growth on YPD agarose plate containing various inhibitors. In brief, overnight cultures were initiated with single colonies in complete YPD with G418 (for complemented strains) at 25ºC and 250 rpm. The overnight culture was added to fresh complete YPD medium (at 1%) without G418, and grown till the $OD_{600}$ of 1-2. The cultures were normalized to $OD_{600}$ of 1.0, serial 10-fold dilutions were made in YPD, and 5 µl of each dilution of all the strains was spotted on YPD agarose plates without or with a test chemical (0.005% MMS, 100 µg/ml etoposide, 100 mM hydroxyurea, 0.1% glyoxal, 100 µM artemisnin, 100 µM chloroquine, 100 μM pepstatin A, 100 μM bestatin, 100 μM lopinavir and 100 μM nelfinavir). The plates were incubated at 30°C for 2-3 days.

The BY4741, ΔScDDI1 and complemented (ΔScddi :: wPfDDI1$_{Myc}$) strains were compared for protein secretion into the culture medium. For each strain, a single colony was inoculated into 5 ml SD medium, incubated at 30ºC for 48 hours with shaking at 180 rpm, and the culture was centrifuged at 3000g for 15 minutes at 25ºC. The cell pellet was dried at 100°C for overnight and the pellet weight was measured. The supernatant was dialysed against PBS at 4°C using a 3.5 kDa cut off dialysis membrane and the protein amount was estimated using BCA method. The total protein amount present in the supernatant was normalized with dry weight of the cell pellet from the same culture and plotted as μg/mg dry weight of the pellet using GraphPad Prism.

**Table S1.** Primers used in the study.

| Primer | Sequence (5'-3') |
|---|---|
| PfDdi-expF | ATGGGATCCGTTTTTATTACAATATCAGACGAT |
| PfDdi-expR | AATTAAAGCTTATAAATCATTGTTTGCATCAATGTC |
| PbDdiexpF | TATGGGATCCGTGTTCATAACAATATCTGATGAT |
| PbDdiexpRm | TAATAAGCTTCTACAGGTCTTCTTCAGAAATCAGCTTTTGTTCTAGTGTGTCTAAGTTTATGTTTTC |
| Pbddimut-R | GTTTGTGCCCCTGAAGCGACAAATGCATG |
| PbDdimut-F | CATGCATTTGTCGCTTCAGGGGCACAAAC |
| PbDdiexpRm | TAATAAGCTTCTACAGGTCTTCTTCAGAAATCAGCTTTTGTTCTAGTGTGTCTAAGTTTATGTTTTC |
| PRPsyn-Fchis | ATCGTACATATGGTGTTTATTACCATTAGCG |
| PRPsyn-Rchis | GAATTACTCGAGCAGATCCTCTTCGCTAATCAG |
| PbDdi1-Fl1F | AATTGCGGCCGCAAAAATTTCAATCATAGAGTATATATAC |
| PbDdi1-Fl1R | ATATGGTACCAGATCTCATTTTGGTCAAATCTTGTGATATTC |
| PbDdi1-Fl2F | ATAGCCTAGGGGCCCATAAAAATATACTATCCG |
| PbDdi1-Fl2R | AATTGGCGCCGGTTATTATTATCTTAAGTTTGTGACCATAC |
| DDiexp-F | ATGGGATCCGTTTTTATTACAATATCAGACGAT |
| DDimyc Rep-R | TAATCTCGAGTTACAGGTCTTCTTCAGAAATAAGCTTTTGTTCTAAATCATTGTTTGCATCAATGTC |
| cDD-F | AATG*GGATCC*GGAGGCGGTGGAGGAATCAGTCTGATTGCGGCGTTA |
| cDD-R | CTAACTCGAGTCAAGCGTAATCTGGAACATCGTATGG |
| PfDdi-reF | AATGAGATCTGTTTTTATTACAATATCAGACGAT |
| PfDdi-cDDR | CCAAGGATCCCAGGTCTTCTTCAGAAATAAGCTTTTG |
| CON5'-F | GACAACATGATGCCCTTATGGAATATG |
| Pvac-R | CGGTCTGCAGCCCTGGAAAACAGGCGAT |
| Hrp2-F | ATCAGGCGCCATATCGAATTCCCGCGATCGATCCTAGGAACATATGTTTAAAAGAAAAATTTAAGATTTACATGATTAGG |
| CON3'-R | CACTAAAATTCATAAGTGAAAAGTTGTCAC |
| Pbspec-F | CACATCAGCTACTAATGTTAGTACATCC |
| PFspec-F | TCTCTTAAATAATCCAGCTTTCAAAACA |
| α6FL2–F | ATAACCTAGGAGACGTGAAAGTTATATCGACC |
| α6FL2-R | AATTGATATCGAATTCCAAACACAAGCAAGCACTTGTGC |
| PfDDi3'U-F | TTATAGTCGACCGTAAATATTATTATCACTATCAATATATATGTAC |
| PfDDi3'U-R | AGCTTCTGCAGTTATTATTCATCTATTAACATAAGGGAAA |
| DDi-con5U | CTAACATTTTAACATTTTAATAATTAAGCATTAAAAATG |
| cDD-R | CTAACTGAGTCAAGGGTAATCTGGAACTCGTATGG |
| HRP2seq-F | CTTTTACAATATGAACATAAAGTACAAC |
| DDi3'int-R | CAACAGAATGTGCTCAGATCATAC |
| PfVMP1-Fep | TAGCTTAGATCTATGGATTCATATGAAGCTAAGAAGAAG |
| PfVMP1-R | CATTAGGTACCTTTTTTTATTTTTCTTGGTGTCATTTCGTTC |
| PfPPF19-Fexp | TAACGCAGATCTGGATCCATGTCAATAATATGCACCATAAGTGGC |
| PfPPF19-Rexp | GACTCTCGAGTTAGGTACCATTCCAAAGTTTTATTGTTTTATCCATTGA |
| ScDDi-Fko | GTATACTTAACGTAGTTAACAAAGTACATACCAAACATAACAGCAAAAATATACGTAAAGAGCGACATGGAGGCCCAGAATA |
| PfDDi-Rki | ACTTATCTATTTGTGTTATGGGCTACATACGTAGAGGCCGATCACAATATCAGTGGTTGCTCACAGTATAGCGACCAGCA |
| PRPsynD-A-F | GTGCATGCATTTGTTGCCAGCGGTGCACAGAGC |
| PfDDi-Rki | GCTCTGTGCACCGCTGGCAACAAATGCATGCAC |
| PfDDi-Fki | GTATACTTAACGTAGTTAACAAAGTACATACCAAACATAACAGCAAAAATATACGTAAAGATGAGTAAAGGAGAAGAACTTTTCAC |
| Sc/PfDDi-5con, | ATTCATCGCCACCGAAAAGATA |
| Sc/PfDDi-3con | TATACTTAACAGAAGTACAATC |
| RVP/UBA-F | TAAGTCAGCATATGGTTGTGTTCATGCTGTATATTCCGGTGGAA |
| PfDDIUbl-R | TACACTCACTCGAGTTAGTGATGGTGATGGTGATGTGCGCTGATTTTTTTGCGAACAAA |

**Table S2. Proteins identified in mass spectrometry analysis of the PfDDI1 immunoprecipitate.** Shown are the score (Sc), coverage (Co), and number of unique peptides (UP) for each protein. The proteins shown were consistently identified in three independent biological replicates. The predicted pathway is based on homologs in other organisms.

| Accession ID | Protein | Experiment 1 | | | Experiment 2 | | | Experiment 3 | | | Pathway |
|---|---|---|---|---|---|---|---|---|---|---|---|
| | | Sc | Co | UP | Sc | Co | UP | Sc | Co | UP | |
| Q8IM03 (PF3D7_1409300) | DNA damage-inducible protein 1 | 205.1 | 35.6 | 21 | 169.4 | 31.0 | 21 | 835.1 | 51.9 | 34 | Bait protein |
| Q8I2Z8 (PF3D7_0915400) | Probable ATP-dependent 6-phosphofructokinase | 144.1 | 24.6 | 26 | 12.1 | 2.7 | 4 | 15.8 | 5.1 | 6 | Glycolysis pathway |
| Q8I3T1 (PF3D7_0517700) | Eukaryotic translation initiation factor 3 subunit B | 123.6 | 36.5 | 22 | 9.9 | 4.3 | 3 | 5.7 | 1.4 | 1 | Protein synthesis |
| O77325 (PF3D7_0308600) | Pre-mRNA-processing factor 19 | 60.8 | 20.3 | 8 | 10.2 | 6.4 | 3 | 12.1 | 7.7 | 2 | DNA repair, mRNA splicing |
| Q8IKH8 (PF3D7_1465900) | 40S ribosomal protein S3 | 46.4 | 41.2 | 8 | 20.7 | 27.6 | 7 | 73.6 | 38.0 | 9 | Translation |
| O00806 (PF3D7_0309600) | 60S acidic ribosomal protein P2 | 42.2 | 49.1 | 5 | 40.7 | 52.7 | 4 | 76.2 | 52.7 | 4 | Translation |
| Q8IIA2 (PF3D7_1126200) | 40S ribosomal protein S18 | 33.3 | 37.2 | 6 | 6.0 | 19.9 | 2 | 33.5 | 35.3 | 4 | Translation |
| Q8I4Y9 (PF3D7_1244100) | N-alpha-acetyltransferase 15, NatA auxiliary subunit | 31.8 | 8.2 | 8 | 3.4 | 1.2 | 1 | 1.7 | 1.8 | 1 | - |
| Q8IAX6 (PF3D7_0814000) | 60S ribosomal protein L13 | 30.6 | 24.7 | 6 | 16.3 | 19.5 | 4 | 51.8 | 22.8 | 6 | Translation |
| Q8IAL3 (PF3D7_0801500) | Nucleolar protein 10 | 28.8 | 16.0 | 7 | 31.4 | 12.8 | 6 | 35.1 | 17.5 | 8 | - |
| Q8ILK3 (PF3D7_1426000) | 60S ribosomal protein L21 | 26.1 | 38.5 | 5 | 4.1 | 10.6 | 1 | 44.0 | 35.4 | 6 | Translation |
| Q8IM23 (PF3D7_1407100) | Fibrillarin | 25.1 | 22.6 | 7 | 11.9 | 12.3 | 4 | 40.7 | 31.8 | 8 | rRNA processing |
| C0H5C2 (PF3D7_1317800) | 40S ribosomal protein S19 | 25.1 | 22.8 | 3 | 7.9 | 18.6 | 2 | 44.2 | 36.6 | 5 | Translation |
| Q8IHT2 (PF3D7_1143400) | Translation initiation factor eIF-4C | 24.5 | 28.4 | 4 | 8.2 | 11.0 | 2 | 11.0 | 16.8 | 2 | Translation |
| Q8IKS3 (PF3D7_1455500) | AP-1 complex subunit gamma | 22.9 | 7.2 | 6 | 2.2 | 1.3 | 1 | 14.8 | 4.3 | 3 | - |
| Q8I0W8 (PF3D7_0513600 ) | Deoxyribodipyrimidine photo-lyase | 22.2 | 6.7 | 6 | 9.0 | 3.2 | 3 | 8.9 | 1.8 | 2 | - |
| Q7K6A0 (PF3D7_0934800) | cAMP-dependent protein kinase catalytic subunit | 21.3 | 24.0 | 8 | 3.7 | 7.3 | 3 | 6.6 | 7.6 | 2 | Signal transduction |
| Q8I5C4 (PF3D7_1229500) | T-complex protein 1 subunit gamma | 21.1 | 15.5 | 8 | 22.8 | 12.9 | 6 | 41.1 | 14.9 | 7 | Protein folding |
| P50250 (PF3D7_0520900) | Adenosylhomocysteinase | 19.6 | 18.0 | 8 | 12.3 | 8.6 | 4 | 56.1 | 22.3 | 10 | Amino-acid biosynthesis |
| O96220 (PF3D7_0214000) | T-complex protein 1 subunit theta | 19.5 | 11.8 | 7 | 13.3 | 9.6 | 5 | 33.1 | 13.7 | 7 | - |
| Q8I2B1 (PF3D7_0102900) | Aspartate--tRNA ligase | 19.4 | 7.4 | 5 | 18.3 | 6.7 | 4 | 23.3 | 10.4 | 6 | - |
| C6KT23 (PF3D7_0618300) | 60S ribosomal protein L27a | 18.7 | 31.8 | 5 | 4.8 | 6.8 | 1 | 8.6 | 23.0 | 3 | Translation |
| Q8I577 (PF3D7_1234500) | Uncharacterized protein | 16.1 | 14.3 | 5 | 4.3 | 2.9 | 1 | 29.4 | 18.5 | 6 | - |
| Q8ILY9 (PF3D7_1410600) | Eukaryotic translation initiation factor 2 gamma subunit, putative | 16.1 | 9.0 | 4 | 16.4 | 12.0 | 5 | 19.1 | 10.5 | 3 | Translation |
| Q8I3R6 (PF3D7_0519400) | 40S ribosomal protein S24 | 15.4 | 31.6 | 5 | 5.7 | 18.1 | 2 | 24.0 | 15.8 | 4 | Translation |
| O77381 (PF3D7_0317600) | 40S ribosomal protein S11 | 12.8 | 26.7 | 4 | 32.5 | 37.9 | 6 | 46.9 | 41.0 | 7 | Translation |
| Q8I3R0 (PF3D7_0520000) | 40S ribosomal protein S9 | 11.6 | 16.9 | 4 | 13.3 | 18.5 | 4 | 42.4 | 28.0 | 6 | Translation |

| | | | | | | | | | | | |
|---|---|---|---|---|---|---|---|---|---|---|---|
| Q8IDR9 (PF3D7_1342000) | 40S ribosomal protein S6 | 10.6 | 16.3 | 3 | 4.9 | 10.8 | 2 | 38.9 | 19.9 | 6 | Translation |
| C6KT56 (PF3D7_0621900) | Signal recognition particle subunit SRP68 | 10.3 | 4.0 | 4 | 15.1 | 6.6 | 5 | 47.6 | 12.0 | 10 | Translation |
| Q8IIB4 (PF3D7_1124900) | 60S ribosomal protein L35 | 10.2 | 27.4 | 4 | 6.7 | 6.5 | 2 | 23.6 | 21.0 | 4 | Translation |
| Q8IBH7 (PF3D7_0728000) | Eukaryotic translation initiation factor 2 subunit alpha | 9.1 | 11.6 | 4 | 12.5 | 14.3 | 4 | 6.2 | 3.7 | 1 | Translation |
| Q8IKU0 (PF3D7_1453800) | Bifunctional glucose-6-phosphate 1-dehydrogenase/6-phosphogluconolactonase | 9.1 | 5.3 | 5 | 17.0 | 6.3 | 5 | 33.5 | 7.7 | 6 | - |
| Q8I5X9 (PF3D7_1206600) | DNA-directed RNA polymerase subunit beta | 8.2 | 2.6 | 4 | 42.9 | 9.4 | 12 | 54.4 | 9.2 | 11 | Transcription |
| Q8IJT9 (PF3D7_1010600) | Eukaryotic translation initiation factor 2 subunit beta | 7.6 | 12.6 | 3 | 5.2 | 5.4 | 1 | 15.6 | 17.6 | 4 | Translation |
| O77364 (PF3D7_0312800) | 60S ribosomal protein L26 | 7.1 | 7.1 | 1 | 3.6 | 7.1 | 1 | 7.8 | 7.1 | 1 | Translation |
| Q8I3X4 (PF3D7_0513300) | Purine nucleoside phosphorylase | 6.0 | 10.6 | 2 | 4.8 | 8.6 | 2 | 29.2 | 24.5 | 4 | - |
| Q8IJV5 (PF3D7_1009000) | Diphthine methyl ester synthase | 5.3 | 9.1 | 2 | 1.7 | 2.6 | 1 | 1.7 | 2.6 | 1 | - |
| Q8IKQ7 (PF3D7_1457300) | MA3 domain-containing protein | 4.8 | 2.8 | 2 | 2.3 | 2.5 | 2 | 25.8 | 5.8 | 3 | - |
| Q8ILW6 (PF3D7_1412800) | Glycylpeptide N-tetradecanoyltransferase | 4.8 | 3.9 | 1 | 4.8 | 4.9 | 2 | 0.0 | 1.7 | 1 | - |
| Q8ID96 (PF3D7_1361000) | Protein arginine N-methyltransferase | 4.7 | 1.9 | 1 | 2.1 | 1.9 | 1 | 11.0 | 1.9 | 1 | - |
| C0H5D7 (PF3D7_1326400) | Translation initiation factor eIF-2B subunit gamma | 4.0 | 1.9 | 1 | 7.7 | 5.7 | 3 | 8.2 | 3.4 | 2 | Translation |
| Q8ID33 (PF3D7_1367500) | NADH-cytochrome b5 reductase | 4.0 | 4.4 | 2 | 3.4 | 4.7 | 2 | 3.9 | 2.5 | 1 | - |
| C6KT96 (PF3D7_0626000) | Uncharacterized protein | 3.0 | 1.4 | 2 | 10.3 | 1.8 | 3 | 6.3 | 1.4 | 2 | - |
| Q8IEM3 (PF3D7_1309100) | 60S ribosomal protein L24 | 3.0 | 6.8 | 1 | 3.8 | 11.7 | 2 | 6.6 | 11.1 | 2 | Translation |
| Q8ILP6 (PF3D7_1420400) | Glycine--tRNA ligase, dATP synthetase | 2.8 | 1.6 | 1 | 2.1 | 0.9 | 1 | 0.0 | 0.8 | 1 | - |
| Q8IEP9 (PF3D7_1306600) | V-type proton ATPase subunit H | 2.7 | 2.9 | 1 | 1.6 | 1.5 | 1 | 5.1 | 3.1 | 1 | - |
| C0H4Y0 (PF3D7_0826500) | Ubiquitin conjugation factor E4 B | 2.3 | 0.9 | 1 | 7.4 | 1.4 | 2 | 6.2 | 1.9 | 2 | UPS |
| Q8I5F9 (PF3D7_1225800) | Ubiquitin-activating enzyme E1 | 2.2 | 1.3 | 1 | 2.8 | 2.2 | 2 | 3.9 | 3.7 | 2 | UPS |
| C0H5B3 (PF3D7_1313800) | Uncharacterized protein | 2.2 | 0.4 | 1 | 1.9 | 0.5 | 2 | 0.0 | 0.3 | 1 | - |
| Q8IIU6 (PF3D7_1105700) | tRNA-splicing ligase RtcB homolog | 2.1 | 3.6 | 2 | 0.0 | 2.0 | 1 | 7.0 | 5.1 | 2 | - |
| C0H4K8 (PF3D7_0705700) | 40S ribosomal protein S29 | 2.1 | 13.0 | 1 | 1.8 | 13.0 | 1 | 5.4 | 14.8 | 1 | Translation |
| O96185 (PFB0460c) | Uncharacterized protein | 2.0 | 0.4 | 1 | 2.8 | 0.5 | 1 | 15.7 | 2.5 | 4 | - |
| Q8IJV3 (PF3D7_1009200) | Small subunit rRNA synthesis-associated protein | 1.9 | 1.1 | 1 | 6.0 | 2.5 | 2 | 4.1 | 0.8 | 1 | - |
| O77323 (PF3D7_0308200) | T-complex protein 1 subunit eta | 1.9 | 3.2 | 1 | 9.6 | 5.2 | 2 | 21.5 | 8.5 | 4 | Protein folding |
| Q8IHY0 (PF3D7_1138500) | Protein phosphatase PPM2 | 1.9 | 1.0 | 1 | 4.6 | 1.0 | 1 | 3.2 | 1.5 | 1 | - |
| Q8IBB6 (PF3D7_0828500) | Translation initiation factor eIF-2B subunit alpha | 1.9 | 2.9 | 1 | 3.0 | 4.1 | 1 | 18.5 | 14.9 | 3 | Translation |
| C6KTB3 (PF3D7_0627700) | Transportin | 1.7 | 1.0 | 1 | 2.0 | 0.8 | 1 | 3.1 | 1.0 | 1 | Protein import into nucleus |
| C0H489 (PF3D7_0405100) | Protein transport protein Sec24B | 1.6 | 0.7 | 1 | 3.1 | 1.0 | 1 | 12.9 | 3.3 | 3 | Protein transport |

| | | | | | | | | | | | |
|---|---|---|---|---|---|---|---|---|---|---|---|
| Q8IHU0 (PF3D7_1142500) | 60S ribosomal protein L28 | 0.0 | 15.8 | 2 | 8.4 | 22.8 | 3 | 16.5 | 15.8 | 2 | Translation |
| Q8IBI3 (PF3D7_0727400) | Proteasome subunit alpha type | 0.0 | 3.9 | 1 | 2.0 | 3.1 | 1 | 8.3 | 6.3 | 1 | UPS |
| C6KTE1 (PF3D7_0630600) | DUF4205 domain-containing protein | 0.0 | 2.3 | 2 | 7.8 | 2.8 | 2 | 18.5 | 7.2 | 6 | UPS |
| Q8I5E0 (PF3D7_1227800) | Histone S-adenosyl methyltransferase | 0.0 | 0.9 | 1 | 5.3 | 1.8 | 2 | 2.4 | 1.9 | 1 | - |

### Figures and legends

```
PfDDI1   MVF----ITISDDNNIITSLDVHEDTEIWTIT--NIIENDFSLNMNINELTYNGNAV--D 52
ScDDI1   MDLT---ISNELTGEIYGPIEVSEDMALTDL--IALLQADCGFDKTKHDLYYNMDILDSN 55
HsDDI2   MLLTVYCVRRDLS-EVTFSLQVDADFELHNFR--ALCELESGIPAAESQIVYAERPLT-D 56
LmDDI1   ----------------------MVQLTINNARGVTLCRVSLPANATVQQLLL-------- 30

PfDDI1   KFDTIKKLNIKEGDLLFVRKKISADIMNDNVNNMSALNN--------------------- 91
ScDDI1   RTQSLKELGLKTDDLLLIRGKISNSIQ----------TDAAT------------------ 87
HsDDI2   NHRSLASYGLKDGDVVILRQKENADPRPPVQFPNLPRIDFSSIAVPGTSSPRQRQPPGTQ 116
LmDDI1   -------------------QLTVA--KPELRQAQAIRNDVRHV--------THRLTPAST 61

PfDDI1   --------ILSTNNNVGNIGNIGNNLNNENV--QNLLNNPAFKTLLDQFKVYQENEYIKK 141
ScDDI1   --LSDEAFIEQFRQELLNN---------QMLRSQLILQIPGLNDLVN------------- 123
HsDDI2   QSHSSPGEITSSPQ----------GLDNPALLRDMLLANPHELS---------------- 150
LmDDI1   TTTTTTSVVSSNAQTLLQAGLVGQGATAETLVVLMAADAPAAAS---------------- 105

PfDDI1   ESEILLEMKNDKSKMAVLKLQDEPLYNAIFSQNLEEIKKIVKEKYETEKKEKEKEQQMYE 201
ScDDI1   -------------DPLLFRERLGPLI---LQR---RYGGYNTAM---N-PFG-IPQDEYT 159
HsDDI2   ----------------LLKERNPPLAEALLSGDLEKFSRVLVEQ---QQDRARREQERIR 191
LmDDI1   ----------------SAAAAAPSPTKAVAAQILDLFGCASA-----SPSAGVRSQASVV 144

PfDDI1   NALKNPLSEDSQKFIYENIYKNEINNNLALAQEHFPEAFGVVFMLYIPVEINKNTVHAFV 261
ScDDI1   RLMANPDDPDNKKRIAELLDQQAIDEQLRNAIEYTPEMFTQVPMLYINIEINNYPVKAFV 219
HsDDI2   LFSADPFDLEAQAKIEEDIRQQNIEENMTIAMEEAPESFGQVVMLYINCKVNGHPVKAFV 251
LmDDI1   PSTMDERQLELQRRIYAQIQQQQIDENLANALEYTPEAFAKVTMLYVPCTINQVLVKAFV 204

PfDDI1   DSGAQSSIMSKKCAQKCNILRLMDKRFTGIAKGVGTKTILGKIHMIDIKIGNYFYAVSLT 321
ScDDI1   DTGAQTTIMSTRLAKKTGLSRMIDKRFIGEARGVGTGKIIGRIHQAQVKIETQYIPCSFT 279
HsDDI2   DSGAQMTIMSQACAERCNIMRLVDRRWAGIAKGVGTQKIIGRVHLAQVQIEGDFLPCSFS 311
LmDDI1   DSGAQNSIMNKRTAERCGLMRLVDVRMRGVAVGVGRQEICGRIHMTPVNLAGMYIPFAFY 264

PfDDI1   IIEDYDIDFIFGLDLLKRHQCLIDFKQNALIIEDN--KIPFLSEKDVISISTQSIDIDAN 379
ScDDI1   VLD-TDIDVLIGLDMLKRHLACVDLKENVLRIAEV--ETSFLSEAEIPKSFQEGLPAPTS 336
HsDDI2   ILEEQPMDMLLGLDMLKRHQCSIDLKKNVLVIGTTGSQTTFLPEGELPECARLAYGAGRE 371
LmDDI1   VIEDQAMDLIIGLDQLKRHQMMIDLKHNCLTIDNI--NVPFLPENDLPALAALGDDENAM 322

PfDDI1   NDL--------------------------------------------------------- 382
ScDDI1   VTT---SSDKPLTPTKTSSTLPPQPGAVPALAPRTGMGPTPTGRSTAGATTATGRTFPEQ 393
HsDDI2   DVRPEEIADQELA-EA----LQKSAEDAERQK------------P--------------- 399
LmDDI1   HA--PRHQDPATTATTASNPAAPVLSEGERQA------------RIEGFMTVSGITDPTQ 368

PfDDI1   ------------------------------------ 382
ScDDI1   TIKQLMDLGFPRDAVVKALKQTNGNAEFAASLLFQ- 428
HsDDI2   ------------------------------------ 399
LmDDI1   AAEL--------------LEAADWNPNVAAALLFDT 390
```

**Fig. S1. Sequence alignment of DDI1 proteins.** The amino acid sequence of *P. falciparum* DDI1 (PfDDI1) was aligned with the sequences of structurally characterized orthologs from *S. cerevisiae* (ScDDI1), *H. sapience* (HsDDI2) and *L. major* (LmDDI1). The UBL domain is marked with a blue line, RVP domain is in blue font and UBA domain is marked with a red line. Conserved amino acid residues are highlighted in yellow and physico-chemically similar residues are highlighted in grey. The number at the end of each line indicates the position of the last amino acid residue in the respective protein. The predicted catalytic aspartate residue is in red font.

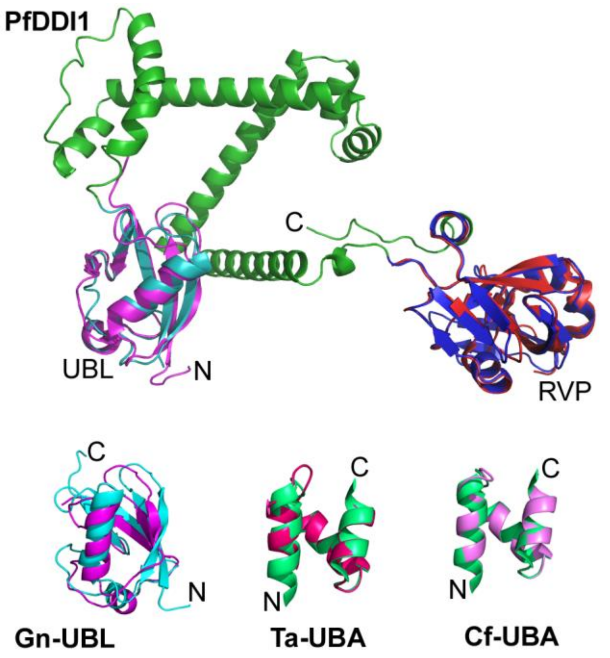


**Fig. S2. Domain structures of PfDDI1 and other apicomplexan DDI1 proteins.** The AlphaFold structure of PfDDI1 (AF-Q8IM03-F1) shows putative UBL (cyan) and RVP (blue)) domains that are superimposed on the reported structures of ScDDI1 UBL (magenta) and RVP (red) domains. The *Gregarina niphandrodes* UBL (Gn-UBL, 2-70), *Theileria annulata* UBA (Ta-UBA, 367-403) and *Cytauxzoon felis* UBA (Cf-UBA, 354-393) domains were predicted by homology modeling. The modelled structure of Gn-UBL (cyan) was superimposed on the reported structure of ScDDI1 UBL (magenta). The modelled structures of Ta-UBA (pink) and Cf-UBA (purple) were superimposed on the reported structure of *Schizosaccharomyces pombe* DDI1 UBA domain (green). N and C are for N- and C-termini, respectively.

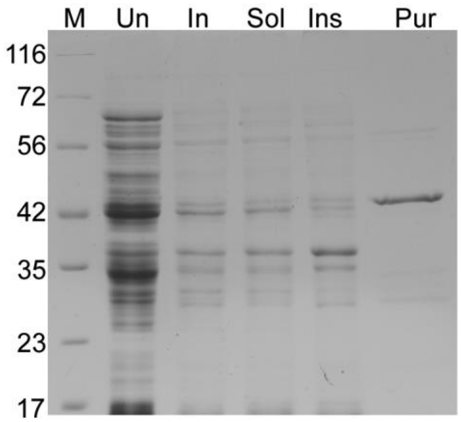


**Fig. S3. Production of recombinant PfDDI1.** The native PfDDI1 coding region was expressed as an N-terminal His-tagged protein using the pQE30 plasmid in *E. coli* M15(pREP4) cells. The recombinant protein was purified from IPTG-induced cells by Ni-NTA chromatography under denaturing conditions and refolded. The lysates of uninduced (Un) and induced (In) cells, soluble (Sol) and insoluble (Ins) fractions of induced cells, and refolded protein (Pur) were run on SDS-PAGE gel and visualized by coomassie staining. The sizes of protein markers (M) are indicated in kDa.

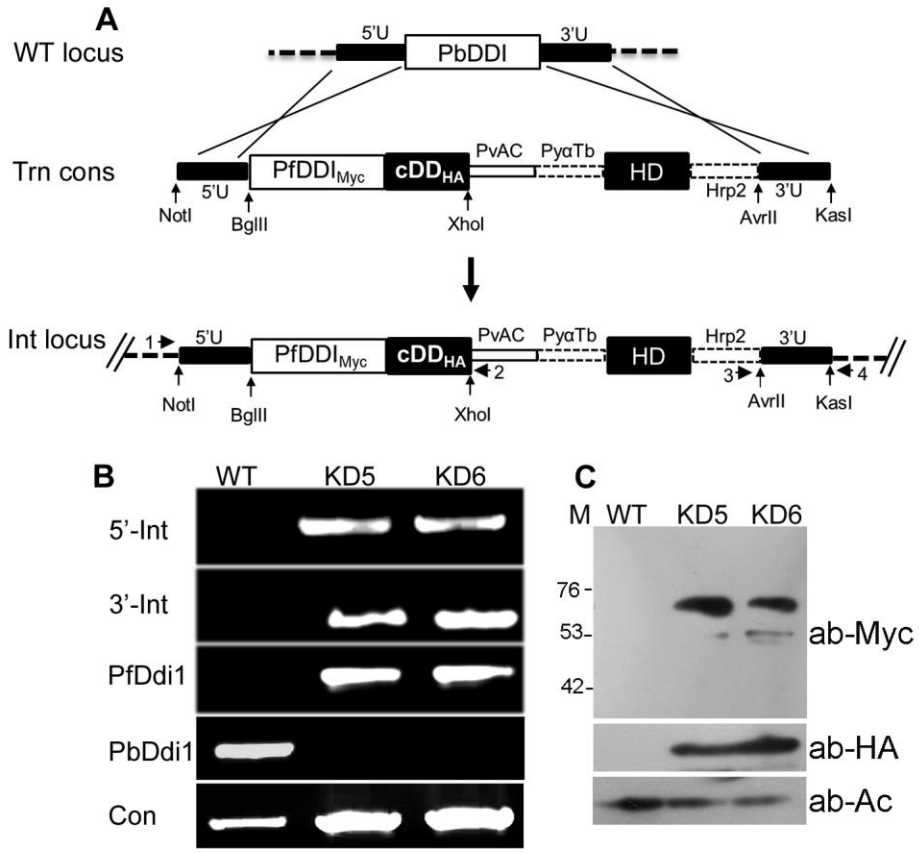


**Fig. S4. Generation of PbKD parasites.** The endogenous PbDDI1 coding region was replaced with PfDDI1$_{Myc}$/cDD$_{HA}$ coding sequence, and cloned parasites were assessed for the presence of knock-down locus by PCR and expression of PfDDI1$_{Myc}$/cDD$_{HA}$ protein by western blotting. **A.** The schematic represents integration of linear transfection construct (Trn cons) into the endogenous PbDDI1 locus (WT locus), resulting into the generation of integration locus (Int locus). Rectangular boxes represent the coding regions for PbDDI1 (PbDDI), PfDDI1 (PfDDI) cDD$_{HA}$ and hDHFR (HD). The flanking untranslated regions (5′U and 3′U), the location and orientation of primers (horizontal arrows), the restriction endonuclease sites (vertical arrows), and regulatory regions in the linear transfection construct (3′U of PvAC, 5′U of PyαTb, 3′U of PfHrp2) are indicated. Locus specific primers (5' integration, 5'-Int (1/2); 3' integration, 3'-Int (3/4); wild type locus, WTsp (1/4); proteasome α-6 gene as a control, Con) were used to differentiate the integration and wild type loci by PCR. **B.** The ethidium bromide stained agarose gel shows PCR products for the indicated regions from the genomic DNAs of wild type (WT) and knock-down

(KD5 and KD6) parasites. **C.** The western blot of WT and knock-down (KD5 and KD6) parasite lysates was probed with antibodies to Myc (ab-Myc), HA (ab-HA) or β-actin (ab-Ac).

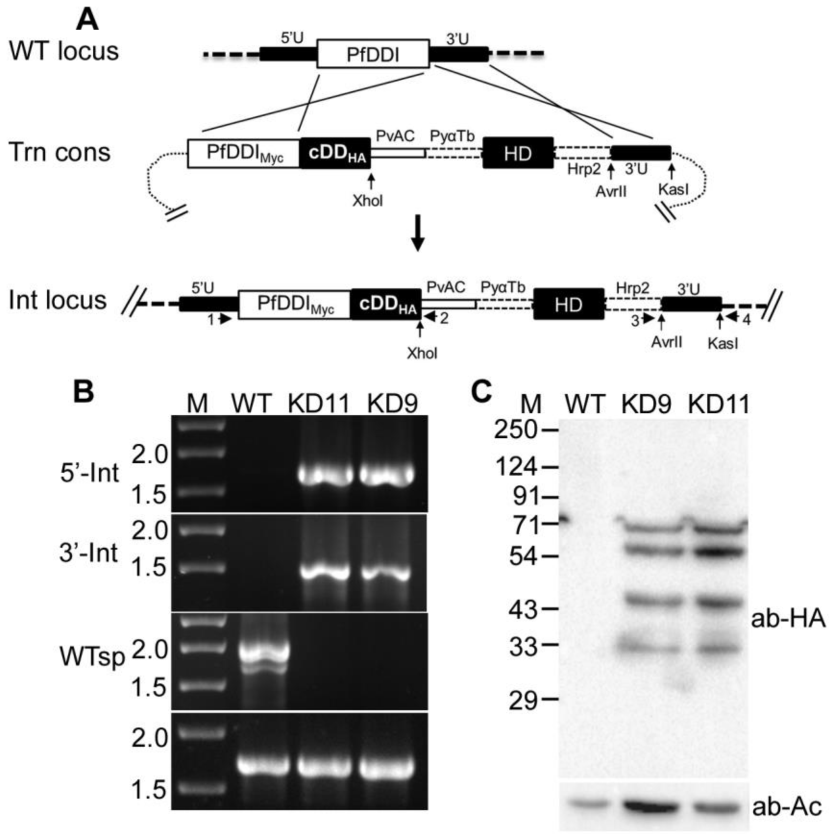


**Fig. S5. Generation of PfKD parasites.** The wild type PfDDI1 coding region was replaced with $PfDDI1_{Myc}/cDD_{HA}$ coding sequence, and the recombinant parasites were assessed for the presence of knock-down locus by PCR and expression of $PfDDI1_{Myc}/cDD_{HA}$ protein by western blotting. **A.** The schematic represents integration of transfection construct (Trn cons) into the endogenous PfDDI1 locus (WT locus) via double homologous crossover, resulting into the generation of integration locus (Int locus). Rectangular boxes represent the coding regions for PfDDI1 (PfDDI), $PfDDI1_{Myc}$ ($PfDDI_{Myc}$), $cDD_{HA}$ and hDHFR (HD). The flanking untranslated regions (5′U and 3′U), the location and orientation of primers (horizontal arrows), the restriction endonuclease sites (vertical arrows), and regulatory regions in the transfection construct (3′U of PvAC, 5′U of PyαTb, 3′U of PfHrp2) are indicated. **B.** Primers specific for different regions (5' integration, 5'-Int (1/2); 3' integration, 3'-Int (3/4); wild type locus, WTsp (1/4); vacuole membrane protein 1 gene as a control, Con) were used in PCR to differentiate the wild type and integration loci. PCR products for the indicated regions were amplified from the genomic DNAs of wild type (WT) and cloned knock-down (KD11 and KD9) parasites, and are shown in the ethidium bromide stained agarose gel. The lane M has DNA markers in kbps. **C.** The western blot of WT and knock-down (KD9 and KD11) parasite lysates was probed with antibodies to HA (ab-HA) or β-actin (ab-Ac).

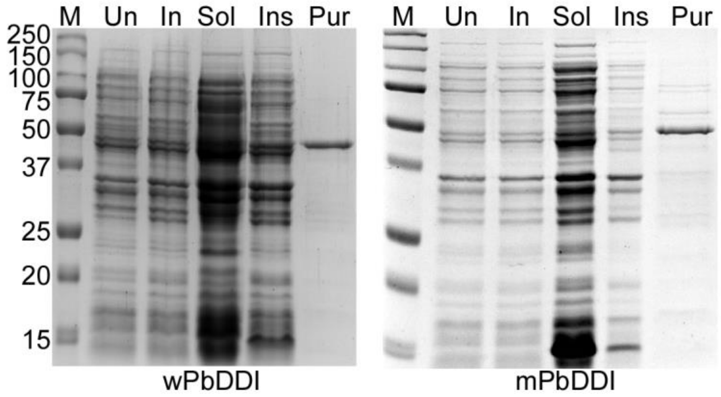


**Fig. S6. Production of recombinant PbDDI1.** Wild type *P. berghei* DDI1 (wPbDDI) and its catalytic mutant (mPbDDI) were expressed as N-terminal His-tagged proteins in *BL21-CodonPlus* (*DE3*) *E. coli* cells using the pRSET-A plasmid. The recombinant proteins were purified by Ni-NTA chromatography from IPTG-induced cells. The coomassie stained SDS PAGE gels show lysates of uninduced (Un) and induced (In) cells, soluble (Sol) and insoluble (Ins) fractions of induced cells, and Ni-NTA purified protein (Pur). The sizes of protein markers (M) are indicated in kDa.

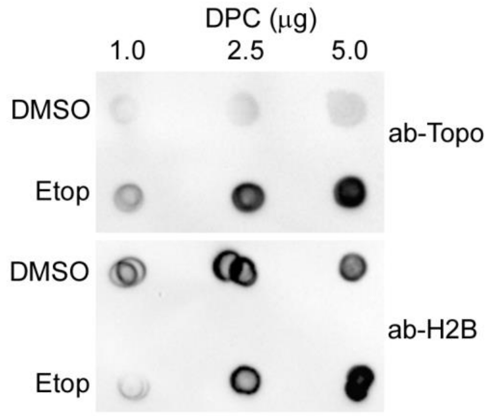


**Fig. S7. DPC preparation from HEK293T cells.** DPCs were isolated from HEK293T cells treated with DMSO or etoposide (Etop). The indicated amounts of DPCs were spotted on a nitrocellulose membrane, and the presence of topoisomerase II-DPC was probed using anti-topoisomerase II antibodies (ab-Topo) antibodies. Antibodies to histone 2B were used as a control to ensure the presence of chromatin.

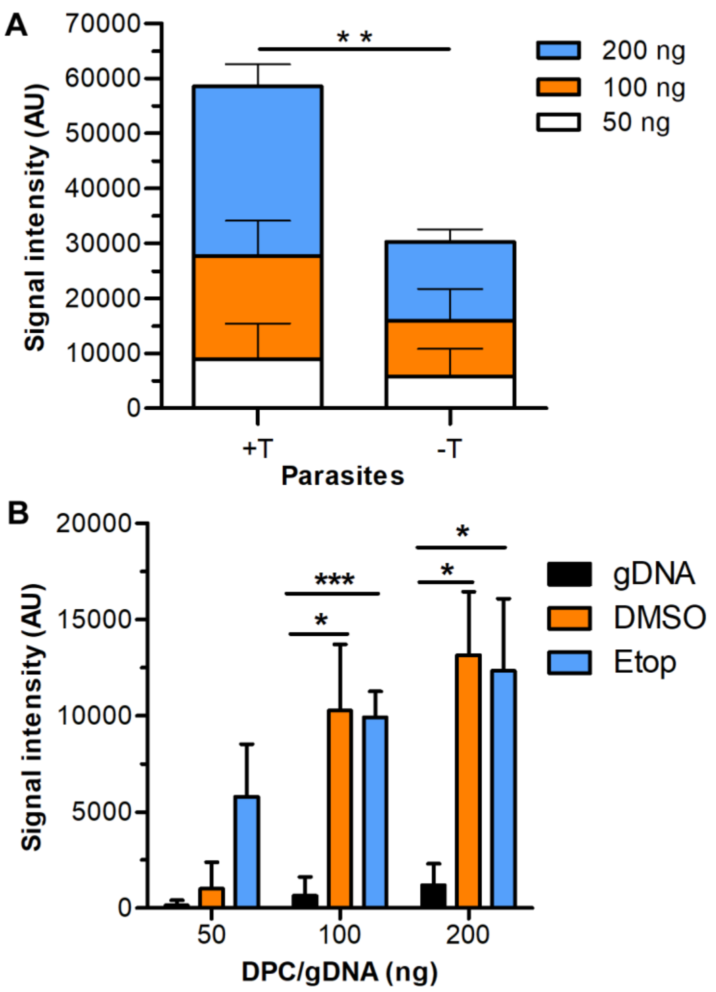


**Fig. S8. Association of PfDDI1 with DPCs. A.** The plot shows signal intensity in arbitrary units (AU) of DPC-associated PfDDI1 in Fig. 8B for PfKD parasites grown in the presence (+T) or absence (-T) of trimethoprim. The data is mean of three independent experiments with SD error bar. **B.** The plot shows signal intensity of recombinant PfDDI1$_{Myc/His}$ associated with the indicated amounts of DPCs or purified parasite gDNA (DPC/gDNA) in Fig. 8D. The signal intensity in Fig. 8C was subtracted from that of Fig. 8D for corresponding DPC/gDNA amount. The data is mean of three independent experiments with SD error bar. Statistical significance is indicated for data sets showing significant p values only (* is p<0.05-0.01, ** is p<0.01-0.001, *** is p<0.001).

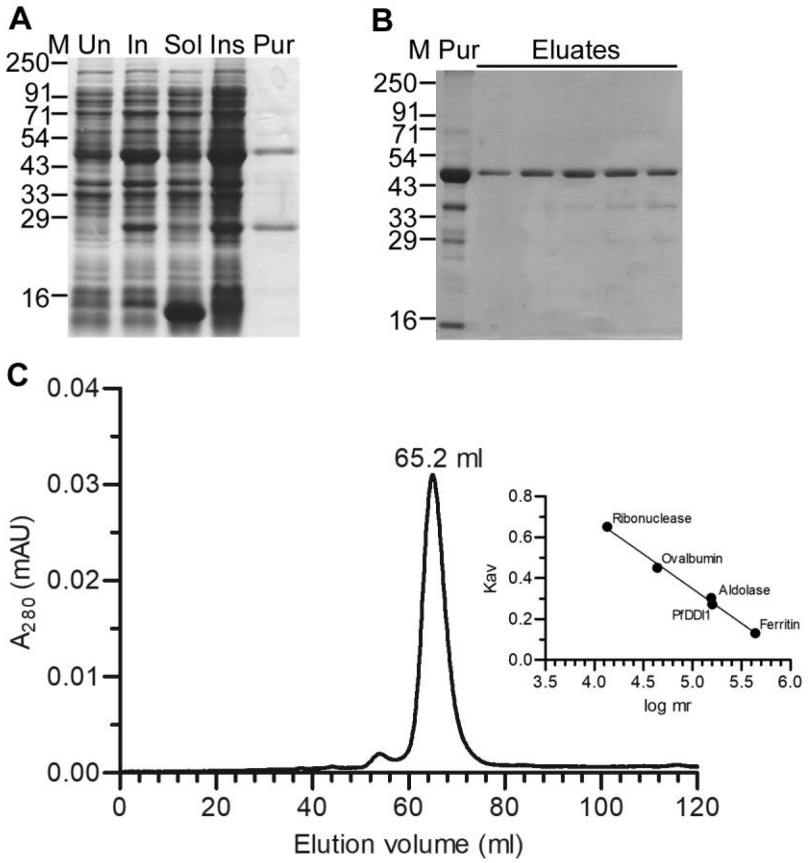


**Fig. S9. Production of recombinant PfDDI1.** The codon optimized synthetic PfDDI1$_{Myc}$ coding region was expressed as a C-terminal His-tagged protein (PfDDI1$_{Myc/His}$) in Rosetta-gami 2(DE3) cells, purified by Ni-NTA chromatography and then by gel filtration chromatography. **A.** The coomassie stained SDS-PAGE gel shows lysates of uninduced (Un) and induced (In) cells, soluble (Sol) and insoluble (Ins) fractions of the induced cells, and Ni-NTA purified protein (Pur). **B.** The Ni-NTA purified protein (Pur) was further purified by gel filtration chromatography, and eluates were analyzed by SDS-PAGE. The coomassie stained SDS-PAGE gel shows different eluates. **C.** The first two eluates were concentrated and run on the gel filtration column and elution volume was noted. Reference protein size markers shown in the inset were also run under identical conditions, elution volumes of the reference size markers were plotted against their sizes, and used to determine the size of $PfDDI_{Myc/His}$. The sizes of protein markers are in kDa (M) in A and B.

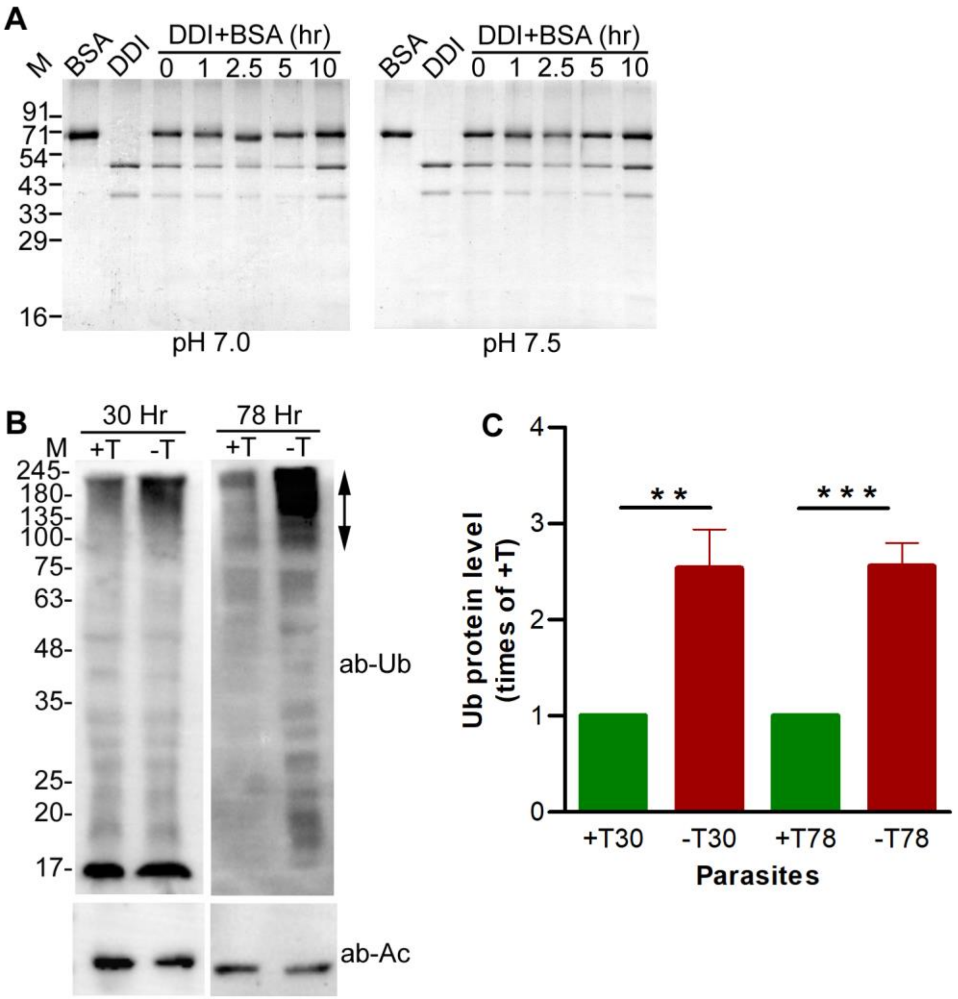


**Fig. S10. Protease assay and accumulation of ubiquitinated proteins. A.** The reaction sample containing BSA and recombinant $PfDDI1_{Myc/His}$ (DDI+BSA) was incubated at 37°C, equal aliquots were collected at the start and at the indicated time points, and the reaction was stopped with SDS-PAGE sample buffer. Control reaction samples contained BSA or DDI, incubated under identical conditions for 10 hours, and the reaction was stopped with SDS-PAGE sample buffer. The DDI+BSA aliquots, BSA and DDI1 samples were run in a 12% SDS PAGE and visualized by staining with coomassie blue. The sizes of proteins markers (M) are in kDa. **B.** PfKD parasites were grown in the presence (+T) or absence (-T) of trimethoprim, harvested at 30 hr and 78 hr time points, and processed for the presence of ubiquitinated proteins by western blotting using anti-ubiquitin antibodies (ab-Ub). β-actin (ab-Ac) was used as a loading control. The arrow indicates ubiquitinated proteins. The sizes of proteins markers (M) are in kDa. **C.** The intensity of ubiquitinated protein signal in +T and -T PfKD parasite lysates in “B” was measured and plotted as Ub protein level (times of +T) on y-axis against the parasite lysates (parasites) on x-axis. The data is mean of three independent experiments with SD error bar. Statistical significance is indicated for data sets showing significant p values (* is $p<0.05-0.01$, ** is $p<0.01-0.001$, *** is $p<0.001$).

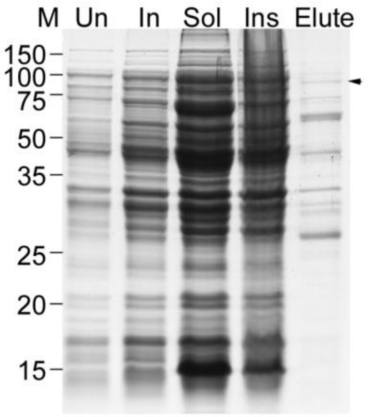


**Fig. S11. Production of recombinant PfPRP19.** The PfPRP19 coding sequence was expressed with an N-terminal GST-tag in *BL21*-(*DE3*) *E. coli* cells using the pGEX-6P-1 plasmid and purified from the IPTG-induced cells. The coomassie stained SDS-PAGE gel show lysates of uninduced (Un) and induced (In) cells, soluble (Sol) and insoluble (Ins) fractions of the induced cells, and the GST-PfPRP19 enriched eluate (Elute). The arrow head indicates predicted size of GST-PfPRP19 (~87.7 kDa) and the sizes of proteins markers are indicated in kDa (M).

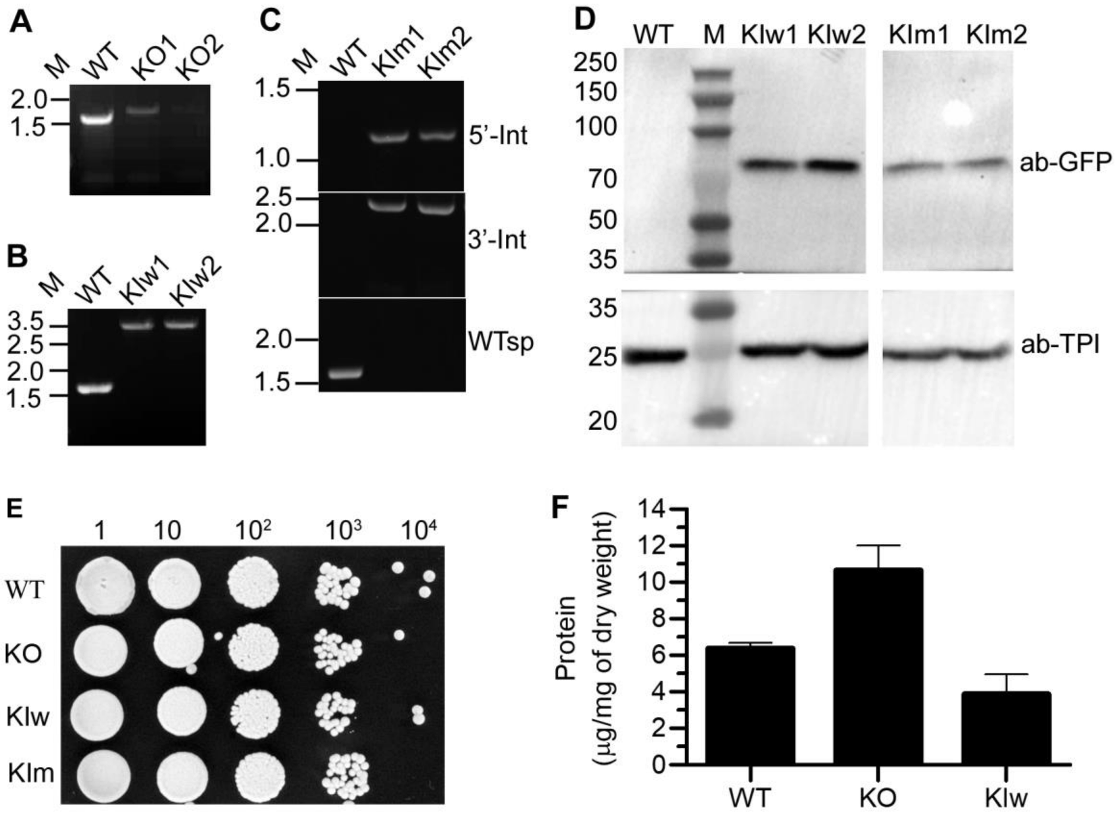


**Fig. S12. PfDDI1 reversed the hypersecretion phenotype of *S. cerevisiae* DDI1 mutant.** The endogenous ScDDI was knocked-out (KO) by replacing the ScDDI1 coding region with a kanamycin cassette. For generation of complemented strains, the ScDDI1 coding region was replaced with GFP-wPfDDI/kanamycin cassette coding for wild type (KIw) or catalytic mutant (KIm) of PfDDI1. The replacement of ScDDI1 was confirmed by PCR of the gDNAs of wild type (WT), knockout (KO1 and KO2) and knock-in (KIw1, KIw2, KIm1 and KIm2) strains using locus-specific primers. **A and B.** The ethidium bromide stained agarose gels contain PCR products for the wild type (WT), knock-out (KO) and wild type PfDDI1-complemented (KIw) loci. **C.** The ethidium bromide stained agarose gel contains PCR products corresponding to the 5' integration (5'-Int), 3' Integration (3'-Int) and wild type-specific (WTsp) loci of WT and mutant PfDDI1-complemented (KIw) strains. The positions of DNA markers are in kbp (M). **D. Western blot analysis of complemented strains.** The lysates of WT, KIw and KIm strains were processed for western blot using anti-GFP antibodies (ab-GFP), and antibodies to triose phosphate isomerase antibodies (ab-TPI) were used as a loading control. The numbers indicate sizes of protein markers in kDa (M). **E. Comparative growth assay.** The overnight cultures of WT, KO, KIw and KIm strains were normalized to $OD_{600}$ of 1.0, and the indicated dilutions of each culture were spotted on an YPD agar plate, which was incubated at 30°C till the colonies appeared. **F. Reversal of hypersecretion phenotype.** WT, KO and KIw strains were grown in SD medium for 48 hours at 30°C, the supernatants were dialyzed against PBS and the protein amounts were estimated. The cell pellets were dried and weighed. The plot shows amount of protein in the supernatant normalized to the dry weight of cell pellet. The data is mean of three independent experiments with SD error bar.
